# Salience-weighted agreement feature hierarchy modulates language comprehension

**DOI:** 10.1101/671834

**Authors:** R. Muralikrishnan, Ali Idrissi

**Affiliations:** Max Planck Institute for Empirical Aesthetics, Frankfurt, Germany; Qatar University, Doha, Qatar

**Keywords:** ERPs, Sentence Processing Subject-Verb Agreement Arabic, Agreement Feature Hierarchy

## Abstract

The brain establishes relations between elements of an unfolding sentence in order to incrementally build a representation of who is doing what based on various linguistic cues. Many languages systematically mark the verb and/or its arguments to imply the manner in which they are related. A common mechanism to this end is subject-verb agreement, whereby the marking on the verb covaries with one or more of the features such as person, number and gender of the subject argument in a sentence. The cross-linguistic variability of these features would suggest that they may modulate language comprehension differentially based on their relative weightings in a given language. To test this, we investigated the processing of subject-verb agreement in simple intransitive Arabic sentences in a visual event-related brain potential (ERP) study. Specifically, we examined the differences, if any, that ensue in the processing of person, number and gender features during online comprehension, employing sentences in which the verb either showed full agreement with the subject noun (singular or plural) or did not agree in one of the features. ERP responses were measured at the post-nominal verb. Results showed a biphasic negativity−late-positivity effect when the verb did not agree with its subject noun in one of the features, in line with similar findings from other languages. Crucially however, the biphasic effect for agreement violations was systematically graded based on the feature that was violated, which is a novel finding in view of results from other languages. Furthermore, this graded effect was qualitatively different for singular and plural subjects based on the differing salience of the features for each subject-type. These results suggest that agreement features, varying in their salience due to their language-specific weightings, differentially modulate language comprehension. We postulate a Salience-weighted Feature Hierarchy based on our findings and argue that this parsimoniously accounts for the diversity of existing cross-linguistic neurophysiological results on verb agreement processing.

## 1 Introduction

### 1.1 Background

An important evolutionary function of human language is to convey ecologically relevant information, such as the state of affairs of the entities in the immediate environment and event descriptions about who is doing what and to whom, so as to elicit an appropriate response in the given context. In order to comprehend the message, the brain thus has to be able to decipher these details incrementally by establishing the relations between the various elements in the unfolding utterance. Furthermore, these relationships must be constructed from the linguistic input even in the presence of intervening material separating the related elements.

Languages differ considerably in the mechanisms they employ in order to establish relations between an event being described in an utterance and the participant(s) of the event. One frequent device attested in many languages is to overtly express such relations on certain elements morphosyntactically (Nichols & Bickel, 2013), such that there is a ‘systematic covariance between a semantic or formal property of one element and a formal property of another (Steele, 1978, p. 610)’, commonly referred to as agreement. Agreement between a verb and its arguments may be based on some or all of the properties or features of the arguments concerned, such as person, number and gender of the nouns, collectively called agreement features (Moravcsik, 1978; Wechsler, 2009) or phi-features. Greenberg (1963) formulated a number of generalisations based on how these features show patterns of dependency and their frequency of occurrence relative to each other across a number of languages of the world. One such generalisation is the Feature Hierarchy shown in (1) below, which has been an important explanatory concept to account for the cross-linguistic diversity in how these features co-occur.

(1) Feature Hierarchy: Person > Number > Gender

Further sub-hierarchies of these features have been proposed in accounting for how they influence certain linguistic phenomena in a variety of typologically diverse languages(Silverstein, 1976; Shlonsky, 1989; Corbett, 2000a,b). The question of course is, whether such a hierarchy of agreement features identified on a cross-linguistic basis plays a role in online language comprehension. Indeed, such a systematic organisation of features is said to be a grammaticalised representation of fundamental cognitive categories (Harley & Ritter, 2002), with the relative position of a feature in the hierarchy reflecting the cognitive salience of the feature in relation to the other features, which in turn is said to correlate with dissociations in their online processing (De Vincenzi, 1999; Carminati, 2005; Acuña-Fariña, 2009). For instance, Carminati (2005, p. 263) proposes a Feature Strength Hypothesis mirroring the hierarchy in (1), suggesting that the cognitive significance of a feature should directly correlate with its relative hierarchical importance in language processing, whereby the more significant a feature, the better its disambiguating power is, and in turn the less processing costs it incurs, and vice versa. This would imply that a differential contribution, if any, of the agreement features must be observable in their respective neural processing correlates. Nevertheless, it is worth noting that the hierarchical importance of features posited based on their conceptual or cognitive significance need not necessarily preclude the possibility that the neural correlates of processing the agreement features in online language comprehension may show language-specific differences depending upon their relative salience in a given language, which in turn would be contingent upon the properties specific to the language concerned.

### 1.2 Previous studies

A number of studies have examined the processing of agreement features in different languages. In behavioural research, many studies employed a completion task involving sentences with agreement errors in the context of complex phrases, such as ‘*The key to the cabinets are on the table.*’ (see for instance Bock & Miller, 1991; Vigliocco, Butterworth, & Semenza, 1995; Hartsuiker, Antón-Méndez, & van Zee, 2001; Eberhard, Cutting, & Bock, 2005; Haskell, Thornton, & MacDonald, 2010, among others), whilst some involved a self-paced reading task (Wagers, Lau, & Phillips, 2009; Lago, Shalom, Sigman, Lau, & Phillips, 2015; Tucker, Idrissi, & Almeida, 2015), in some cases with eye-tracking (Pearlmutter, Garnsey, & Bock, 1999; Dillon, Mishler, Sloggett, & Phillips, 2013). An overarching finding from this line of research is that, agreement processes in sentence comprehension and production are complex, and involve factors that are ‘not only syntactic, not only semantic, and not only pragmatic, but all of these at once’ (Eberhard et al., 2005, p. 531). The cross-linguistic importance of agreement as a cue to sentence interpretation was demonstrated in a range of offline experiments conducted within the scope of the Competition Model (e.g., MacWhinney, Bates, & Kliegl, 1984; Bates & MacWhinney, 1989; Bates, 1999). These studies further showed that the strength of agreement as a cue to interpretation varies across different languages (for an overview, see Bates, McNew, Devescovi, & Wulfeck, 2001).

In order to gain insights into the neurocognitive mechanisms underlying the processing of agreement features, several previous studies have employed the ERP technique, which is particularly well-suited for studying language comprehension in real time thanks to its excellent temporal resolution. ERP studies investigating subject-verb agreement typically employ a violation paradigm, whereby the verb does not agree with its subject in the feature of interest. When ERPs measured at the anomalous verb are compared with those measured at the verb that shows correct agreement, this then is said to shed light on the neural correlates of processing the agreement feature under investigation.

Kutas & Hillyard (1983) reported one of the earliest ERP studies on subject-verb agreement in English and found that agreement violations elicited ERP effects that are different in scalp distribution to those elicited by semantic anomalies. Ever since, several ERP studies on the processing of agreement features in a number of languages have been reported (see Molinaro, Barber, & Carreiras, 2011, for a detailed review), the vast majority of which are on Indo-European languages. Results from these studies show that agreement violations of various types of dependencies (i.e., subject-verb, adjective-noun, article-noun etc.) generally elicit a left-anterior negativity (LAN) effect around 300 to 500 ms after the onset of the violation followed by a late-positivity (P600) effect in the 500 to 700 ms window and/or later. As Table 1 illustrates, this is true for the number feature in Dutch, English, Finnish, German, Italian and Spanish; for the gender feature in German, Italian and Spanish; for the person feature in German and Spanish. However, there are a few studies in which an N400 effect ensued instead of a LAN, or no negativity effect ensued at all. These include studies that examined number violations in Basque, English and Hindi; gender violations in Dutch, French, Hebrew, Hindi and Spanish; person violations in Basque and Spanish.

**Table 1:** A sample of findings previously reported in the literature for single-feature violations.

| Number violations |  |  |
| --- | --- | --- |
| Dutch | LAN–P600 | Hagoort, Brown, & Groothusen (1993); Kaan (2002) |
| English | LAN–P600 | Osterhout & Mobley (1995); Coulson, King, & Kutas (1998) |
| Finnish | LAN–P600 | Palolahti, Leino, Jokela, Kopra, & Paavilainen (2005) |
|  | LAN–P600 | Leinonen, Brattico, Järvenpää, & Krause (2008) |
| German | LAN–P600 | Roehm, Bornkessel, Haider, & Schlesewsky (2005) |
| Italian | LAN–P600 | Angrilli et al. (2002); De Vincenzi et al. (2003) |
| Spanish | LAN–P600 | Barber & Carreiras (2005); Mancini, Molinaro, Rizzi, & Carreiras (2011a) |
| Basque | P600 | Díaz, Sebastián-Gallés, Erdocia, Mueller, & Laka (2011) |
|  | N400–P600 | Zawiszwewski, Santestegán, & Laka (2016) |
|  | N400–P600 | Martinez de la Hidalga, Zawiszwewski, & Laka (2019) |
| English | P600 | Kaan, Harris, Gibson, & Holcomb (2000) |
| Hindi | P600 | Nevins, Dillon, Malhotra, & Phillips (2007) |
| Gender violations |  |  |
| German | LAN–P600 | Gunter, Friederici, & Schriefers (2000) |
| Italian | LAN–P600 | Molinaro, Vespignani, & Joba (2008) |
| Spanish | LAN–P600 | Barber & Carreiras (2005); Alemán Bañón & Rothman (2016) |
| Dutch | N400–P600 | Hagoort et al. (1993) |
| French | P600 | Frenck-Mestre, Osterhout, McLaughlin, & Foucart (2008) |
| Hebrew | N400–P600 | Deutsch & Bentin (2001, with an animacy interaction) |
| Hindi | P600 | Nevins et al. (2007) |
| Spanish | N400–P600 | Guajardo & Wicha (2014) |
| Person violations |  |  |
| German | LAN–P600 | Rossi, Gugler, Hahne, & Friederici (2005) |
| Spanish | LAN–P600 | Silva-Pereyra & Carreiras (2007) |
| Basque | N400–P600 | Zawiszwewski & Friederici (2009); Zawiszwewski et al. (2016) |
|  | N400–P600 | Martinez de la Hidalga et al. (2019) |
| Spanish | N400–P600 | Mancini et al. (2011a) |

Methodologically, it is of some import here to acknowledge the inherent difficulty in designing fully balanced experiments to investigate the processing of agreement features in any language. Discussing the merits of fully counter-balanced or symmetric designs, Steinhauer & Drury (2012) highlight the fact that it may be possible to employ symmetric designs in most studies examining, say, number violations, whereas it may not be viable to the same degree to employ such designs in studies investigating other kinds of violations. For instance, none of the widely-studied languages allow for fully crossed separate feature violations for the three agreement features. As an example, an agreement violation in English such as ‘*\*The girl eat the cake*’ could be construed at once as a number violation (cf. ‘*They eat the cake.*’) as well a person violation (cf. ‘*I eat the cake.*’) depending upon whether it is the ‘*They*’ or the ‘*I*’ contrast that one considers. Similar syncretisms exist to varying degrees in the agreement systems of French, German or Dutch; Italian or Spanish do not express gender agreement on the verb; Basque verbs express gender in the second person alone and only in some paradigms; Russian verbs express person and number but not gender in the present tense, number and gender but not person in the past tense; and Chinese or Japanese do not mark the verb for agreement.

Perhaps as an alternative means to mitigate such intrinsic limitations, some studies have compared a pair of feature violations with each other or with a combined violation of more than one feature. In the absence of a viable alternative, these studies provide important insights into the processing of agreement, which may not otherwise be available. However, a necessary limitation of these studies may be that their findings may not necessarily generalise always for the processing of all three features in isolation in a given language. A handful of such combined violation studies on subject-verb agreement have reported a modulation of effects (i.e., a quantitative ERP difference) based on the violating feature(s). These include the study on Basque cited above, in which the late-positivity was larger for person and combined person-number violations compared to number violations (Zawiswewski et al., 2016), the study on Hindi mentioned above, in which the late-positivity was found to be larger for combined person-gender violations versus number or gender or combined number-gender violations (Nevins et al., 2007), a study on Spanish that reported a larger late-positivity for combined person-number violations versus person or number violations (Silva-Pereyra & Carreiras, 2007), with the latter concluding that these results do not support the Feature Strength Hypothesis mentioned earlier. By contrast, in another Spanish study, Mancini et al. (2011a) reported a different distribution of the late-positivity in addition to a qualitative difference in the negativities, namely a LAN effect for the number feature and an N400 effect for the person feature.

To briefly summarise findings from ERP studies on agreement, whilst the LAN-P600 biphasic ERP effect is more common for agreement violations of many types, it is by no means universal, especially for subject-verb agreement. An N400 is elicited instead of a LAN in many cases, and no negativity at all ensues in some cases. Thus, neurophysiological evidence or counter-evidence for a hierarchy of features based on differences in their salience is inconclusive, with some studies reporting processing differences between the agreement features and others countering this, sometimes within the same language (compare for instance Alemán Bañón & Rothman, 2016; Silva-Pereyra & Carreiras, 2007). Further, a crucial aspect to note in this regard is that, although each feature has been studied in detail in a number of languages, and in some cases combinations of features compared with each other, to our knowledge, none of the studies reported to date compared the processing of person, number and gender agreement systematically within a single experiment. However, such a comparison would be imperative in order to shed light on whether the hierarchy of features postulated based on differences in their cognitive and language-specific salience and distribution across languages has a neural equivalent to it.

Does the hierarchy of agreement features indeed have a neural equivalent to it? If so, what is the nature of such a dissociation in neural correlates: is it a quantitative ERP difference (i.e., graded amplitude modulation of effects), or a qualitative one (i.e., different ERP effects)? Do the specific syntactic properties of a language interact with such a dissociation? In other words, do language-specific weightings of the features matter? In the following section, we argue that Arabic provides an ideal testing ground for investigating this.

### 1.3 Arabic as a test case

In order to investigate whether the hierarchy of agreement features postulated based on differences in their cognitive salience has a neural equivalent to it, certain properties appear to be crucial in determining the language of choice. A first such property is that the language in question should be morphologically rich such that all three agreement features, namely person, number, and gender are overtly expressed on the verb, either suffixally or in a fused form. A second and related property that becomes relevant is that these features should be expressed independently of each other, i.e., they are not underspecified for a certain feature in most cases such that the violation of each feature can be studied without simultaneously violating other features. For instance, commonly studied morphologically rich languages such as German or Spanish underspecify gender in verb agreement. A third property that is desirable for present purposes is that the feature markings on the target (i.e., the agreeing element (Corbett, 1983); here, the verb) are not always a one-to-one reflection of the agreement features of the element agreed with (i.e., the controller; here, the subject noun) in the language in question. In other words, the realisation of one or more features on the target is not only based on the features of the controller, but sometimes dependent upon certain structural properties of an utterance, such that changes in that property give rise to a concomitant change in the feature marking on the verb despite the subject being identical in both instances.

Arabic is a case in point in this regard. A Semitic language with upwards of 290 million speakers (Simons & Fenning, 2017) in countries spread around a great part of West Asia and North Africa, Arabic exhibits a typical example of diglossia (Ferguson, 1959), whereby a standard written variety for formal purposes and a number of colloquially spoken dialects exist in parallel. Modern Standard Arabic (Ryding, 2005) is the written standard taught in schools and universities (and the variety used for the stimuli here). Nouns are lexically gendered as feminine or masculine even if they are inanimate, and can express three different numbers, namely singular, dual and plural. Arabic is ideally suited to investigate possible processing differences between the agreement features in the context of subject-verb agreement mainly due to the fact that all three agreement features are expressed on the verb (with a few exceptions: gender is not expressed in the first person), as suffixes in the perfective and circumfixes in the imperfective. For example, let us consider a verb such as *‘to write’*, which in Arabic would be the triconsonantal root *k-t-b*, with its default citation form being *kataba ‘he wrote’*. Restricting ourselves to the past tense (and therefore only suffixes), and only to singular and plural forms, the verbal paradigm for this root would look as follows: -

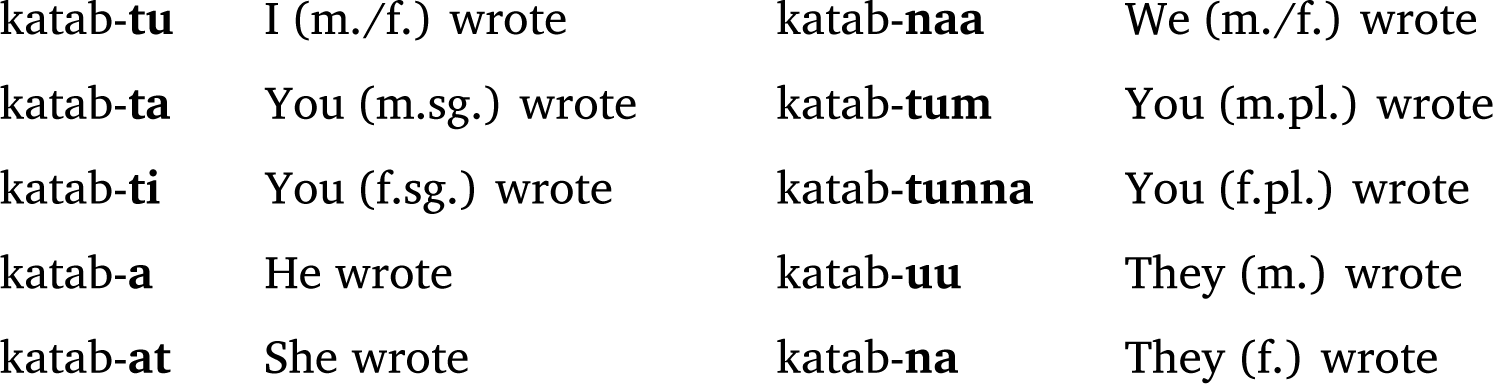

For an exhaustive introduction to Arabic verbal paradigms, the interested reader is referred to Ryding (2005), but for the present purposes, this example should suffice to illustrate the crucial point discussed earlier, namely that the processing of all three agreement features can each be independently examined in Arabic. A list of minimal pair examples is provided in the supplementary supporting information illustrating the properties of Arabic agreement that are most relevant for our present purposes.

### 1.4 Relevant properties of Arabic agreement

The Arabic agreement system is morphologically rich, relatively complex, and shows synchronic and diachronic variation between the standard and spoken varieties as well as amongst the dialects (Ferguson, 1997). Of particular relevance here is the word-order dependent asymmetry in subject-verb agreement in case of plural subject nouns (Aoun, Benmamoun, & Choueiri, 2010). The verb generally shows strict or full agreement if the subject precedes the verb (SV order, but see further), whereby it is obligatory for the verb to agree with the subject in person, number and gender, both in Standard Arabic as well as the dialects. By contrast, if the verb precedes the subject (VS order), the verb agrees with the subject in person and gender only, and expresses singular agreement for singular as well as plural nouns that are overtly present. However, if the plural subject were to be covert or dropped, the verb must show plural agreement. That is, number agreement is not only dependent upon whether the subject is singular or plural, but also contingent upon the relative order of the verb and the subject, as well as whether or not the subject is overt. In other words, under certain circumstances, an identical verb form is sometimes correct and sometimes incorrect based on the above mentioned factors that go beyond the number property of the subject noun per se. Thus, a simple feature-matching between the number property of the subject and the verb would not always lead to the correct interpretation.

A further aspect of Arabic agreement relevant for the present purposes is the interaction of the number feature with the humanness property of the subject noun (Ryding, 2005). If the subject is singular, the verb agrees with it in person, number and gender, regardless of whether the noun is human or nonhuman. If the subject is plural, by contrast, agreement depends upon the humanness property of the noun. For human plural nouns, the verb shows number (plural) and gender agreement, whereas for nonhuman plural nouns, the verb must show singular feminine agreement in Standard Arabic (Belnap & Shabaneh, 1992), a pattern that has been referred to as *deflected agreement* (Ferguson, 1997). Nevertheless, there are exceptions to this rule bothways; i.e., nouns such as ‘people’, ‘family’ and a few other collective nouns denoting groups of human referents as abstractions are treated as nonhuman plurals, and as such commonly require deflected agreement (Ryding, 2005). Similarly, even though the subject is actually a nonhuman plural noun, strict agreement may be preferred instead of deflected agreement for idiomatic or stylistic reasons (Ambros, 1977), and sociolinguistic, pragmatic or communicative strategic reasons (Ferguson & Barlow, 1988). To add to this complexity, dialects allow considerable variation in this regard (D’Anna, 2017; Belnap & Shabaneh, 1992; Ambros, 1977) such that ‘alternative patterns of agreement are acceptable without apparent change in meaning’ (Ferguson & Barlow, 1988, p. 16), with the differences in nuance between simple plurality versus individuation, distributive or enumerative plurality playing a role in this regard. This is perhaps not surprising, given that on a more general level, ‘[w]hile person and gender categories seem to have little effect on the meaning of a verb, […] number is somewhat different. The number of participants in a situation, whether agents or recipients of an action, can affect the situation profoundly.’(Bybee, 1985, p. 23).

In sum, it is apparent that the number feature plays a significant role in the Arabic agreement system, interacting in complex ways with properties that are structural (word-order) as well as at the syntax-semantics interface (humanness). That is, plurality of the controller (i.e., subject noun) is key, both for the word-order and overtness based agreement asymmetry as well as humanness based deflected agreement in Arabic. These characteristics of the Arabic agreement system allow investigating potential differences between the agreement features in language processing not possible in quite a comparable manner in many other widely-spoken languages. Furthermore, neurolinguistic investigations of diverse languages are crucial for gaining a broader understanding of the neural underpinnings of language (Bornkessel-Schlesewsky & Schlesewsky, 2016), and as such, neurophysiological studies on Arabic are an important contribution from a widely-spoken but extremely understudied language.

### 1.5 The present study

We report a visual ERP study here, in which we investigated the processing of subject-verb agreement in simple intransitive SV sentences in Arabic. In this experiment, ERP responses were measured at the post-nominal verb as participants read the stimuli in a rapid serial visual presentation. We specifically examined the differences, if any, that ensue in the processing of person, number and gender features during online comprehension, employing sentences in which the verb either showed full agreement with the subject noun (singular or plural) or did not agree in one of the features. The motivations to exclusively employ intransitive sentences for the purposes of the experiment are two-fold. First, there is evidence suggesting that verb valency or transitivity has a strong influence on incremental interpretation, and in processing agreement in particular (Burkhardt, Fanselow, & Schlesewsky, 2007). Second, in line with the principle of relational minimality that posits that the parser initially prefers to construct minimal structures (see Bornkessel & Schlesewsky, 2006, among others), there is evidence that the initial argument is preferentially interpreted as the subject of an intransitive rather than a transitive structure (the so-called *subject-preference*) from a variety of languages, including German (Schriefers, Friederici, & Kuhn, 1995), Turkish (Demiral, Schlesewsky, & Bornkessel-Schlesewsky, 2008) and Mandarin Chinese (Wang, Schlesewsky, Bickel, & Bornkessel-Schlesewsky, 2009) among others.

Furthermore, given the significance of the number feature in the Arabic agreement system, we hypothesised that processing differences may ensue between singular and plural agreement. Therefore, the subject number (i.e., singular versus plural) constituted an experimental factor in the design and analysis of our study (see Section 2.2 for further details).

## 2 Methods

### 2.1 Participants

Thirty-four persons, most of them students at the United Arab Emirates University in Al Ain, participated in the experiment after giving informed consent, and received monetary compensation for their participation. All participants were male^1^, and were right-handed native Arabic speakers, with normal or corrected-to-normal vision and normal hearing. Three further participants had to be excluded from the final data analyses on the basis of excessive EEG artefacts and/or too many errors in the behavioural control task. Since Arabic is diglossic (see Section 1.3), we collected information about the native dialect of participants. The self-reported dialect of Arabic that participants spoke natively at home was as follows. Palestinian: 11; Syrian: 8; Jordanian: 5; Egyptian: 3; Iraqi and Emirati: 2 each; Lebanese, Moroccon and Sudanese: 1 each.

### 2.2 Experimental Design

We employed an experimental design constituting Arabic sentences in the SV order with singular and plural subject nouns. All critical sentences were of the form adverb - subject - verb - prepositional phrase. The adverb was always *‘yesterday’*, subject nouns were always animate and human, and the verbs were in the simple past tense. The verb in each sentence was either in correct agreement with its subject noun, or it did not agree in either person, or number, or gender with the subject (see Section 2.3 for details about the materials). Note that any violation always involved one feature only. This yielded a design consisting of eight critical conditions that differed based on two factors, namely, whether the subject-type (ST) was singular or plural; and whether the condition-type (CT) was acceptable or a person violation or a number violation or a gender violation. The motivation at the outset to define and manipulate subject-type as a primary experimental factor in the present study was crucially informed by the significant role that the number feature plays in the Arabic agreement system (see Section 1.4 above). Such a design would allow us to observe the ERP effects at the position of the verb when information about the subject noun is already fully available. Any differences in ERP effects at the verb would reflect the differences in the processing of the different agreement features. Importantly, any potential subject-type specific differentiation of effects in processing agreement would also become apparent.

Table 2 provides an overview of the factors and their levels, with the condition codes relevant to each level, as well as examples pertaining to each condition. Note that Arabic is written and read from right to left, and the example sentences in the table follow this convention. The condition labels that are abbreviated in the table are as follows: SACP: Singular subject, Acceptable agreement; SGEN: Singular subject, Gender agreement violation; SNUM: Singular subject, Number agreement violation; SPER: Singular subject, Person agreement violation; PACP: Plural subject, Acceptable agreement; PGEN: Plural subject, Gender agreement violation; PNUM: Plural subject, Number agreement violation; PPER: Plural subject, Person agreement violation.

**Table 2:** Critical factors and their individual levels, with condition codes and a complete set of corresponding example stimuli.

| F a c t o r | L e v e l | C o n d i t i o n s |
| --- | --- | --- |
| ST : Subject-Type | S : Singular noun | SACP, SGEN, SNUM, SPER have a human singular subject, half of them feminine and half masculine |
|  | P : Plural noun | PACP, PGEN, PNUM, PPER have a human plural subject, half of them feminine and half masculine |
| CT : Condition-Type | ACP : Acceptable | SACP, PACP have a verb that shows full agreement with its subject noun |
|  | GEN : Violation of Gender Agreement | SGEN, PGEN have a verb that violates gender agreement |
|  | NUM : Violation of Number Agreement | SNUM, PNUM have a verb that violates number agreement |
|  | PER : Violation of Person Agreement | SPER, PPER have a verb that violates person agreement |

SACP
بالأمس بالأمس بالأمس بالأمس الممرضة الممرضة الممرضة الممرضة حَضَرَتْ حَضَرَتْ حَضَرَتْ حَضَرَتْ إلى المكان إلى المكان إلى المكان إلى المكان ʔl-makaan ʔilaa ʔl-makaan ʔilaa [DEF-place] [to] [come]<sub>Past-3sg.F</sub> [DEF-nurse]<sub>3sg.F</sub> [yesterday]
\* SGEN
بالأمس بالأمس بالأمس بالأمس الممرضة الممرضة الممرضة الممرضة حَضَرَ حَضَرَ حَضَرَ حَضَرَ إلى المكان إلى المكان إلى المكان إلى المكان ʔl-makaan ʔilaa ʔl-makaan ʔilaa [DEF-place] [to] [come]<sub>Past-3sg.M</sub> [DEF-nurse]<sub>3sg.F</sub> [yesterday]
\* SNUM
بالأمس بالأمس بالأمس بالأمس الممرضة الممرضة الممرضة الممرضة حَضَرْنَ حَضَرْنَ حَضَرْنَ حَضَرْنَ إلى المكان إلى المكان إلى المكان إلى المكان ʔl-makaan ʔilaa ʔl-makaan ʔilaa [DEF-place] [to] [come]<sub>Past-3pl.F</sub> [DEF-nurse]<sub>3sg.F</sub> [yesterday]
\* SPER
بالأمس بالأمس بالأمس بالأمس الممرضة الممرضة الممرضة الممرضة حَضَرْتُ حَضَرْتُ حَضَرْتُ حَضَرْتُ إلى المكان إلى المكان إلى المكان إلى المكان ʔl-makaan ʔilaa ʔl-makaan ʔilaa [DEF-place] [to] [come]<sub>Past-1sg</sub> [DEF-nurse]<sub>3sg.F</sub> [yesterday]
Singular Conditions. Right to left Arabic text with corresponding glosses. Intended meaning: ‘Yesterday the female nurse came to the place.’ A \* before condition labels indicates ungrammaticality.

### 2.3 Materials

As a first step, 120 intransitive verbs were used to construct acceptable sentences with singular nouns, half of which were masculine and the other half feminine. The plural acceptable sentences were then generated from these such that the subject nouns and verbs now showed plurality. Finally, three violation conditions were generated from each of the 240 acceptable sentences such that the agreement marking on the verb either violated person or number or gender in each instance (i.e., never was a combination of these features violated). This resulted in a total of 120 sets of sentences in eight critical conditions, thus 960 critical sentences. As Steinhauer & Drury (2012) have succinctly pointed out, it is crucial in ERP studies involving violations of the sort we have employed here for the experimental design to be fully symmetrical in order to rule out baseline problems as well as confounds resulting from lexical and contextual differences. In recognition of this important fact, our critical conditions were fully counter-balanced, such that there were 60 masculine nouns and 60 feminine nouns used as subjects in our stimuli; each subject noun in our experiment occurred both in its singular and plural forms an equal number of times; similarly, for a given subject noun, the identical verb was used with agreement markings as appropriate for the manipulation. Furthermore, since all the 120 subject nouns occurred in all the 8 critical conditions, there was an equal number of masculine (60) and feminine (60) nouns in acceptable sentences, person violation sentences, number violation sentences and gender violation sentences.

As is apparent from the examples in Table 2, the critical verb (third word from the right in each sentence) expresses the person, number and gender features according to the experimental manipulation involved, such that the affixes on the verb are different between the conditions. A comparable experimental manipulation is not viable in many of the widely-studied languages (see Section 1.2 above). Crucially however, the material preceding the critical position of interest within each subject-type is always identical for a given verb in all four conditions (acceptable, and the three violations). That is to say, for a given verb and subject-type, the context in which the verb is processed is identical, and in view of the counter-balancing of the materials in our experiment, this symmetric design would enable us to ‘avoid virtually all problems that are due to both lexical and contextual differences’ (Steinhauer & Drury, 2012, p. 141). In other words, any effect of lexical frequency and morphological complexity of the verb would apply to all conditions identically equally, and therefore does not confound potential language-related effects of interest such as N400, LAN and P600 observed at the position of the verb. Physical differences, such as the length of the plural verb forms versus their singular counterparts may engender amplitude differences in the P200 time-window, but should not be a cause for concern as far as the later effects of interest are concerned.

Fillers were constructed to ensure that the sentence structure is not predictable based on the first word alone and that the overall number of acceptable versus violation sentences is counterbalanced. Thus there were sentences involving subject and object relative clauses, verb-initial orders and a few semantic anomalies. The 960 target sentences were distributed into five unique sets according to a latin square, such that each list contained 36 sentences per critical condition. Of the resulting 288 critical sentences in a given list, 72 were acceptable and the rest were violations. Fillers were interspersed with these lists such that, overall, each stimulus list ended up with an equal number of acceptable and violation sentences, with an equal number of sentences with a masculine or feminine, singular or plural noun. This resulted in a total of 600 sentences per stimulus list (288 critical sentences and 312 fillers). These were each pseudo-randomised to obtain five stimulus lists, one of which was used for every participant. The presentation of the randomised lists was counterbalanced across participants.

### 2.4 Tasks

Given the use of a violation paradigm, an acceptability judgement task followed the presentation of each stimulus sentence, which required ‘yes’ or ‘no’ as answers. In addition, in order to ensure that participants would process the sentences attentively, a probe word detection task followed the acceptability judgement task. The probe task was constructed in such a way that an equal number of trials required ‘yes’ or ‘no’ as answers. If the probe word was one of the words that occurred in the preceding stimulus, this required a ‘yes’, whereas if it was novel, it required a ‘no’. Crucially, the word position from which the probe word was chosen was equiprobable across the experiment as well as within each condition, which meant that participants had to be very attentive throughout stimulus presentation so as to perform the task correctly.

### 2.5 Procedure

The methods and procedure employed in the experiment followed the guidelines of the Helsinki declaration. The experiment was performed in the EEG laboratory of the Department of Linguistics at the United Arab Emirates University in Al Ain. In accordance with local regulations, participants gave written informed consent, and received monetary compensation for their participation. All possible efforts were undertaken during and after the experiment to guarantee anonymity to the participants and protection of their personal data. Participants filled an Edinburgh-Handedness questionnaire in Arabic, and dominant right-handers alone were accepted for participation. They received printed instructions about the experiment and the task they had to perform.

After setting up the electrode cap, the participant moved to a sound-proof chamber, where they were seated on a comfortable chair and were requested to avoid abrupt and drastic movements, especially of the head. Then a resting state EEG was recorded for possible frequency-based EEG analyses later, where the participant had to sit still for two minutes with no specific task to perform. Two more minutes of resting state EEG was recorded, but now the participant had to close their eyes. After a short pause, the experimental session commenced, which consisted of a short practice followed by the actual experiment. Stimuli were presented using the Presentation software (www.neurobs.com) that recorded, among other things, the trial number, reaction time and the button responses. All physical parameters, such as the brightness and contrast settings of the monitor, distance between the chair and the monitor, and the lighting inside the chamber were maintained the same for all the participants.

The structure of each trial in the experiment was as follows. The flat-screen LCD monitor was clear before the trial commenced. A fixation asterisk was shown in the centre of the screen for 500 ms, after which the screen became blank for 100 ms. Then the rapid serial visual presentation of the stimulus sentence started. Each stimulus sentence was presented in four separate chunks, namely the adverb, the subject noun, the verb and the prepositional phrase. Since some prepositions form a single orthographic unit together with the noun that follows them, whereas others do not, the sentence-final prepositional phrase comprising of a preposition and a noun was always presented as a single chunk regardless of whether or not the preposition concerned formed a single orthographic unit together with the noun that follows it. Each of the first three chunks (words) appeared in the centre of the screen and remained for 600 ms, after which the screen became blank for 100 ms before the appearance of the next word. The prepositional phrases (containing two words) were presented for 750 ms. The word presentation rate we employed is in line with several existing studies on languages that are orthographically and/or or morphologically complex, such as Mandarin Chinese (Wang et al., 2009), Icelandic (Bornkessel-Schlesewsky, Roehm, Mailhammer, & Schlesewsky, 2020), Japanese (Wang & Schumacher, 2013) and Turkish (Demiral et al., 2008), among others, in order to provide a comfortable reading rate for participants. After the last chunk of the stimulus sentence was presented, the screen was blank for 500 ms. Following this, a pair of smileys appeared on screen, which prompted the participant to judge the acceptability of the sentence that just preceded. After a maximum of 2000 ms or after a button press, whichever was earlier, the screen became blank again for 500 ms. A time-out was registered when no button was pressed within 2000 ms. Then, a probe word appeared in the middle of the screen for a maximum of 2000 ms, within which the participant had to detect whether the word was present in the preceding stimulus sentence or not. When no button was pressed within 2000 ms, a time-out was registered. At the end of the trial, the screen became blank for 1500 ms (inter-stimulus interval) before the next trial started.

Before the actual experiment commenced, there was a short practice consisting of twelve trials, which helped participants to get used to the task and to feel comfortable about the pace of the trials and the blinking regime. For a given participant, none of the experimental stimuli occurred in their practice phase. The task was identical to that of the experiment phase. The EEG of the participants was not recorded in this phase.

In the main phase of the experiment, one of the five sets of materials as mentioned above was chosen to be presented in 12 blocks of 50 trials each. All critical trials and most filler trials consisted of 4 chunks (words/phrases), with a handful of filler trials consisting of 3/5/6 chunks. There were equal number of probe words that required ‘Yes’ or ‘No’ as answers in each block. Half the number of participants had the ‘Yes’ button on the right side and the other half had it on the left side so as to counterbalance for any right-dominance effects. The ‘Yes’ button being on the right or left was also counterbalanced across the stimuli sets. There was a short pause between blocks. Resting state EEG was again recored at the end of the experimental session. The duration of the overall experimental session including the practice and breaks but excluding the set-up time was about 1 hour and 10 minutes on an average.

### 2.6 EEG recording and pre-processing and statistical analysis

The EEG was recorded by means of 25 AgAgCl active electrodes fixed at the scalp by means of an elastic cap (Easycap GmbH, Herrsching, Germany). AFZ served as the ground electrode. Recordings were referenced to the electrode placed on the left mastoid online, but re-referenced offline to the average of the electrodes placed on left and right mastoids. The electrooculogram (EOG) was monitored by means of elecrodes placed at the outer canthus of each eye for the horizontal EOG and above and below the participant’s right eye for the vertical EOG. Electrode impedances were kept below appropriate levels such as to ensure a good quality signal with minimal noise. All EEG and EOG channels were amplified using a BrainAmp amplifier (Brain Products GmbH, Gilching, Germany) and recorded with a digitisation rate of 250 Hz. The EEG data thus collected was pre-processed for further analysis using a 0.3−20 Hz bandpass filter offline in order to remove slow signal drifts that might lead to stimulus-independent differences between conditions. Although baseline corrections are usually employed to exclusively remove differences in the critical epoch due to slow signal drifts, given that stimulus-induced ERP effects are not completely independent of EEG activity in the pre-stimulus interval (see Barry, Rushby, Smith, Clarke, & Croft, 2006; Makeig, 2002, for e.g.,), a distortion of effects in the critical ERP epochs may ensue due to transient signal differences between conditions in the baseline interval (see also Alday, 2019). The filter settings we employed effectively circumvents this issue without the need for a baseline correction^2^, and are sufficiently broad to include language-related ERP activity that is typically in the frequency range of about 0.5 to 5 Hz (Roehm, Winkler, Swaab, & Klimesch, 2002). Previous studies employing identical filter settings to ours report no significant differences in results between applying this bandpass filter versus applying baseline corrections (see Burkhardt et al., 2007; Choudhary, Schlesewsky, Roehm, & Bornkessel-Schlesewsky, 2009, for example). Furthermore, these filter settings are in line with several published ERP studies on language processing and has been employed in studies on languages as varied as Basque (Mancini, Massol, Duñabeitia, Carreiras, & Molinaro, 2019), German (Frenzel, Schlesewsky, & Bornkessel-Schlesewsky, 2015; Schumacher & Hung, 2012), Hindi (Choudhary et al., 2009), Japanese (Wolff, Schlesewsky, Hirotani, & Bornkessel-Schlesewsky, 2008), Mandarin Chinese (Wang et al., 2009), Spanish (Mancini, Molinaro, Rizzi, & Carreiras, 2011b) and Tamil (Muralikrishnan, Schlesewsky, & Bornkessel-Schlesewsky, 2015). The statistical analyses were performed on this data, but an 8.5 Hz low-pass filter was further applied to the grand-average ERPs for display purposes in order to achieve smoother ERP plots.

Artefacts in the continuous EEG data were rejected in two stages. First, epochs in which the signals from the EOG channels exceeded the threshold of 40 *μ*V were automatically rejected. Second, the thus resulting continuous EEG data was further visually inspected to remove eye blink artefacts and other non-stereotypical artefacts such as sudden head movements, chewing etc. that were not already rejected by the automatic rejection procedure. Trials in which the participants did not perform the probe detection task (see Section 2.4 for details about the tasks) were considered invalid trials and were excluded from the analysis at the outset; of the remaining valid trials (of about 34 per condition), an average of about 5 trials per condition had to be removed due to artefacts. Therefore the number of trials entering the averaging procedure did not differ significantly across conditions after artefact rejection, and data from about 28 to 29 trials remained in the analysis in each condition (Mean / standard deviation per condition: SACP = 29.44 / 3.08; SGEN = 29.14 / 3.27; SNUM = 28.61 / 2.70; SPER = 28.61 / 2.84; PACP = 29.58 / 2.81; PGEN = 28.97 / 3.44; PNUM = 28.85 / 3.11; PPER = 28.50 / 3.13).

ERPs were calculated for each participant from 200 ms before the onset of the verb until 1200 ms after onset (so −200 ms to 1200 ms). These were averaged across items per condition per participant before computing the grand-average ERPs across participants per condition. Repeated-measures analyses of variance (ANOVAs) were computed for the statistical analysis of the ERP data, involving the within-participants factors subject-type and condition-type for mean amplitude values per time-window per condition in 4 lateral Regions of Interest (ROIs) and 3 midline ROIs. The lateral ROIs (LATR in the statistical tables) were defined as follows: LA, comprised of the left-anterior electrodes F7, F3, FC5 and FC1; LP, comprised of the left-posterior electrodes P7, P3, CP5 and CP1; RA, comprised of the right-anterior electrodes F8, F4, FC6 and FC2; and RP, comprised of the right-posterior electrodes P8, P4, CP6 and CP2. The midline ROIs (MIDR in the statistical tables) were defined as follows: FC, comprised of the fronto-central electrodes FZ and FCZ; CP, comprised of the centro-pariteral electrodes CZ and CPZ; and PO, comprised of the parieto-occipital electrodes PZ and POZ.

The statistical analysis of the ERP data was carried out in a hierarchical manner in R (R Core Team, 2019). In order to examine the potential differences between the subject-types statistically (and not just by comparing results from separate analyses of the two subject-types), we included all 8 critical conditions (i.e., from both subject-types) in an omnibus ANOVA in each time-window of interest. Specifically, an ANOVA involving the within-participants factors subject-type and condition-type was computed for the lateral and midline regions separately in each time-window of interest, with the lateral analysis involving the lateral ROI factor (LATR) and the midline analysis involving the midline ROI factor (MIDR). In each analysis, as stated in Section 2.2, the primary factor was subject-type. This choice was motivated by the importance of the number property in the Arabic agreement system (see Section 1.4. To avoid excessive type 1 errors due to violations of sphericity, the correction of Huynh & Feldt (1970) was applied when the analysis involved factors with more than one degree of freedom in the numerator. Interactions that are at least marginally significant were resolved for the individual levels hierarchically. Factors with more than two levels, which resulted in a significant effect, were further resolved by comparing their individual levels pairwise. This procedure meant that the analysis was performed as follows: in each analysis, if there was a main effect of subject-type, this would show a general difference between the subject-types. If there was a main effect of condition-type, given that condition-type differed on 4 levels (ACP, GEN, NUM, PER), this effect would be resolved by pairwise comparisons of the 4 levels (shown as, say, ACP+NUM in the statistical tables to indicate a pairwise comparison of the acceptable condition versus the number violation condition). An effect resulting from such an individual pairwise comparison would be reported as significant only if it was still significant after applying the modified Bonferroni correction (Keppel, 1991). This resulted in a conservative significance scale for the pairwise comparisons, such that p values must be < 0.025 to be considered significant, and 0.025 < p < 0.04 already considered marginal, and p > 0.04 are not significant). If there was an interaction involving subject-type and condition-type, then the interaction will be resolved first by subject-type and then, if the effect of condition-type is significant in a given subject-type, that effect will be resolved by pairwise comparisons as described above. Further, given a resolvable effect was significant both with and without a lateral or midline ROI interaction in a certain analysis, then this suggests a topographically qualified difference, and therefore only the interaction involving the concerned ROI factor (LATR or MIDR) would be resolved further.

The statistical tables reporting ANOVA results follow these conventions: main effects and/or interactions that were at least marginally significant are reported; a factor or interaction following a bullet point means that factor or interaction was at least marginally significant; factorial resolutions are shown with an arrow followed by the factor and level; pairwise comparisons are shown using the format ACP+PER, where ACP and PER are the levels being compared pairwise; DF implies degrees of freedom; three stars beside p-values imply a significance of p <= 0.001; two stars imply p <= 0.01; a single star implies p <= 0.05; a single pale star implies a marginal (p <= 0.08) effect. For pairwise comparisons, the equivalent significance levels as per modified Bonferroni correction are p <= 0.0005, p <= 0.005, p <= 0.025 and p <= 0.04 respectively.

The tables use the following abbreviations for the factors and levels: ST: subject-type; ST = S: singular subjects; ST = P: plural subjects; CT: condition-type; ACP: acceptable conditions showing correct agreement; GEN: gender agreement violation; NUM: number agreement violation; PER: person agreement violation; LATR: lateral ROIs; LA: left-anterior; LP: left-posterior; RA: right-anterior; RP: right-posterior; MIDR: midline ROIs; FC: fronto-central; CP: centro-parietal; PO: parieto-occipital.

## 3 Results

### 3.1 Behavioural data

The mean acceptability ratings for the stimuli, as well as the probe detection accuracy for the critical conditions, shown in Table 3, were calculated using the behavioural data collected during the experiment. Only those trials in which the acceptability judgement task following each trial was performed (i.e., not timed out) were considered for the analysis. Further, the acceptability data presented here pertain only to those trials in which the participants performed the probe detection task correctly (see Section 2.4 for details about the tasks). Acceptability was highest for the conditions with no violations, whereas it was the lowest for conditions with number violations. Across conditions, acceptability was relatively slightly higher for conditions with a plural subject as compared to the corresponding conditions with a singular subject. The overall accuracy was very high across all conditions. Owing to the fact that the reaction time data are not time-locked to the critical manipulation in the stimulus sentences, they are not reported here (but are available from the corresponding author on request). Figure 1 shows raincloud plots (Allen, Poggiali, Whitaker, Marshall, & Kievit, 2019) of the behavioural acceptability judgements illustrating the differences in ratings between the subject-types. Panel A shows the by-participant variability of acceptability ratings, with the individual data points representing the mean by-participant acceptability of each subject-type and condition-type combination. Panel B shows the by-item variability of acceptability ratings, with the individual data points representing the mena by-item acceptability of each subject-type and condition-type combination.

**Figure 1:**
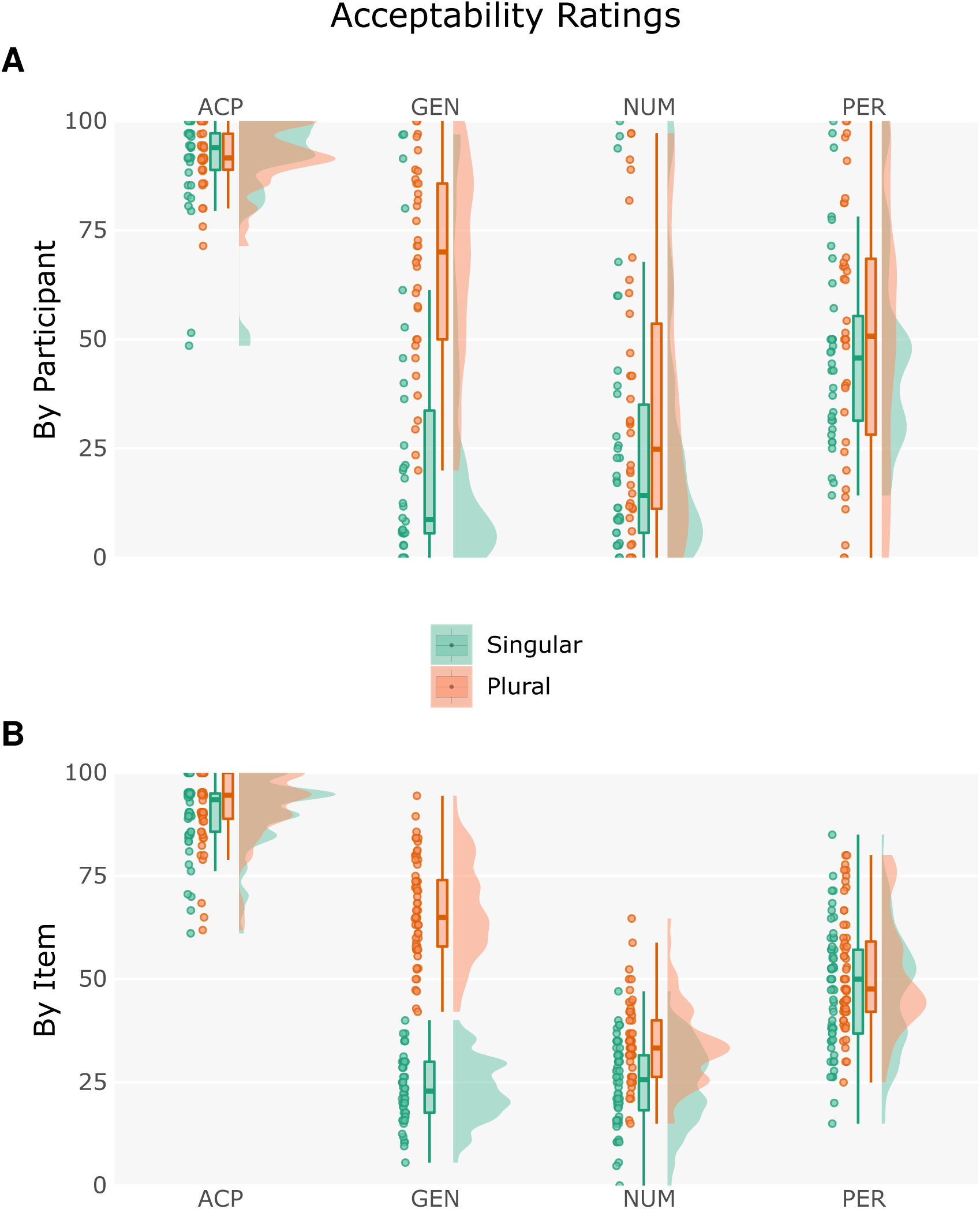
Raincloud plot of the acceptability ratings, showing by-participant variability (A) and by-item variability (B).

**Table 3:** Sentence acceptability ratings and probe detection accuracy

| Condition | Sentence Acceptability Rating % | SD | Probe Detection Accuracy % | SD |
| --- | --- | --- | --- | --- |
| SACP | 90.51 | 11.86 | 95.74 | 4.54 |
| SGEN | 23.68 | 29.67 | 94.48 | 4.10 |
| SNUM | 25.55 | 28.76 | 94.16 | 5.63 |
| SPER | 47.62 | 22.53 | 95.58 | 3.93 |
| PACP | 91.68 | 7.21 | 95.66 | 4.44 |
| PGEN | 67.00 | 22.99 | 94.42 | 4.89 |
| PNUM | 33.91 | 30.82 | 94.91 | 4.45 |
| PPER | 51.83 | 30.02 | 93.57 | 5.22 |

The statistical analysis of the behavioural data was performed by means of ANOVAs involving the within-subjects factors subject-type (ST) and condition-type (CT), and the random factors participants (F1) and items (F2). The statistical analysis was carried out in a hierarchical manner as described Section 2.6 above, with the exception that there was no ROI factor involved in the behavioural data analysis. Table 4 shows a summary of effects on the behavioural data collected during the experiment.

**Table 4:** ANOVA: Acceptability Data the verb.

| Effects on Acceptability | DF | F1:Participants | DF | F2:Items |
| --- | --- | --- | --- | --- |
| ♦ ST | 1,33 | 56.39 <.001 *** | 1,59 | 246.80 <.001 *** |
| ♦ CT | 3,99 | 105.60 <.001 *** | 3,177 | 642.50 <.001 *** |
| └CT = ACP+PER ♦ CT | 1,33 | 100.70 <.001 *** | 1,59 | 747.10 <.001 *** |
| └CT = ACP+NUM ♦ CT | 1,33 | 138.20 <.001 *** | 1,59 | 2503.00 <.001 *** |
| └CT = NUM+PER ♦ CT | 1,33 | 54.05 <.001 *** | 1,59 | 115.30 <.001 *** |
| └CT = ACP+GEN ♦ CT | 1,33 | 118.10 <.001 *** | 1,59 | 1298.00 <.001 *** |
| └CT = GEN+PER ♦ CT | 1,33 | 5.40 0.026 * | 1,59 | 8.99 0.003 ** |
| └CT = NUM+GEN ♦ CT | 1,33 | 48.57 <.001 *** | 1,59 | 130.90 <.001 *** |
| ♦ ST x CT | 3,99 | 30.96 <.001 *** | 3,177 | 142.60 <.001 *** |
| └ST = S ♦ CT | 3,99 | 122.40 <.001 *** | 3,177 | 613.30 <.001 *** |
| └CT = ACP+PER ♦ CT | 1,33 | 100.90 <.001 *** | 1,59 | 483.80 <.001 *** |
| └CT = ACP+NUM ♦ CT | 1,33 | 141.10 <.001 *** | 1,59 | 1512.00 <.001 *** |
| └CT = NUM+PER ♦ CT | 1,33 | 80.24 <.001 *** | 1,59 | 117.50 <.001 *** |
| └CT = ACP+GEN ♦ CT | 1,33 | 138.30 <.001 *** | 1,59 | 1967.00 <.001 *** |
| └CT = GEN+PER ♦ CT | 1,33 | 108.10 <.001 *** | 1,59 | 151.00 <.001 *** |
| └ST = P ♦ CT | 3,99 | 52.95 <.001 *** | 3,177 | 315.50 <.001 *** |
| └CT = ACP+PER ♦ CT | 1,33 | 66.45 <.001 *** | 1,59 | 487.80 <.001 *** |
| └CT = ACP+NUM ♦ CT | 1,33 | 102.80 <.001 *** | 1,59 | 1077.00 <.001 *** |
| └CT = NUM+PER ♦ CT | 1,33 | 11.68 0.001 ** | 1,59 | 53.87 <.001 *** |
| └CT = ACP+GEN ♦ CT | 1,33 | 44.88 <.001 *** | 1,59 | 188.70 <.001 *** |
| └CT = GEN+PER ♦ CT | 1,33 | 14.56 <.001 *** | 1,59 | 71.44 <.001 *** |
| └CT = NUM+GEN ♦ CT | 1,33 | 48.92 <.001 *** | 1,59 | 253.80 <.001 *** |

There were main effects of subject-type and condition-type on the acceptability in the analysis by participants as well as the analysis by items. Resolving the effect of condition-type by comparing the condition-types pairwise showed a significant simple effect of condition-type for all possible comparisons in both analyses, with the largest effect for the ACP + NUM comparison. The interaction subject-type x condition-type was significant in both the analyses, which when resolved for subject-type showed an effect of condition-type for both subject-types. This was further resolved by comparing the condition-types pairwise, which showed a simple effect of condition-type for all comparisons except NUM + GEN in both analyses when the subject was singular. When the subject was plural, there was a simple effect of condition-type for all comparisons in both analyses. There were no effects on the probe detection accuracy.

### 3.2 ERP data

The ERPs at the verb are shown in Figure 2 for the singular subject conditions, and in Figure 3 for the plural subject conditions. Visual inspection of the ERP data showed that the violation conditions mostly engendered a negativity followed by a late-positivity. This qualitative observation was further supported by a running t-test of significance on the ERP plots, as shown in Figure S4, which revealed at least two time-window clusters that should be significant. Along with these observations, considering ERP components that have been known to be relevant for processing agreement in previous studies, we chose two time-windows for analysis, namely 400−600ms and 700−850 ms. Figure 5 shows the topographic map of the ERPs at the position of the verb in the 400−600ms and 700−850 ms time-windows for the two subject-types in the three violation conditions, after the effects for the acceptable condition in each case has been subtracted. (See Section 2.6 for details regarding the statistical analysis).

**Figure 2:**
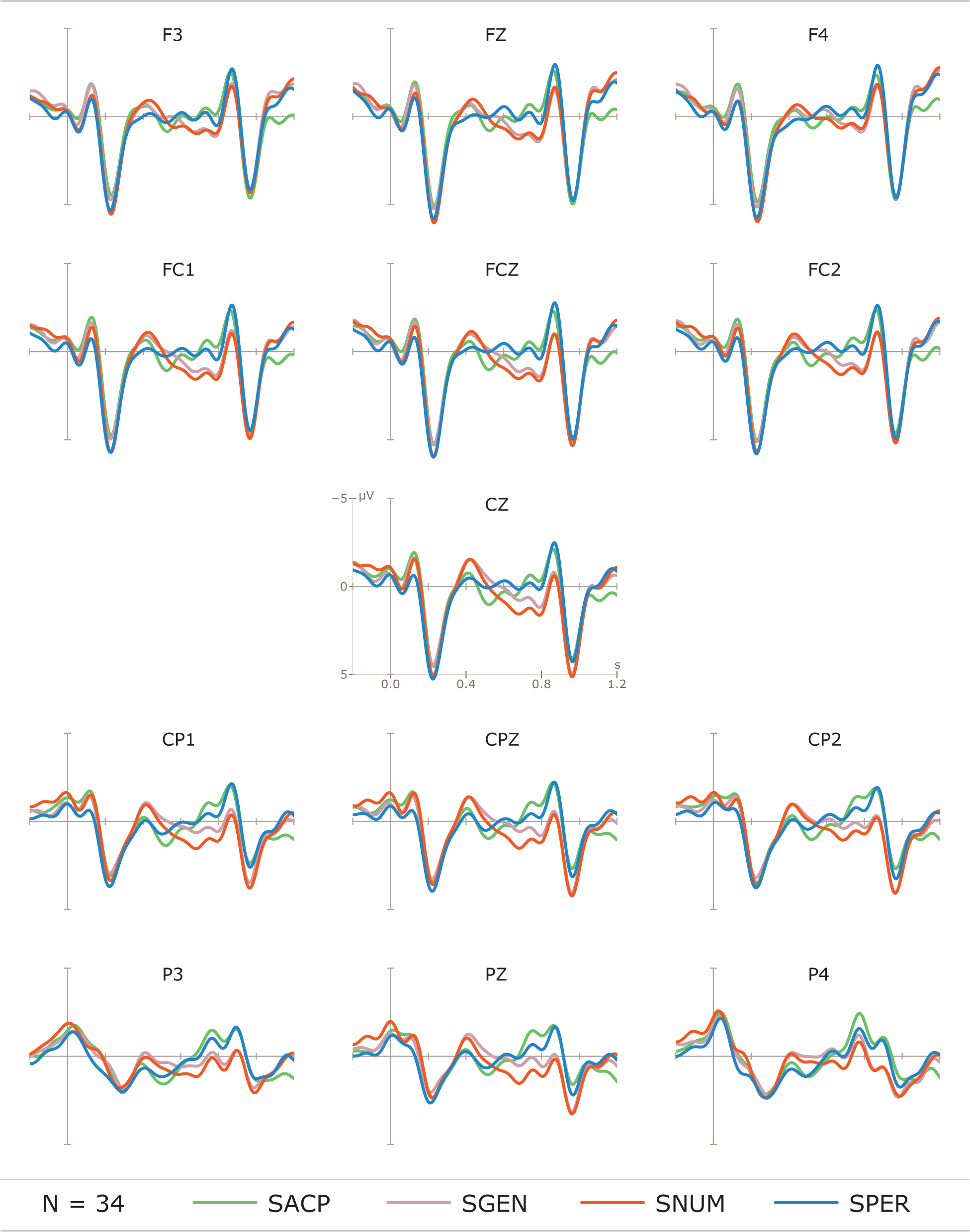
ERPs at the verb: Singular subject conditions.

**Figure 3:**
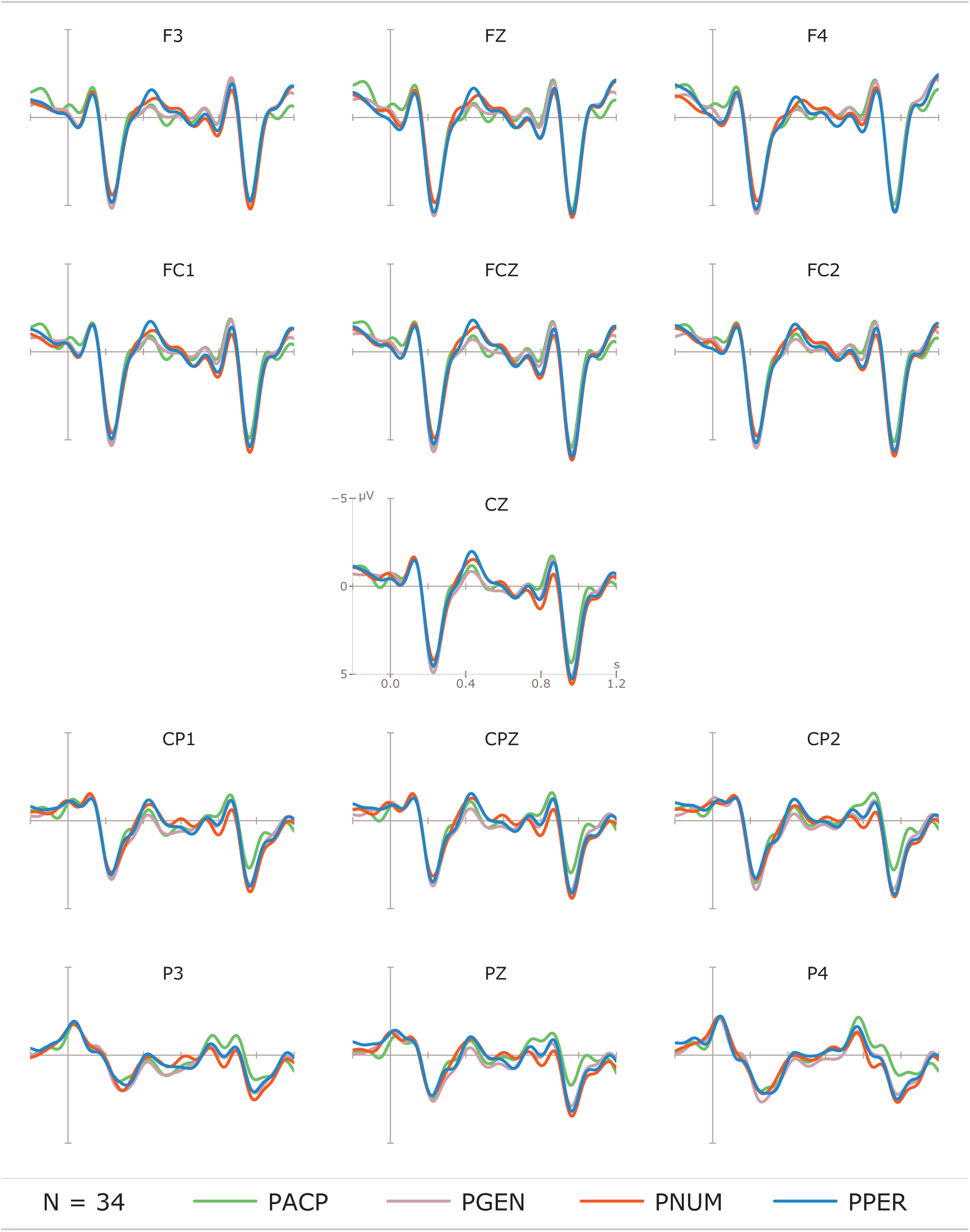
ERPs at the verb: Plural subject conditions.

**Figure 4:**
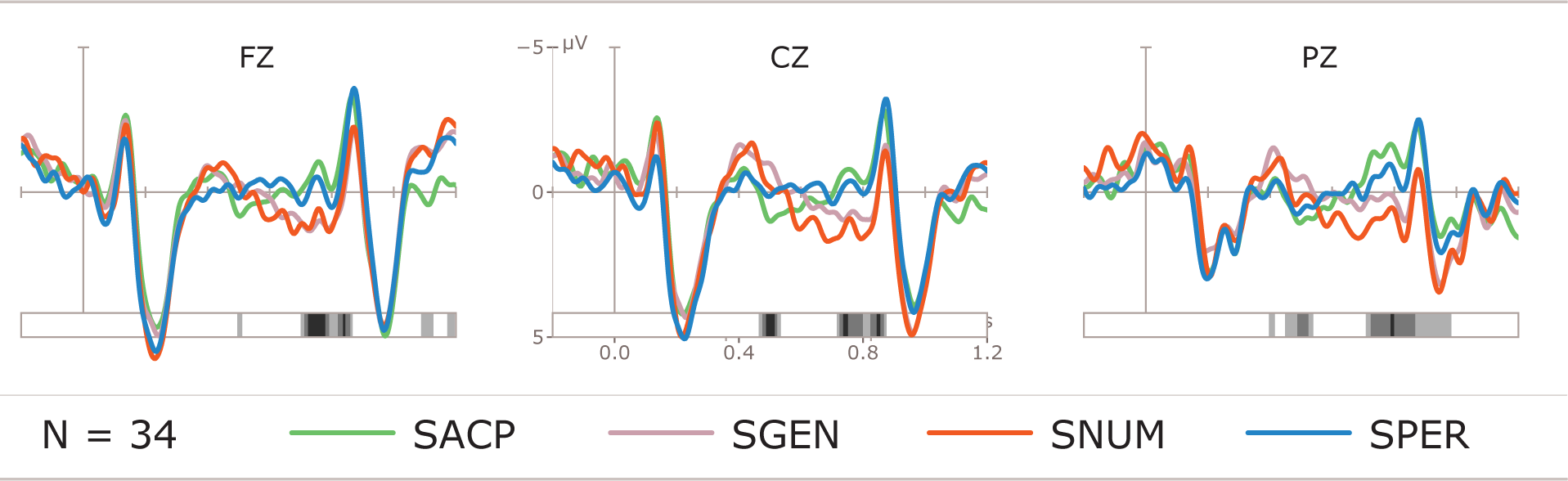
ERPs at the verb with running t-test to detect potentially significant time-windows: Singular subject conditions.

**Figure 5:**
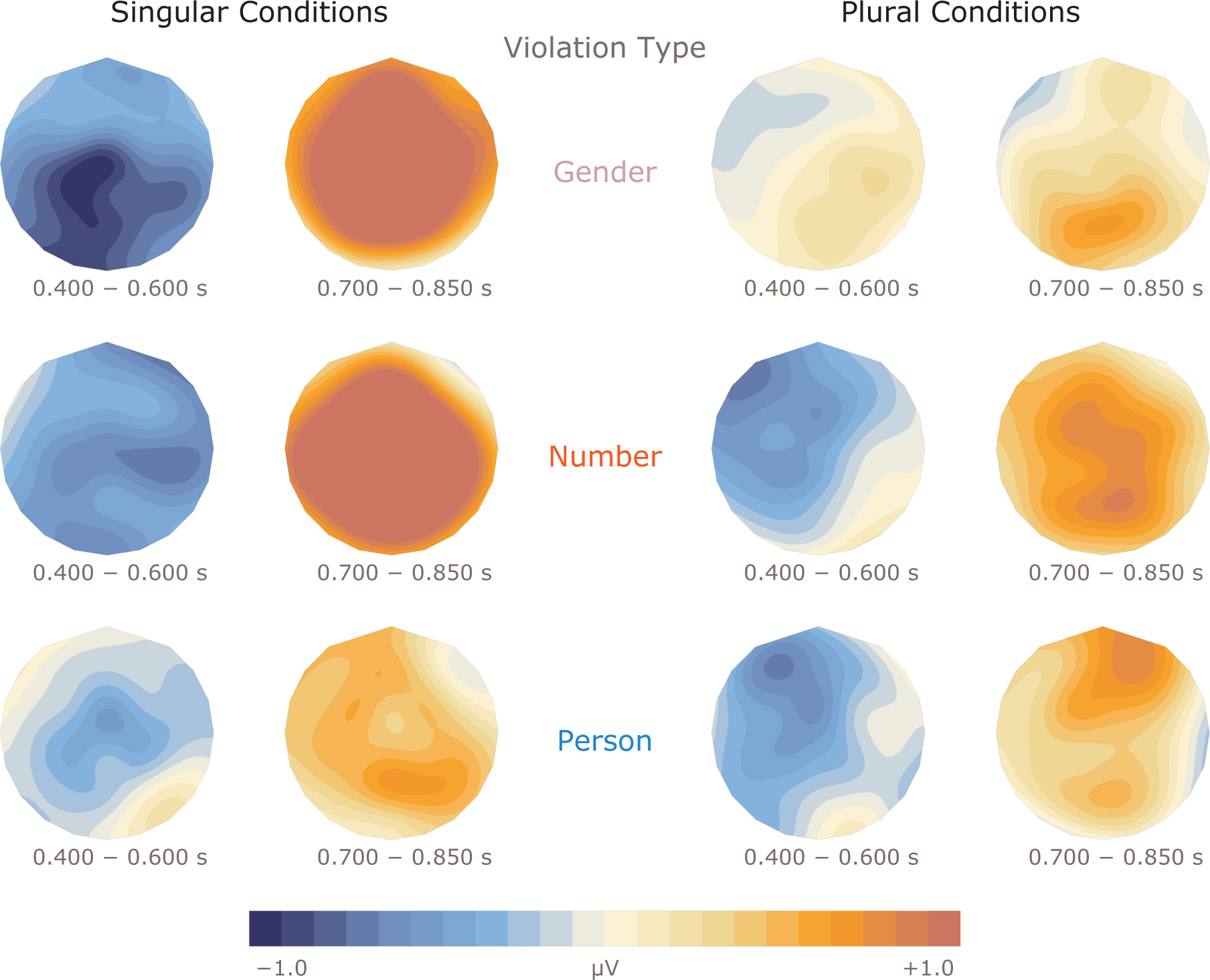
Topography of effects at the verb.

#### 3.2.1 Time-window 400−600 ms

The predominant effect in this time-window is the graded negativity engendered by the violation conditions as opposed to the acceptable conditions, showing clear differences in effects between the two subject-types. Number violations engendered a large broadly distributed negativity effect regardless of subject-type. The modulation of negativities evoked by person and gender violations differed based on whether the subject-type was singular or plural, with gender violations evoking a larger effect than person violations when the subject was singular, and person violations evoking a larger effect than gender violations when the subject was plural. Similarly, the topographic distribution of these negativities also varied, with the effect for person violations showing a central distribution, and that for gender violations showing a centro-parietal distribution. Table 5 and Table 6 show a summary of all the effects that reached at least marginal significance at the position of the verb in the 400−600 ms time-window for the lateral and midline regions respectively.

**Table 5:** ANOVA: ERPs at the Verb : 400−600 ms

| Effects in time-window 400–600 ms : Lateral Regions |  |  |  | DF | F | pHF |  |
| --- | --- | --- | --- | --- | --- | --- | --- |
| ◆ ST |  |  |  | 1,33 | 5.55 | 0.024 | ★ |
| ◆ CT |  |  |  | 3,99 | 3.24 | 0.025 | ★ |
| ↳<sub>CT</sub> = ACP+NUM | ◆ CT |  |  | 1,33 | 9.25 | 0.004 | ★★ |
| ◆ ST x CT |  |  |  | 3,99 | 3.03 | 0.032 | ★ |
| ◆ LATR x ST x CT |  |  |  | 9,297 | 2.37 | 0.035 | ★ |
| ↳<sub>LATR</sub> = LP | ◆ ST x CT |  |  | 3,99 | 4.54 | 0.005 | ★★ |
|  | ↳<sub>ST</sub> = S | ◆ CT |  | 3,99 | 6.31 | <.001 | ★★★ |
|  |  | ↳<sub>CT</sub> = ACP+NUM | ◆ CT | 1,33 | 5.76 | 0.022 | ★ |
|  |  | ↳<sub>CT</sub> = ACP+GEN | ◆ CT | 1,33 | 11.95 | 0.001 | ★★ |
|  |  | ↳<sub>CT</sub> = PER+GEN | ◆ CT | 1,33 | 9.70 | 0.003 | ★★ |
|  | ↳<sub>ST</sub> = P | ◆ CT |  | 3,99 | 2.74 | 0.050 | ★ |
|  |  | ↳<sub>CT</sub> = PER+GEN | ◆ CT | 1,33 | 5.38 | 0.026 | ☆ |
|  |  | ↳<sub>CT</sub> = NUM+GEN | ◆ CT | 1,33 | 8.87 | 0.005 | ★★ |
| ↳<sub>LATR</sub> = RP | ◆ ST x CT |  |  | 3,99 | 6.17 | <.001 | ★★★ |
|  | ↳<sub>ST</sub> = S | ◆ CT |  | 3,99 | 7.04 | <.001 | ★★★ |
|  |  | ↳<sub>CT</sub> = ACP+NUM | ◆ CT | 1,33 | 10.25 | 0.003 | ★★ |
|  |  | ↳<sub>CT</sub> = PER+NUM | ◆ CT | 1,33 | 6.25 | 0.017 | ★ |
|  |  | ↳<sub>CT</sub> = ACP+GEN | ◆ CT | 1,33 | 11.48 | 0.001 | ★★ |
|  |  | ↳<sub>CT</sub> = PER+GEN | ◆ CT | 1,33 | 10.56 | 0.002 | ★★ |

**Table 6:**
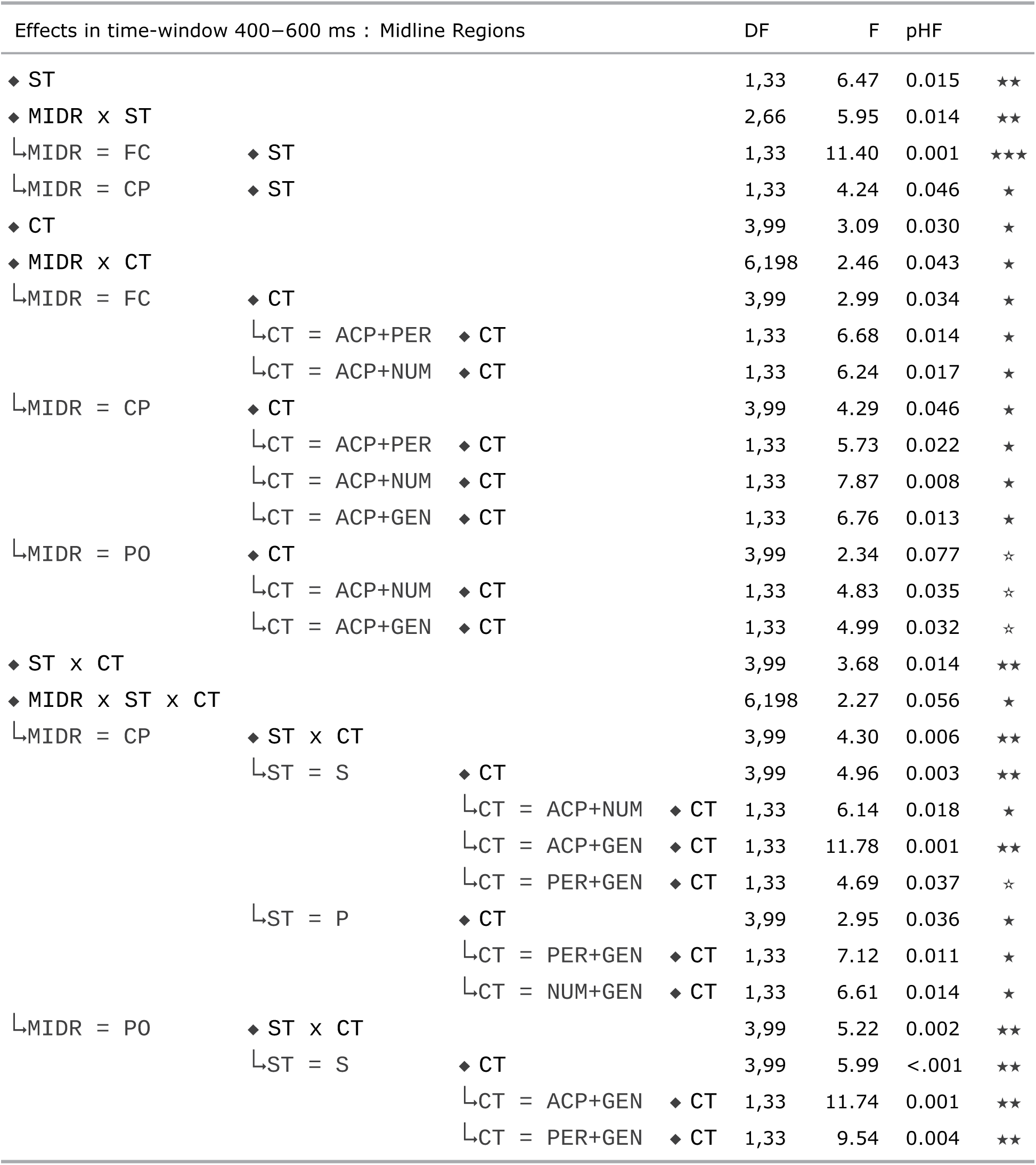
ANOVA: ERPs at the Verb : 400−600 ms

| Effects in time-window 400–600 ms : Midline Regions |  | DF | F | pHF |  |
| --- | --- | --- | --- | --- | --- |
| ♦ ST |  | 1,33 | 6.47 | 0.015 | ★★ |
| ♦ MIDR x ST |  | 2,66 | 5.95 | 0.014 | ★★ |
| ↳MIDR = FC | ♦ ST | 1,33 | 11.40 | 0.001 | ★★★ |
| ↳MIDR = CP | ♦ ST | 1,33 | 4.24 | 0.046 | ★ |
| ♦ CT |  | 3,99 | 3.09 | 0.030 | ★ |
| ♦ MIDR x CT |  | 6,198 | 2.46 | 0.043 | ★ |
| ↳MIDR = FC | ♦ CT | 3,99 | 2.99 | 0.034 | ★ |
|  | ↳CT = ACP+PER ♦ CT | 1,33 | 6.68 | 0.014 | ★ |
|  | ↳CT = ACP+NUM ♦ CT | 1,33 | 6.24 | 0.017 | ★ |
| ↳MIDR = CP | ♦ CT | 3,99 | 4.29 | 0.046 | ★ |
|  | ↳CT = ACP+PER ♦ CT | 1,33 | 5.73 | 0.022 | ★ |
|  | ↳CT = ACP+NUM ♦ CT | 1,33 | 7.87 | 0.008 | ★ |
|  | ↳CT = ACP+GEN ♦ CT | 1,33 | 6.76 | 0.013 | ★ |
| ↳MIDR = PO | ♦ CT | 3,99 | 2.34 | 0.077 | ☆ |
|  | ↳CT = ACP+NUM ♦ CT | 1,33 | 4.83 | 0.035 | ☆ |
|  | ↳CT = ACP+GEN ♦ CT | 1,33 | 4.99 | 0.032 | ☆ |
| ♦ ST x CT |  | 3,99 | 3.68 | 0.014 | ★★ |
| ♦ MIDR x ST x CT |  | 6,198 | 2.27 | 0.056 | ★ |
| ↳MIDR = CP | ♦ ST x CT | 3,99 | 4.30 | 0.006 | ★★ |
|  | ↳ST = S ♦ CT | 3,99 | 4.96 | 0.003 | ★★ |
|  | ↳CT = ACP+NUM ♦ CT | 1,33 | 6.14 | 0.018 | ★ |
|  | ↳CT = ACP+GEN ♦ CT | 1,33 | 11.78 | 0.001 | ★★ |
|  | ↳CT = PER+GEN ♦ CT | 1,33 | 4.69 | 0.037 | ☆ |
|  | ↳ST = P ♦ CT | 3,99 | 2.95 | 0.036 | ★ |
|  | ↳CT = PER+GEN ♦ CT | 1,33 | 7.12 | 0.011 | ★ |
|  | ↳CT = NUM+GEN ♦ CT | 1,33 | 6.61 | 0.014 | ★ |
| ↳MIDR = PO | ♦ ST x CT | 3,99 | 5.22 | 0.002 | ★★ |
|  | ↳ST = S ♦ CT | 3,99 | 5.99 | <.001 | ★★ |
|  | ↳CT = ACP+GEN ♦ CT | 1,33 | 11.74 | 0.001 | ★★ |
|  | ↳CT = PER+GEN ♦ CT | 1,33 | 9.54 | 0.004 | ★★ |

In the lateral regions, there were a main effects of subject-type and condition-type. Resolving the condition-type by comparing the condition-types pairwise showed a simple effect of condition-type for the comparison ACP + NUM (p = 0.004). The interaction subject-type x condition-type as well as the three-way interaction LATR x subject-type x condition-type were significant. Resolving the lateral ROI interaction for the individual regions showed that there was an effect of the interaction subject-type x condition-type in the left-posterior and right-posterior regions. This effect was resolved for subject-type in each concerned region, which showed that in the left-posterior and right-posterior regions, there was an effect of condition-type when the subject was singular. Resolving this further by comparing the condition-type pairwise revealed that there was a simple effect of condition-type for the comparisons ACP + NUM (all ps <= 0.022), ACP + GEN (all ps = 0.001) and PER + GEN (all ps <= 0.003) in both left-posterior as well as right-posterior regions. This effect reached significance for the comparison PER + NUM (p = 0.017) in the right-posterior region alone. In the left-posterior region, there was an effect of condition-type when the subject was plural, which when resolved by comparing the condition-types pairwise showed a simple effect of condition-type for the comparison NUM + GEN (0.005) and a marginal effect of condition-type for the comparison PER + GEN (0.026).

In the midline regions, there were main effects of subject-type and condition-type. The interaction MIDR x subject-type was significant, which when resolved for the individual midline ROIs showed a simple effect of subject-type in the front-central and centro-parietal midline regions. The interaction of MIDR x condition-type was likewise significant, which when resolved for the midline ROIs showed an effect of condition-type in the fronto-central and centro-parietal midline regions, and this effect was marginal in the parieto-occipital midline region. Comparing the condition-types pairwise in each of the three midline regions showed a simple effect of ACP + PER (all ps <= 0.022) in the front-central and centro-parietal midline regions. The effect of subject-type was significant for the comparison ACP + NUM (all ps <= 0.017) in the fronto-central and centro-parietal midline regions, whereas this effect was marginal (p = 0.035) in the parieto-occipital midline region. For the comparison ACP + GEN, there was a significant effect of subject-type in the centro-parietal midline region (p = 0.013), whereas this effect was marginal in the parieto-occipital midline region (p = 0.032). The three-way interaction MIDR x subject-type x condition-type was significant, which was resolved for the individual midline ROIs. This revealed an effect of the interaction subject-type x condition-type in the centro-parietal and parieto-occipital midline regions. Resolving this effect further, there was an effect of condition-type in both regions when the subject was singular, which when resolved by pairwise comparisons of the condition-types showed a simple effect of condition-type for the comparison ACP + GEN (all ps = 0.001) in both regions, for the comparison PER + GEN in the parieto-occipital region (p = 0.004), with this effect being marginal in the centro-parietal region (p = 0.037), and for the comparison ACP + NUM (p = 0.018) in the centro-parietal midline region. In the centro-parietal midline region, there was an effect of condition-type when the subject was plural, which when resolved using pairwise comparisons of condition-types showed a simple effect of condition-type for the comparisons PER + GEN (p = 0.011) and NUM + GEN (p = 0.014).

Therefore all the negativites that were significant in the 400−600 ms time-window showed a central and centro-parietal maximum. However, a left-lateral trend was apparent in the scalp topography of the negativities engendered by person violations when the subject was plural. Since the topography of negativity effects have important consequences for their functional interpretation (here, N400 versus LAN), we conducted a post-hoc analysis of the ERPs in an additional time-window, namely 300−500 ms, that was slightly earlier than the 400−600 ms time-window, specifically to test whether these effects perhaps show a left-anterior maximum in this early time-window. The predominant effect in this time-window was the graded negativity for the violation conditions as opposed to the acceptable conditions (becoming more prominent in the 400−600 ms time-window as reported above), with the gradedness differing based on whether the subject was singular or plural. Crucially however, there were no effects that reached significance in the left-anterior regions. Furthermore, there were no interactions involving the lateral regions of interest. Therefore we did not find any evidence in our data for a left-anterior maximum for any of the negativity effects. Nevertheless, the full statistics for the 300−500 ms time-window are provided for reference in the supplementary supporting information.

#### 3.2.2 Time-window 700−850 ms

The predominant effect in this time-window is the late-positivity effect for violation conditions as opposed to acceptable conditions. Crucially, this effect is not only graded based on the violating feature, but its modulation is also different based on the subject-type. The effect thus shows a four-way gradation when the subject is singular, with number violations evoking the largest positivity effect as opposed to acceptable sentences, followed by gender violations and then by person violations. By contrast, only a three-way difference ensues for plural subjects, with the largest positivity effect for number violations followed by smaller but virtually identical positivities for person violations and gender violations as opposed to acceptable sentences. Table 7 and Table 8 show a summary of all the effects that reached at least marginal significance at the position of the verb in the 700−850 ms time-window in the lateral and midline regions respectively.

**Table 7:**
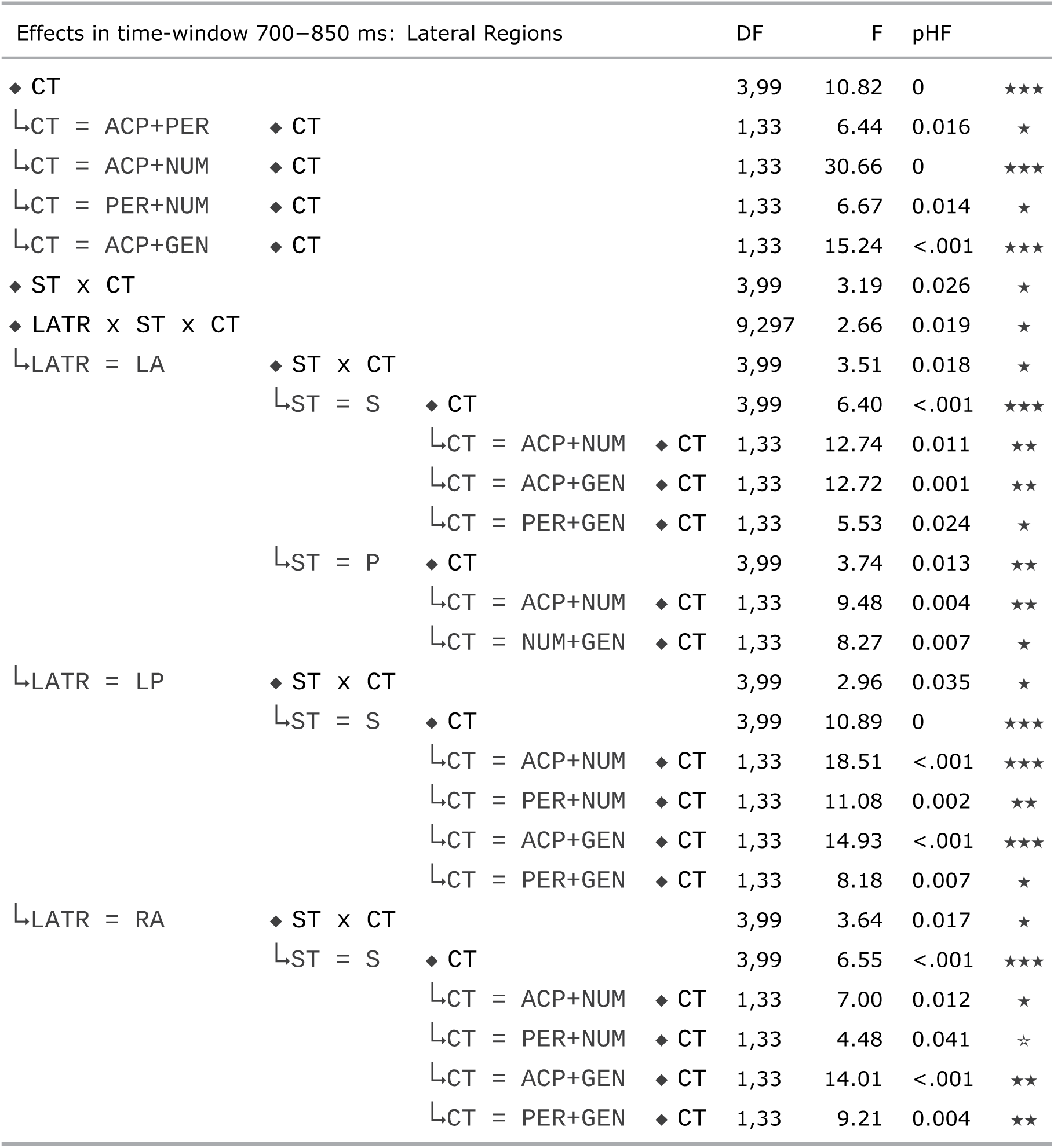
ANOVA: ERPs at the Verb : 700−850 ms

**Table 8:**
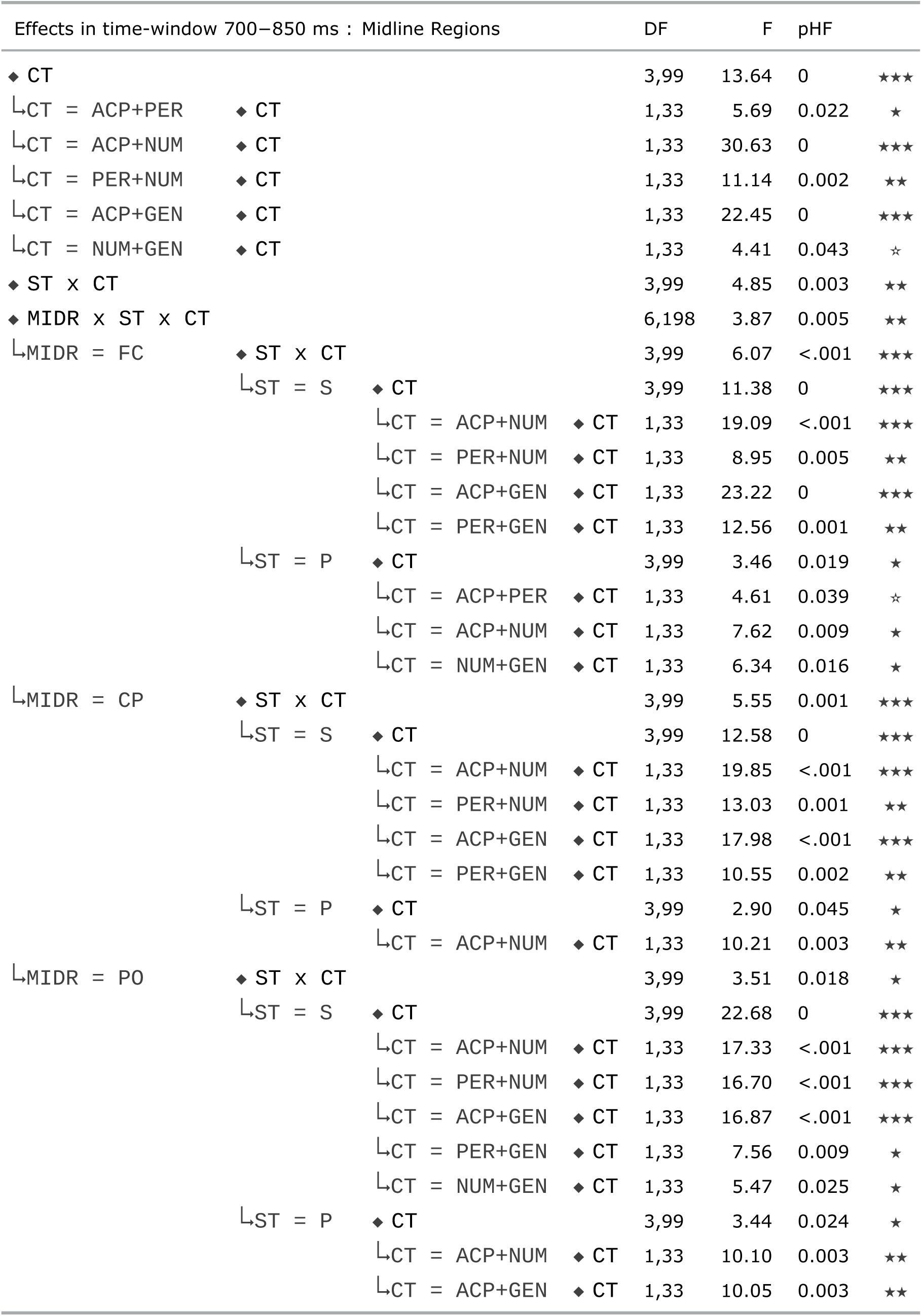
ANOVA: ERPs at the Verb : 700−850 ms

In the lateral regions, there was a main effect of condition-type, which was resolved resolved by comparing the condition-types pairwise. This showed a significant effect of condition-type for the comparisons ACP + PER (p = 0.016), ACP + NUM (p = 0), ACP + GEN (p < 0.001) and PER + NUM (p = 0.014) in all the lateral regions. The interaction subject-type x condition-type and the three-way interaction LATR x subject-type x condition-type were significant. Resolving this interaction for the individual lateral ROIs showed that the interaction subject-type x condition-type was significant in the left-anterior, left-posterior and right-anterior regions. This interaction was resolved for subject-type in all regions concerned, which showed that, when the subject was singular, there was an effect of condition-type in the left-anterior, left-posterior and right-anterior regions, which was resolved by comparing condition-types pairwise in these regions. This showed a simple effect of condition-type for the comparisons ACP + NUM (all ps <= 0.011), ACP + GEN (all ps <= 0.001) and PER + GEN (all ps <= 0.024) in all three regions concerned. There was an effect of condition-type for the comparison PER + NUM (p = 0.002) in the left-posterior region, and the effect was marginal in the right-anterior region. When the subject was plural, there was an effect of condition-type only in the left-anterior region, which when resolved further by comparing the condition-types pairwise revealed a simple effect of condition-type for the comparisons ACP + NUM (p =0.004) and NUM + GEN (p = 0.007).

In the midline regions, there was a main effect of condition-type, which when resolved by comparing the condition-types pairwise showed a significant effect of condition-type for the comparisons ACP + PER (p = 0.0022), ACP + NUM (p = 0), ACP + GEN (p = 0), PER + NUM (p = 0.002) and NUM + GEN (p = 0.043). The interaction subject-type x condition-type and the three-way interaction MIDR x subject-type x condition-type were significant. Resolving this interaction for the individual midline ROIs showed that the interaction subject-type x condition-type was significant in all three midline regions. This interaction was resolved for subject-type in all regions concerned, which showed that, when the subject was singular, there was an effect of condition-type in all three regions (all ps = 0), which was resolved by comparing condition-types pairwise in these regions. There was a simple effect of condition-type for the comparisons ACP + NUM (all ps < 0.001), ACP + GEN (all ps < 0.001), PER + NUM (all ps <= 0.005) and PER + GEN (all ps <= 0.009) in all the midline regions. This effect of condition-type was significant for the comparison NUM + GEN (p <= 0.025) in the parieto-occipital midline region. When the subject was plural, there was an effect of condition-type in all three regions (all ps <= 0.045), which was resolved by comparing condition-types pairwise in these regions. There was a simple effect of condition-type for the comparison ACP + NUM in all the regions (all ps <= 0.009). This effect of condition-type was marginal for the comparison ACP + PER (p = 0.039) and significant for the comparsion NUM + GEN (p = 0.016) in the fronto-central midline region. The effect of condition-type was significant for the comparison ACP + GEN (p = 0.003) in the parieto-occipital midline region.

### 3.3 Summary of results

The ERP results from our study can be summarised as follows.

- Violation of the agreement features elicited a biphasic negativity−late-positivity effect as opposed to acceptable sentences.
- However, this biphasic effect was graded based on the feature that was violated as well as the subject-type. Whilst number violations evoked the largest effects overall, the effects for person and gender violations were smaller in comparison, and showed a qualitative difference based on whether the subject-type was singular or plural.
- In the negativity time-window, a three-way distinction ensued that was different for the two subject-types. Number violations evoked a large broadly distributed negativity effect for both subject-types. Gender violations evoked a larger negativity effect as opposed to person violations when the subject was singular, with a centro-parietal maximum. Person violations evoked a larger negativity effect as opposed to gender violations as well as acceptable conditions when the subject was plural, with a fronto-central distribution. A visually apparent left-anterior trend for this latter effect was however not significant statistically.
- In the late-positivity time-window, there was a four-way gradation of the positivity effect when the subject was singular, whereas it was a three-way gradation when the subject was plural. Number violations evoked the largest positivity effect in both subject-types. Gender violations evoked a larger positivity effect compared to person violations when the subject was singular, but no such difference ensued when the subject was plural.

Based on the latency and topography of effects as well as the experimental conditions, the negativity effects in our study can be plausibly interpreted as instances of an N400 effect^3^ and the late-positivities as instances of a P600 effect. Agreement violations of the sort we employed have been traditionally viewed as formal morphosyntactic rule violations, which are associated with concomitant LAN effects that are interpreted as indicative of the detection of the morphosyntactic violation (Münte, Szentkuti, Wieringa, Matzke, & Johannes, 1997; Münte, Matzke, & Johannes, 1997; Friederici, 2002; Bornkessel & Schlesewsky, 2006). Further, the likelihood of observing a LAN has been thought to be in direct proportion to the morphological richness of a language (Friederici & Weissenborn, 2007), which nevertheless is said to depend upon whether morphological marking is crucial for assigning syntactic roles in the given language (Friederici, 2011). However, Molinaro, Barber, & Carreiras (2011) draw attention to the fact that the nature of the complexities involved in morphological decomposition for feature identification may dissociate whether a LAN or, alternatively, an N400 ensues. Indeed, Choudhary et al. (2009) provide converging evidence from Hindi that this dissociation results from whether an interpretively relevant cue is violated (in which case an N400 ensues), or alternatively the violation involves a cue that is irrelevant for interpretation. In view of this, the absence of a LAN effect and the presence of an N400 in our study is not surprising, given that agreement computation in Arabic is not always predictable based on the features of the agreement controller alone, but is dependent upon specific syntactic properties of the construction involved, such as word-order and whether or not the subject is overt, as well as properties at the syntax-semantic interface such as humanness (see below for a detailed discussion). The P600 effects in our study can be plausibly interpreted as reflecting repair or reanalysis processes associated with agreement violations (Friederici, 2002, 2011; Bornkessel & Schlesewsky, 2006), thought to be triggered by domain-general conflict monitoring processes (Van de Meerendonk, Chwilla, & Kolk, 2013).

## 4 Discussion

We investigated the processing of subject-verb agreement in simple intransitive Arabic sentences in this ERP experiment, and present here the findings from one of the first neurocognitive studies examining sentence level processes in Arabic, thus providing first insights into the online processing of the language.

### 4.1 Hierarchical modulation of effects

As illustrated by the topographic map of effects in Figure 5, our results show that violations of the features not only modulate the biphasic pattern of ERPs, but they also show a qualitative difference in their modulation based on whether the subject was singular or plural. The modulation of effects for the violation conditions as opposed to the acceptable conditions in our study can be represented as a hierarchy as follows:

(2) Modulation of N400 Singular subject conditions : Number / Gender > Person Plural subject conditions : Number / Person > Gender
(3) Modulation of P600 Singular subject conditions : Number > Gender > Person Plural subject conditions : Number > Person / Gender

A handful of previous studies investigating processing differences between agreement features (in the subject-verb context or otherwise) have indeed reported a modulation of late-positive P600 effects, albeit for across subject-type comparisons (Kaan et al., 2000; Deutsch & Bentin, 2001) or for combined versus single feature violations (Nevins et al., 2007; Zawiswewski et al., 2016; Alemán Bañón & Rothman, 2016).^4^ However, to our knowledge, a modulation of negativities (LAN or N400) has not been reported to date in the context of agreement violations involving all three features^5^, and ours may be one of the first studies on agreement processing to report such a hierarchical modulation of the N400 based on the agreement feature involved.

The modulation of effects in our study, especially of the negativity, cannot be satisfactorily explained simply based on differences in the complexity between the conditions, nor the frequency of a certain type of violation, nor orthographic salience, nor even the physical characteristics of the morphosyntactic markings per se. This is because, as we have described in Section 2.3, for a given verb in each subject-type, the material preceding it was always identical. This meant that, at the position of the verb, potential effects of differences in lexical frequency and orthographic or morphological complexity apply to all conditions identically equally, and therefore do not confound the interpretation of language-related effects of interest at the verb. Physical characteristics of the stimuli such as the visual length of the verb appear to be comparable across conditions, despite minor differences in the P200 amplitudes that were nevertheless not significant (see Figure S4).

Rather, we argue that these results suggest that the processing system takes into account the relative salience of the features in the given language. If so, this would be in line with the view put forward by Molinaro, Barber, & Carreiras (2011) in their exhaustive review of cross-linguistic findings on processing agreement, namely that the processing of agreement dependencies is sensitive to the feature involved as well as the constituents expressing the dependency, and that this can result in modulations of negativity effects (such as LAN and/or N400) and late-positivity effects^6^. Specifically, the processing of subject-verb agreement is said to involve semantic factors in addition to morphosyntactic information (Molinaro, Vespignani, Zamparelli, & Job, 2011), and the extent to which lexical-semantic factors play a role in computing subject-verb agreement in a given language is said to have a direct bearing on whether a LAN or an N400 effect is engendered (Molinaro, Barber, Caffarra, & Carreiras, 2015).

It may be of some relevance here to briefly turn to an ongoing debate in the ERP literature on morphosyntactic processing, which relates to inter-participant variation of ERP effects. The claim is that most of the LAN effects reported for agreement processing in the literature may simply be artefactual resulting from components that partially overlap temporally and spatially, which happen to be aggregated together due to the grand-averaging of individual ERPs (Tanner, 2018, 2015; Tanner & Van Hell, 2014). Whilst we did not find evidence for a LAN effect in our study, it is important to consider whether individual variation amongst our participants might be able to explain some of our results. However, several points speak against such a possibility. Firstly, individual variation, if any, would have been systematic across conditions for a given participant. Secondly, in view of our within-participants design, any such variation would have equally contributed to all the critical conditions since all of them consisted of the same type of violation, namely a violation of an agreement feature between the subject and the verb. Thirdly, as Grey, Tanner, & Van Hell (2017) have reported, variation in ERP effects engendered during morphosyntactic processing across individuals is said to be minimal in case of dominant right-handers (as opposed to left-handers). All our participants were dominantly right-handed individuals (see Section 2.5). Crucially therefore, given that any potential artefactual outcome resulting from the grand-averaging procedure would be equally applicable to all our critical conditions, individual variation may not fully account for, nor nullify, the subject-type specific differences in the modulation of effects we found in our study.

### 4.2 Salience-weighted Feature Hierarchy

In view of the agreement feature hierarchy in (1) originally proposed based on cross-linguistic distribution of these features (Greenberg, 1963), the graded effects in the present study are not surprising. Rather, our results provide neurophysiological evidence for the idea that the relative position of a feature in the hierarchy directly reflects its relative cognitive salience (Harley & Ritter, 2002, but see below) and thereby its relative importance in language processing (Carminati, 2005) in comparison to other features. Nevertheless, the modulation of effects found in our study suggests that the feature hierarchy based on distributional properties and conceptual or cognitive aspects of the features may differ from a hierarchy based on the relative language-specific salience of the features, which need not be identical across languages. That is, the hierarchy of feature salience appears to differ based on language-specific properties, and may exhibit slight variations even within a given language depending upon the specifics of the construction involved and general language-use in the speech community as well as the way each feature is morphologically encoded in the language. We argue below that this variation is neither an aberration nor arbitrary. Rather, it is systematic based on the relative weightings of the salience of a given feature in a language.

We postulate a salience hierarchy of agreement features that is language-specific rather than universal. The properties of a given language as well as the relative salience of the agreement features determine the hierarchy and thereby differentially modulate language comprehension. If the hierarchy is flat such that there is minimal difference in salience between the features, then no modulation of ERP effects should be expected for single feature violations in the given language. If there is a hierarchical difference in salience between the features, then there would be a concomitant modulation of ERP effects, such that violating a highly salient agreement feature evokes a larger effect compared to violating a less salient feature.

The idea that some properties in a language may be relatively more salient than others for language comprehension is not new. This was first put forward in the context of the Competition Model (e.g., MacWhinney et al., 1984; Bates & MacWhinney, 1989; Bates, 1999), in which such linguistic properties or cues with language-specific weightings interact during language comprehension. Agreement is one such cue, the strength of which has been shown to vary across languages (for an overview, see Bates et al., 2001). Within the domain of online language processing, Bornkessel-Schlesewsky & Schlesewsky (2009) have proposed that information types such as case-marking, word-order, animacy etc., called prominence information, all interact during online language comprehension, and that their relative prominence depends upon their language-specific weightings. For instance, word-order has higher weighting in English than, say, animacy, and determines the argument roles, whereas in a language like Turkish, case is more prominent, and so on. Furthermore, in a recent article, Bornkessel-Schlesewsky & Schlesewsky (2019) have argued for a neurobiologically plausible model, whereby they posit that all the language-related negativities form a family of functionally related rather than distinct negativities, and that amplitude modulations of negativities are said to reflect ‘precision-weighted predictive coding errors, with precision (the inverse of variance) reflecting the relevance of a particular stimulus feature in a given language (Bornkessel-Schlesewsky & Schlesewsky, 2019, p. 11)’.

We extend the idea of weightings to the domain of agreement features, such that they are organised on a salience-weighted hierarchy that modulates language comprehension, with differences in their salience due to specific properties of a given language determining their position on the hierarchy. Further, this hierarchy interacts with other prominence scales and discourse context such that the modulation of effects may vary within the language based on the specific type of construction (Mancini et al., 2011a). This proposal would parsimoniously account for the fact that ERP effects for subject-verb agreement violations show a graded hierarchy in certain languages / constructions, whereas such a pattern does not ensue in others.

### 4.3 Language-specific feature weightings

In support of our proposal that the hierarchy of agreement features may very well be language-specific based on the relative salience of the features in the given language, we motivate here a language-internal explanation of how these different weightings come about in Arabic.

As mentioned in Section 1.4 when motivating Arabic as a suitable language for the purpose of investigating our research question, verb agreement is not merely dependent upon the features of the subject noun in Arabic. Specifically, number and gender agreement in particular involve complex interactions with syntactic and semantic properties of a sentence and its event participants, such that verb agreement in these features is not always determined by the number and gender properties of the subject noun concerned.

To reiterate the salient points about the Arabic agreement system, number agreement is not only dependent upon the number property of the subject noun, but also upon the relative order of the verb and the subject as well as whether or not the subject is going to be overt. In other words, under certain circumstances, an identical verb form is sometimes correct and sometimes incorrect based on factors that go beyond the number property of the subject noun per se. Thus, a simple feature-matching between the number property of the subject and the verb would not always lead to the correct interpretation.

Similarly, gender agreement depends upon an interplay betweeen the number feature and the humanness property of the subject noun, whereby plural masculine and feminine nouns require the verb to show full agreement if the noun is human, whereas for non-human (animate and inanimate) plural nouns, regardless of whether they are masculine or feminine, the verb must show singular-feminine agreement. Gender contrast is preserved in human subjects but neutralised in non-human subjects. In effect, determining the correct gender agreement for a given utterance is dependent upon the gender, number as well as the position of the subject noun on the animacy hierarchy (specifically humanness).

By contrast, the person feature does not exhibit any variation of this sort in Arabic as far as agreement is concerned. In other words, an agreement violation involving the person feature is always reliably a violation, and therefore, the processing system need not consider global properties of the language when evaluating person agreement in Arabic.

Therefore, evaluating number and gender agreement involves factors beyond the number and gender properties of the noun, which co-determine the processing of these features. In other words, number and gender, in that order, are more salient in Arabic for processing subject-verb agreement than person. This would explain the hierarchy of effects we observed in our study for singular subjects. However, results for plural subject nouns would suggest that the hierarchy is qualitatively different from that for singular subjects, in that the gender feature may be less salient than the person feature for plural subjects.

There may be at least three language-internal reasons for this difference. First, when speaking about or adressing a group of individuals involving both women and men, speakers of Arabic overwhelmingly tend to use masculine agreement.

Second, and related to the previous point, many spoken varieties of Arabic show no gender distinction between the masculine and feminine in the plural, with the masculine plural affix used as morphological marker for both. That is, based on language-use, a verb that shows masculine agreement for a group of individuals also involving women is not a violation in Arabic. Third, the humanness interaction described above, namely that non-human masculine plural nouns require the verb (and adjectives etc.) to show singular feminine agreement across the board, means that roughly half of the utterances involving masculine plural nouns require feminine agreement. That is, a verb that shows feminine agreement in the context of masculine plural subjects (albeit non-human) is not always a violation either. Simply put, both mismatching paradigms, namely, masculine subjects requiring feminine verb agreement and subjects involving feminine referents requiring and/or allowing masculine verb agreement, exist in the language. As a consequence, gender agreement mismatches involving plural nouns (masculine or feminine) are not always violations, which explains the fact that the effects for gender violations in our study were smallest for plural subjects.

### 4.4 Implications and Outlook

Previous results from morphologically rich languages such as Hindi and Basque are of particular relevance here. In Hindi, in which subject-verb agreement is purely based on the features of the nominative subject noun, Nevins et al. (2007) have found that no modulation of effects ensued based on whether number or gender is violated. Although their study did not include a pure person violation, there was a combined person+number violation condition, which elicited a larger P600 effect. Furthermore, the P600 effect for the combined violation of person+number was larger than that for both the combined number+gender violation and individual number and gender violations. A potential caveat for the larger P600 in their study for person+number violations may have been the relative infrequency of this condition in comparsion to the other violation conditions in their stimuli. Remaining ambivalent about whether or not the infrequency of this condition played a role, the authors attributed the larger P600 effect to the increased orthographic salience of person violations in Hindi rather than an additive effect, but added that it may be due to a cross-linguistically privileged status of the person feature. In Basque, Zawiswewski et al. (2016) have reported a larger P600 effect for person violations as opposed to number violations. By contrast, the effect for a combined violation of person+number did not differ from that for a person violation, with these two conditions showing an equally larger P600 effect compared to number violation. This would imply that the person feature is more salient in Basque, and that a combined violation of a more salient and less salient feature would not lead to additive effects. In the first instance, these results may appear to suggest that the person feature is somehow more significant and especially salient than the other features. However, two studies that specifically investigated person agreement are a case in point here. Seemingly anomalous, but alternatively interpretable person agreement (the so-called ‘unagreement’) in Spanish (Mancini et al., 2011b) and Basque (Mancini et al., 2019) evoke negativities (albeit with topographic and latency differences) but no P600 in comparison to outright and irresolvable person violations. Taken together, these results and the results from our study do not speak for a universal special status for the person feature, but rather provide converging evidence to our claim that modulation of effects correlate with language-specific differences in the salience of a given feature.

Consequently, we argue that a parsimonious account of apparently contradictory results from Hindi and Basque mentioned above on the one hand and our results from Arabic on the other would be along the lines of what we propose here, namely that the agreement features person, number and gender may show different hierarchies (or no hierarchies at all) depending upon the specific properties of a language, based on their relative salience in the given language, with differences in language-specific weightings resulting from a number of language-internal reasons.

Our proposal here is not entirely incompatible with the idea of orthographic salience contributing to the larger P600 effect for the combined person+number violation in Hindi. However, a more general explanation for the increased salience of the person feature in Hindi may be that, whereas number and gender are expressed on the main verb even in the perfective aspect when the agent is in ergative case, person marking is underspecified by contrast, and only available from the auxiliary in the perfective aspect. In the imperfective aspect, when the person feature is exclusively available on the main verb as in the stimuli employed by Nevins et al. (2007), it is more prominent and salient, and a violation of this highly salient feature would have led to a concomitantly larger P600 effect in their study. Similarly, in the Basque study by Zawiswewski et al. (2016), the word-final subject morpheme in the auxiliary verb for the number violation was orthographically minimally different from that for the grammatical condition (*\*-zue* versus *-zu*), whereas the morpheme indicating person and person+number violations were quite distinct (*\*-t* and *\*-gu* versus *-zu*), and therefore much more salient visually and phonetically. This may have contributed to the equally larger P600 for these latter violations as opposed to number violations in that study. The behavioural data that the authors have reported strongly supports this claim, such that the violations involving the distinct morphemes were almost always accurately detected as such, and much quicker, in comparison to number violations, which were detected with a lower accuracy, and at a slower pace.

In sum, existing results from Hindi and Basque can be accounted for based on the increased salience of the person feature (in the Hindi study, due to the orthography and exclusive availability of person in their stimuli; in the Basque study, perhaps due to the orthographic distinctness of the morpheme indicating person violation, but see below), without having to resort to a universally privileged or significant status for the person feature. To reiterate what we have stated earlier in Section 4.2, a potential special status for a feature posited based on its conceptual and cognitive aspects may differ from its linguistic status in a given language based on its salience in the agreement system of the language concerned. In other words, the person feature is more salient in Hindi and Basque, albeit due to very different reasons, leading to a similar pattern of results in these languages. By contrast, the person feature seems to be the least salient feature in Arabic in comparison to number or gender, as explained earlier. Our proposal here of a language-specific salience-weighted feature hierarchy parsimoniously accounts for these results as well as qualitative ERP differences that have been sometimes reported for the different feature violations within a given language, such as the results from Spanish (Mancini et al., 2011a). Further, it satisfactorily provides a general and overarching account for cross-linguistic differences in the pattern of results reported for subject-verb agreement violations, such as in Dutch (Kaan et al., 2000) and Hebrew (Deutsch & Bentin, 2001), among others.

Compelling evidence in support of our proposal for a salience-weighted feature hierarchy that would preclude a universal special status for a given agreement feature comes from a recent study on Basque, in which Martinez de la Hidalga et al. (2019) investigated the processing differences between person and number agreement in unaccusative verbs (e.g., ‘*to fall*’) versus unergative verbs (e.g., ‘*to run*’), and have reported N400 modulations based on the feature violated, and P600 modulations that differed based on the feature violated and the verb type. Person violations elicited a larger N400 effect than number violations in both verb-types. Interestingly, number violations elicited a larger P600 effect as opposed to person violations when the verb was unaccusative, whereas person violations elicited a larger P600 effect as opposed to number violations when the verb was unergative, thus showing the opposite pattern. The authors interpreted these results as a result of salience differences between the two verb-types: unergative verbs involve agent arguments that are commonly highest on the animacy hierarchy (i.e., human) and therefore the person feature is more salient for unergative verbs, thus resulting in a larger P600 for person violations compared to number violations. By contrast, unaccusative verbs involve theme arguments that are often lowest on the animacy hierarchy and thus the person feature is less salient, resulting in the opposite pattern for unaccusative verbs. Thus the results of this study strongly support our argument here, namely that a conceptual or cognitive special status for a feature (in this case, person) may not always translate to a similarly higher level of salience at a linguistic level universally across languages. Further, in line with our own findings, they neatly illustrate that differences in salience weightings may lead to different hierarchies for the same feature across different constructions within a given language depending upon a number of factors that are specific to the language concerned.

Furthermore, the concept of salience-based weightings enables generating specific hypotheses about agreement processing in understudied languages. For instance, whilst we do not yet have any independent evidence from Arabic for whether or not multiple feature violations would lead to additive effects when the features are on a hierarchy based on their salience differences, results from Hindi and Basque point to the fact that an additive effect need not necessarily ensue. Nevertheless, further research is necessary to investigate the question in detail. Furthermore, if agreement is always regular in a language and there are no apparent salience differences between the features based on the properties of the language concerned, then no modulation of effects should be expected, whereas in languages in which the features have different salience weightings either due to orthographic reasons (as in Hindi), or syntactic or semantic reasons (as in Arabic or Basque), then modulations of effects should be expected.

Our proposal for a Salience-weighted Feature Hierarchy is conceptually in line with the agreement feature hierarchy (Greenberg, 1963) and the Feature Strength Hypothesis (Carminati, 2005), and provides neurophysiological evidence for the idea that agreement processes involve syntactic, semantic and pragmatic factors all at once (Eberhard et al., 2005; Vigliocco et al., 1995). However, our findings do not support a universal special status for specific features (say, person) in language processing purely based on their distributional and cognitive or conceptual properties across languages. Rather, the crucial difference between what we postulate here and previous proposals is that, the hierarchy of features may differ based on language-specific characteristics and salience, whereby the relative position of a feature depends upon the language-speicific weighting of the feature. Our proposal parsiomoniously accounts for the fact that ERP effects for agreement violations may show a graded hierarchy in certain languages / constructions, whereas such a pattern may not ensue in others.

### 4.5 Conclusion

In the study presented here, we investigated the processing differences between person, number and gender agreement in Arabic, a widely-spoken but understudied language. One of the first neurophysiological studies examining sentence level processes in Arabic to date, the findings from our study are an important addition to existing cross-linguistic results on online language comprehension. Thanks to the within-language within-participants systematic comparison of all three agreement features in the context of subject-verb agreement in our study, we were able to show for the first time that the agreement features differentially modulate language comprehension based on their relative salience. The salience weightings of features may and do differ across languages depending upon the specific properties of the language concerned, thus giving rise to a hierarchy of agreement features that is language-specific rather than universal. Such a Salience-weighted Feature Hierarchy, we argue, would parsimoniously account for the diversity of existing cross-linguistic neurophysiological results on verb agreement processing, without having to resort to a universal special status for a certain feature, nor having to assume a universally static hierarchy of features that does not take into account the typological properties of individual languages. Furthermore, our proposal enables generating specific testable hypotheses about agreement processing in languages that are as yet unstudied or understudied. In this respect, our study is an important contribution towards understanding how the human brain processes and comprehends vastly diverse languages with equal ease and poise.

## Acknowledgements

The research reported here conducted in the context of several research visits by R. Muralikrishnan was partly supported by a Research Institute grant (G1001) from New York University in Abu Dhabi to Prof.Alec Marantz, whom we should like to thank here. Ali Idrissi was at the United Arab Emirates University in Al Ain during the study.

## Compliance statement

We report how we determined our sample size, all data exclusions, all inclusion / exclusion criteria, whether inclusion / exclusion criteria were established prior to data analysis, all manipulations, and all measures in the study. No part of the study procedures / analyses was pre-registered prior to the research being conducted. Data and analysis code will be made available upon request by contacting the corresponding author.

## Supplementary Materials

### S1 Supplementary Materials

#### S1.1 Arabic agreement examples

The minimal pair examples provided in the following pages illustrate the important aspects of the Arabic agreement system. Examples A1 to A4 contain human subjects, B1 to B4 contain non-human animate subjects, C1 to C4 contain non-human inanimate subjects. In each set, x1 and x2 contain masculine nouns (in of the respective animacy classes A, B or C as mentioned above), and x3 and x4 contain feminine nouns. Similarly, in each set, x1 and x3 contain singular nouns, x2 and x4 contain their plural counterparts. Each sentence is provided in the SV and VS orders. At the end, the dropped-subject variants of examples A2 and A4 are provided in A2-DS and A4-DS respectively.

The following are the salient points to be noted:

- In all sets, regardless of the number property of the subject noun, the verb in the VS order is always singular. This illustrates the agreement asymmetry based on word-order (Aoun et al., 2010).
- In sets B and C, except when the subject noun is singular masculine, the verbs and adjectives show singular feminine agreement across the board, regardless of the gender of the non-human (animate or inanimate) noun. This illustrates deflected agreement (Ferguson, 1997).
- When a plural subject is not going to be overt (i.e., dropped) in a sentence, the verb must show full number agreement. The dropped-subject variants of examples A2 and A4 illustrate this.

We thank Moustafa Mansour (p.c.) for these minimal pair examples.

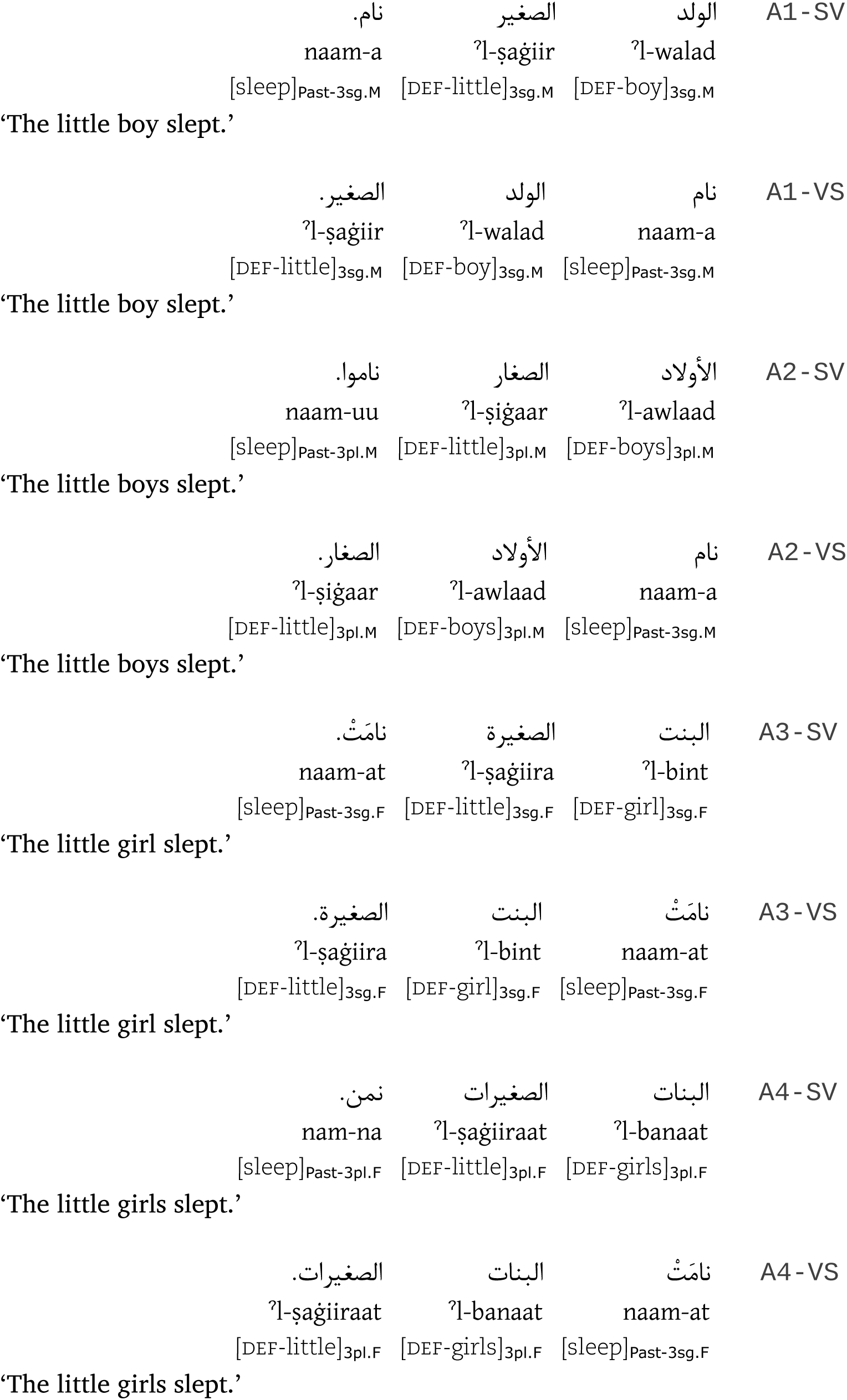
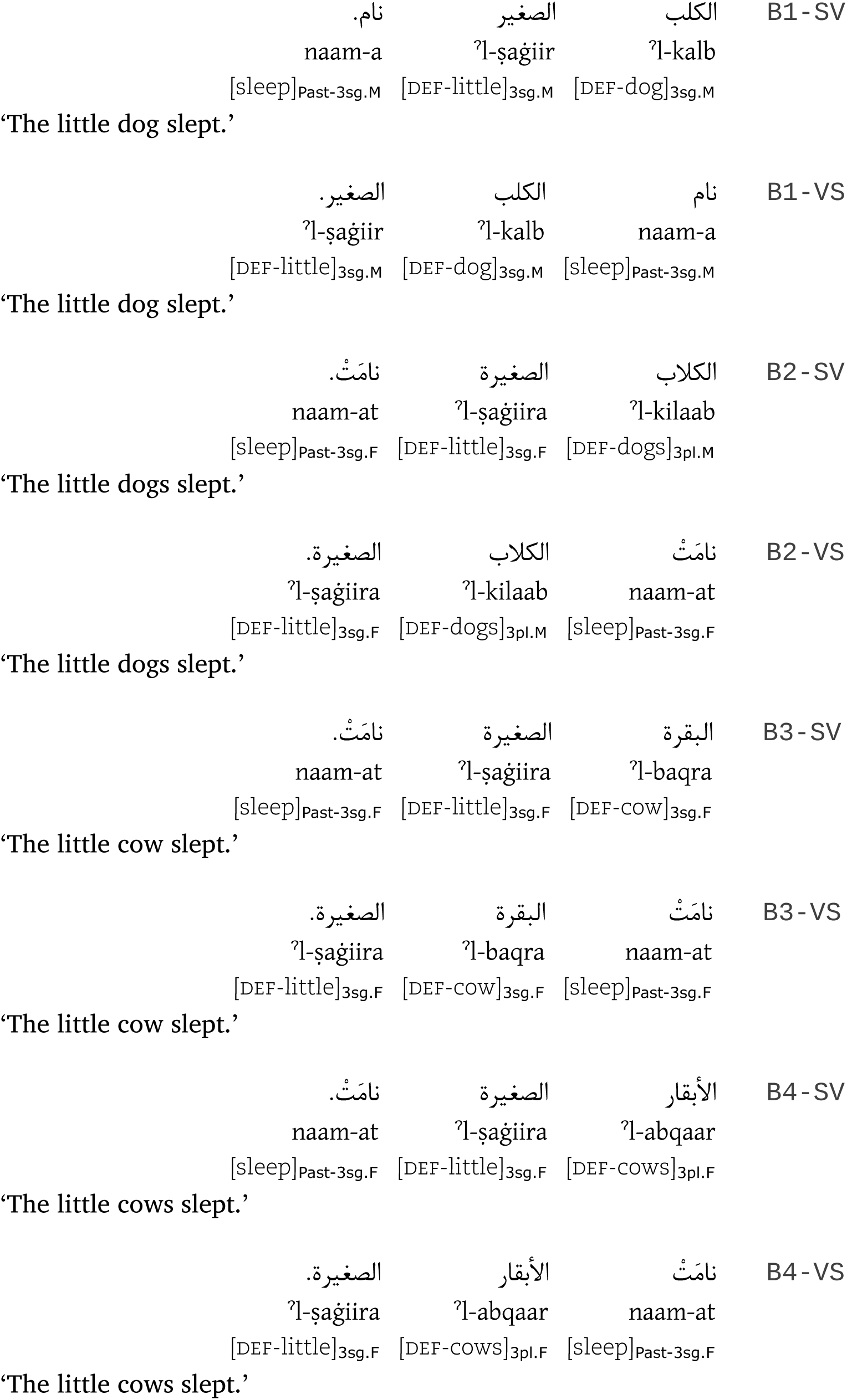
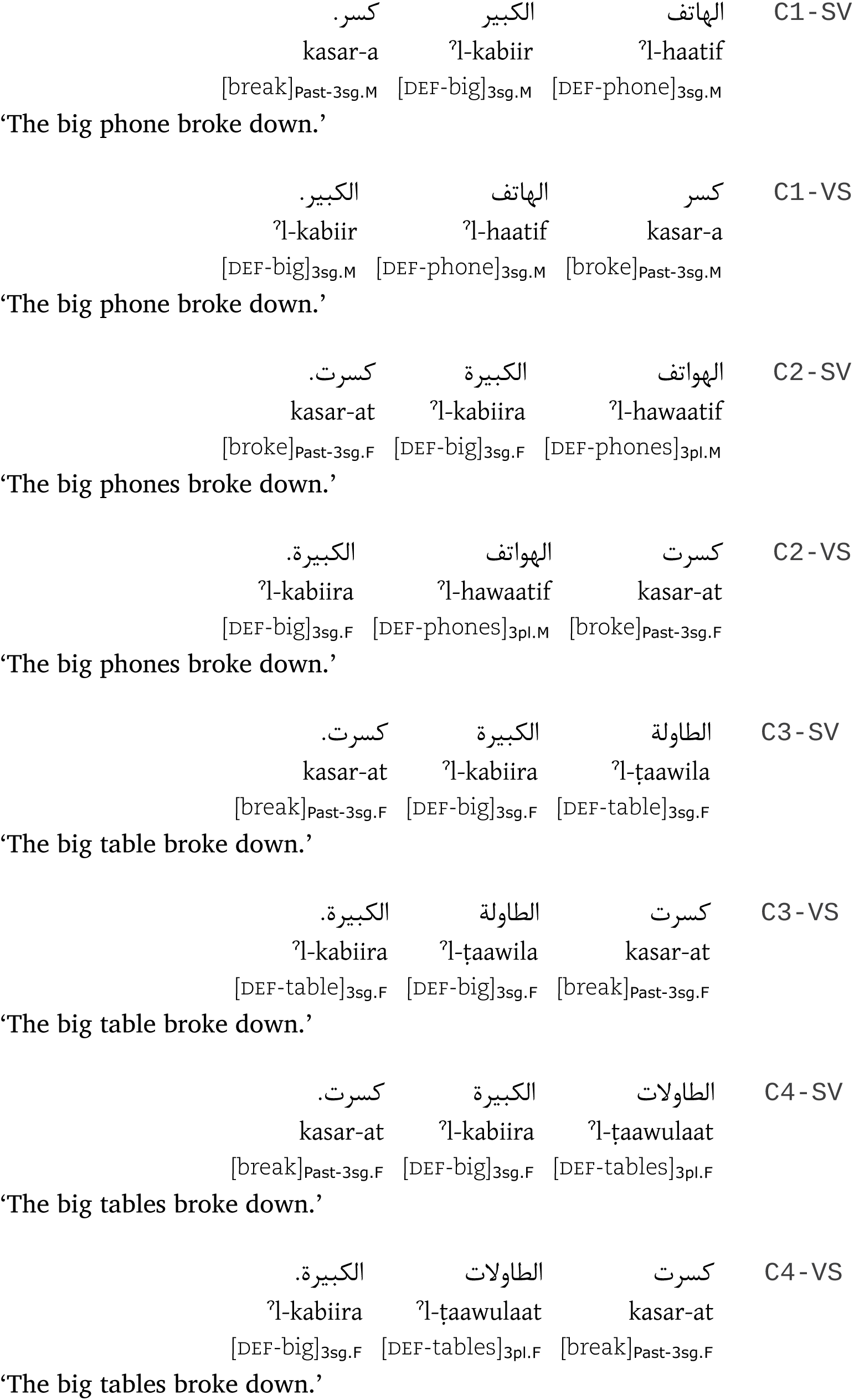
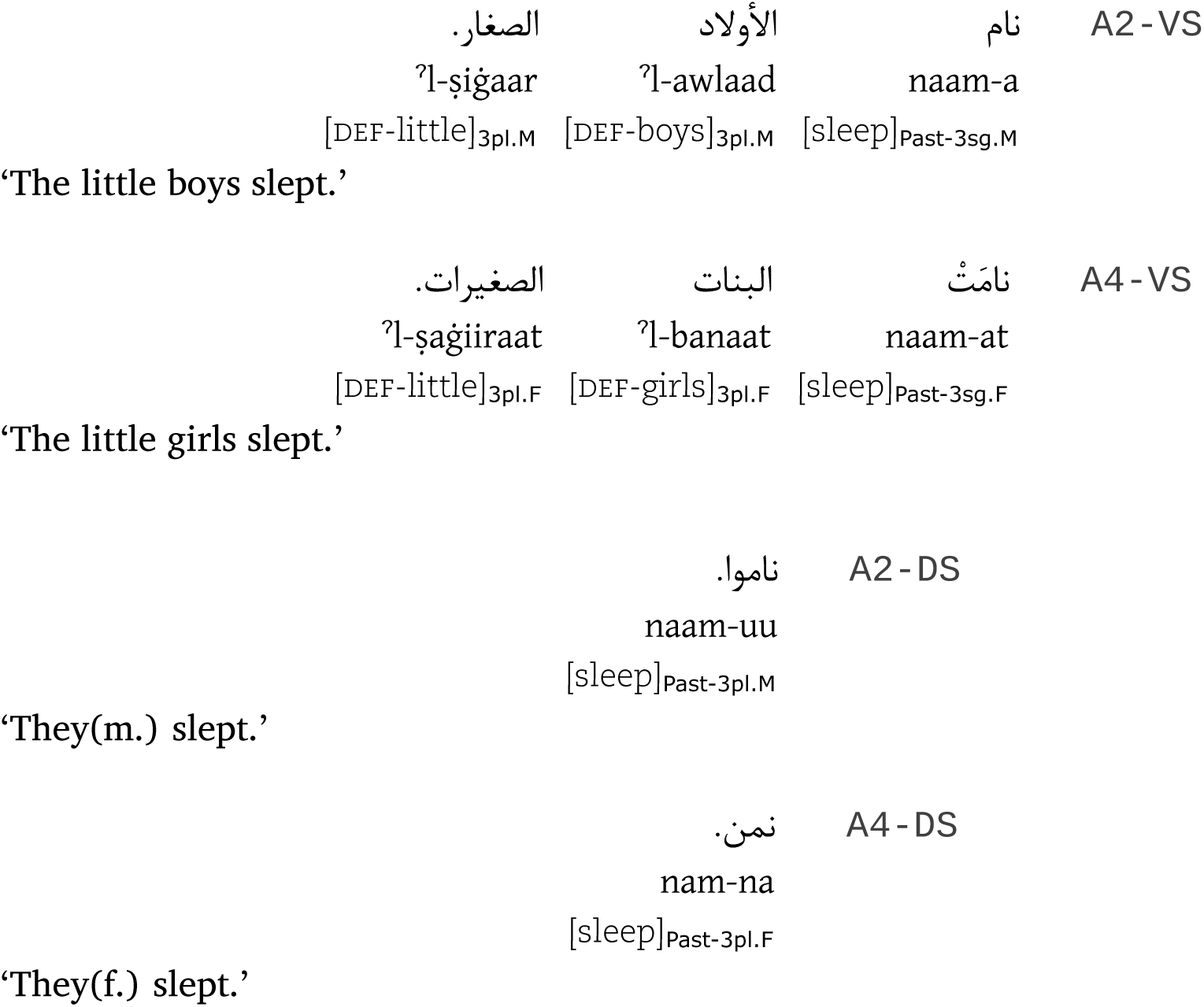

#### S1.2 Confirmatory post-hoc analyses

##### S1.2.1 Time-window 300−500 ms

As mentioned in the main article, we analysed the data in the 300−500 ms time-window post-hoc specifically to test whether we can find evidence for a LAN effect in our data. As the data presented here show, we could not find any evidence for the same. This time-window does not convey any new information other than confirming the pattern of effects found in the 400−600 ms time-window reported in the main article. In view of this, and the fact that it partially overlaps in time with the one mentioned above, we do not discuss data from this time-window. Nevertheless it is provided here for reference.

The predominant effect in this time-window is the graded negativity for the violation conditions as opposed to the acceptable conditions, with the gradedness differing based on whether the subject was singular or plural. Table S1 shows a summary of all the effects that reached at least marginal significance at the position of the verb in the 300−500 ms time-window.

There were no main effects. The interaction ROI x subject-type was significant in the midline regions, which when resolved for the individual ROIs showed a simple effect of subject-type in the fronto-central midline region. The interaction subject-type x condition-type was significant in the lateral and midline regions. This effect was resolved in the lateral regions alone (due to a ROI interaction in the midline regions), which showed an effect of condition-type when the subject was singular. This was resolved further by comparing the condition-types pairwise, which showed a simple effect of condition-type for the comparison PER + NUM alone.

The three-way interaction ROI x subject-type x condition-type was significant in the midline regions alone, which was resolved for the individual ROIs. This showed a marginal effect of subject-type x condition-type in the fronto-central midline region, and the effect was significant in the central, centro-parietal, parietal and parieto-occipital midline regions, which was resolved for subject-type in each region. When the subject-type was singular, there was an effect of condition-type in the central, centro-parietal, parietal and parieto-occipital midline regions. Resolving this effect by comparing the condition-types pairwise revealed the following. There was a simple effect of condition-type in the central, centro-parietal and parieto-occipital midline regions for the comparison ACP + NUM, and this effect was marginal in the parietal midline region. Comparing PER + NUM, there was a simple effect of condition-type in the centro-parietal and parieto-occipital midline regions, with this effect being marginal in the central midline region. Comparing ACP + GEN, there was simple effect of condition-type in the central, parietal and parieto-occipital midline regions, with this effect being marginal in the centro-parietal region. Comparing PER + GEN, there was a simple effect of condition-type in the parieto-occipital midline region, with this effect being marginal in central, centro-parietal and parietal midline regions. When the subject-type was plural, there was an effect of condition-type in the fronto-central, central and centro-parietal midline regions. Resolving this effect by comparing the condition-types pairwise revealed the following. There was a simple effect of condition-type in the fronto-central, central and centro-parietal midline regions for the comparison PER + GEN. For the comparison NUM + GEN, there was a marginal effect of condition-type in the central and centro-parietal midline regions.

**Table S1:** ANOVA: ERPs at the Verb : 300−500 ms

| Effects in time-window 300–500 ms |  |  |  | DF | Lateral Regions |  | DF | Midline Regions |  |
| --- | --- | --- | --- | --- | --- | --- | --- | --- | --- |
| ◆ MIDR x ST |  |  |  |  |  |  | 5,165 | 4.56 | ★ |
| ↳ MIDR = FCZ |  |  |  |  |  |  | 1,33 | 4.47 | ★ |
| ◆ ST x CT |  |  |  | 3,99 | 2.99 | ★ | 3,99 | 3.70 | ★★ |
| ↳ ST = S | ◆ CT |  |  | 3,99 | 3.02 | ★ |  |  |  |
|  | ↳ CT = PER+NUM | ◆ CT |  | 1,33 | 6.77 | ★ |  |  |  |
| ◆ MIDR x ST x CT |  |  |  |  |  |  | 15,495 | 3.03 | ★★ |
| ↳ MIDR = FCZ | ◆ ST x CT |  |  |  |  |  | 3,99 | 2.62 | ☆ |
|  | ↳ ST = P | ◆ CT |  |  |  |  | 3,99 | 2.59 | ☆ |
|  |  | ↳ CT = PER+GEN | ◆ CT |  |  |  | 1,33 | 5.96 | ★ |
| ↳ MIDR = CZ | ◆ ST x CT |  |  |  |  |  | 3,99 | 4.79 | ★★ |
|  | ↳ ST = S | ◆ CT |  |  |  |  | 3,99 | 3.89 | ★★ |
|  |  | ↳ CT = ACP+NUM | ◆ CT |  |  |  | 1,33 | 7.47 | ★ |
|  |  | ↳ CT = PER+NUM | ◆ CT |  |  |  | 1,33 | 4.88 | ☆ |
|  |  | ↳ CT = ACP+GEN | ◆ CT |  |  |  | 1,33 | 5.86 | ★ |
|  |  | ↳ CT = PER+GEN | ◆ CT |  |  |  | 1,33 | 5.11 | ☆ |
|  | ↳ ST = P | ◆ CT |  |  |  |  | 3,99 | 3.11 | ★ |
|  |  | ↳ CT = PER+GEN | ◆ CT |  |  |  | 1,33 | 8.46 | ★ |
|  |  | ↳ CT = NUM+GEN | ◆ CT |  |  |  | 1,33 | 4.97 | ☆ |
| ↳ MIDR = CPZ | ◆ ST x CT |  |  |  |  |  | 3,99 | 4.27 | ★★ |
|  | ↳ ST = S | ◆ CT |  |  |  |  | 3,99 | 4.38 | ★★ |
|  |  | ↳ CT = ACP+NUM | ◆ CT |  |  |  | 1,33 | 6.59 | ★ |
|  |  | ↳ CT = PER+NUM | ◆ CT |  |  |  | 1,33 | 7.38 | ★ |
|  |  | ↳ CT = ACP+GEN | ◆ CT |  |  |  | 1,33 | 4.78 | ☆ |
|  |  | ↳ CT = PER+GEN | ◆ CT |  |  |  | 1,33 | 5.35 | ☆ |
|  | ↳ ST = P | ◆ CT |  |  |  |  | 3,99 | 2.43 | ☆ |
|  |  | ↳ CT = PER+GEN | ◆ CT |  |  |  | 1,33 | 5.85 | ★ |
|  |  | ↳ CT = NUM+GEN | ◆ CT |  |  |  | 1,33 | 5.41 | ☆ |
| ↳ MIDR = PZ | ◆ ST x CT |  |  |  |  |  | 3,99 | 4.39 | ★★ |
|  | ↳ ST = S | ◆ CT |  |  |  |  | 3,99 | 3.92 | ★★ |
|  |  | ↳ CT = ACP+NUM | ◆ CT |  |  |  | 1,33 | 5.11 | ☆ |
|  |  | ↳ CT = ACP+GEN | ◆ CT |  |  |  | 1,33 | 7.04 | ★ |
|  |  | ↳ CT = PER+GEN | ◆ CT |  |  |  | 1,33 | 5.23 | ☆ |

...Table S1 continued
| Effects in time-window 300–500 ms |  |  |  | DF | Lateral Regions | DF | Midline Regions |  |
| --- | --- | --- | --- | --- | --- | --- | --- | --- |
| LMIDR = | POZ | ◆ ST x CT |  |  |  | 3,99 | 3.72 | ★★ |
|  |  | ↳ST = S | ◆ CT |  |  | 3,99 | 5.89 | ★★★ |
|  |  |  | ↳CT = ACP+NUM | ◆ CT |  | 1,33 | 8.94 | ★★ |
|  |  |  | ↳CT = PER+NUM | ◆ CT |  | 1,33 | 6.09 | ★ |
|  |  |  | ↳CT = ACP+GEN | ◆ CT |  | 1,33 | 10.51 | ★★ |
|  |  |  | ↳CT = PER+GEN | ◆ CT |  | 1,33 | 7.52 | ★ |

##### S1.2.2 Alternative filter settings

The EEG data collected during the experiment was pre-processed for analysis using a 0.3−20 Hz bandpass filter offline in order to remove slow signal drifts that might lead to stimulus-independent differences between conditions. As discussed in the EEG preprocessing section of the main article, the filter settings we employed effectively circumvents distortion issues without the need for a baseline correction, and are sufficiently broad to include language-related ERP activity that is typically in the frequency range of about 0.5 to 5 Hz (Roehm et al., 2002). Previous studies employing identical filter settings to ours report no significant differences in results between applying this bandpass filter versus applying baseline corrections (see Burkhardt et al., 2007; Choudhary et al., 2009). Nevertheless, in order to verify that this is the case with our data as well, we reanalysed the data post-hoc using two slightly different bandpass filter settings: by increasing the low-pass cut-off frequency alone (0.3 −30 Hz) or by decreasing the high-pass cut-off frequency in addition to increasing the low-pass cut-off (0.1−30 Hz).

Figure S1 shows the ERPs of the entire sentence epoch for the singular subject conditions and for the filter settings we employed for the analysis of our data, namely a 0.3−20 Hz bandpass filter. This figure illustrates that the effects of interest at the critical position in our study, namely the verb (the third N1-P2 complex in the figure starting after about 1.5 seconds into the trial), do not appear to dependent upon the ERPs before the pre-critical positions. Further, no significant ERP differences are apparent in the pre-critical positions, perhaps as an expected consequence of our fully counter-balanced design and identical materials preceding the verb in a given subject-type.

Figure S2 shows the ERPs of the entire sentence epoch for the singular subject conditions from the analysis using the 0.3−30 Hz bandpass filter. ERPs using this filter setting remain very much comparable to those we report. Figure S3 shows the ERPs of the entire sentence epoch for the singular subject conditions from the analysis using the 0.1−30 Hz bandpass filter. As an expected consequence of the lower high-pass cutoff in this setting, slow signal drifts were apparent within the sentence epoch, which however could be mostly rectified to some extent using a pre-critical baseline correction (−200 to 0 ms), with some of the differences originating starting from the sentence onset still remaining. Nevertheless, the overall pattern of ERPs at the critical position remained similar to the other two analyses. Differences in magnitude and presence or absence of some effects (which are to be expected in a comparison of this sort) notwithstanding, Table S2 provides an overview of this confirmatory analysis, and highlights the crucial parts of this analysis, which show a consistent pattern that obtains in all of the filter settings. These observations are further supported by a running t-test of significance on the ERP plots comparing the two different filter settings, as shown in Figure S4 and Figure S5 respectively.

**Figure S1:**
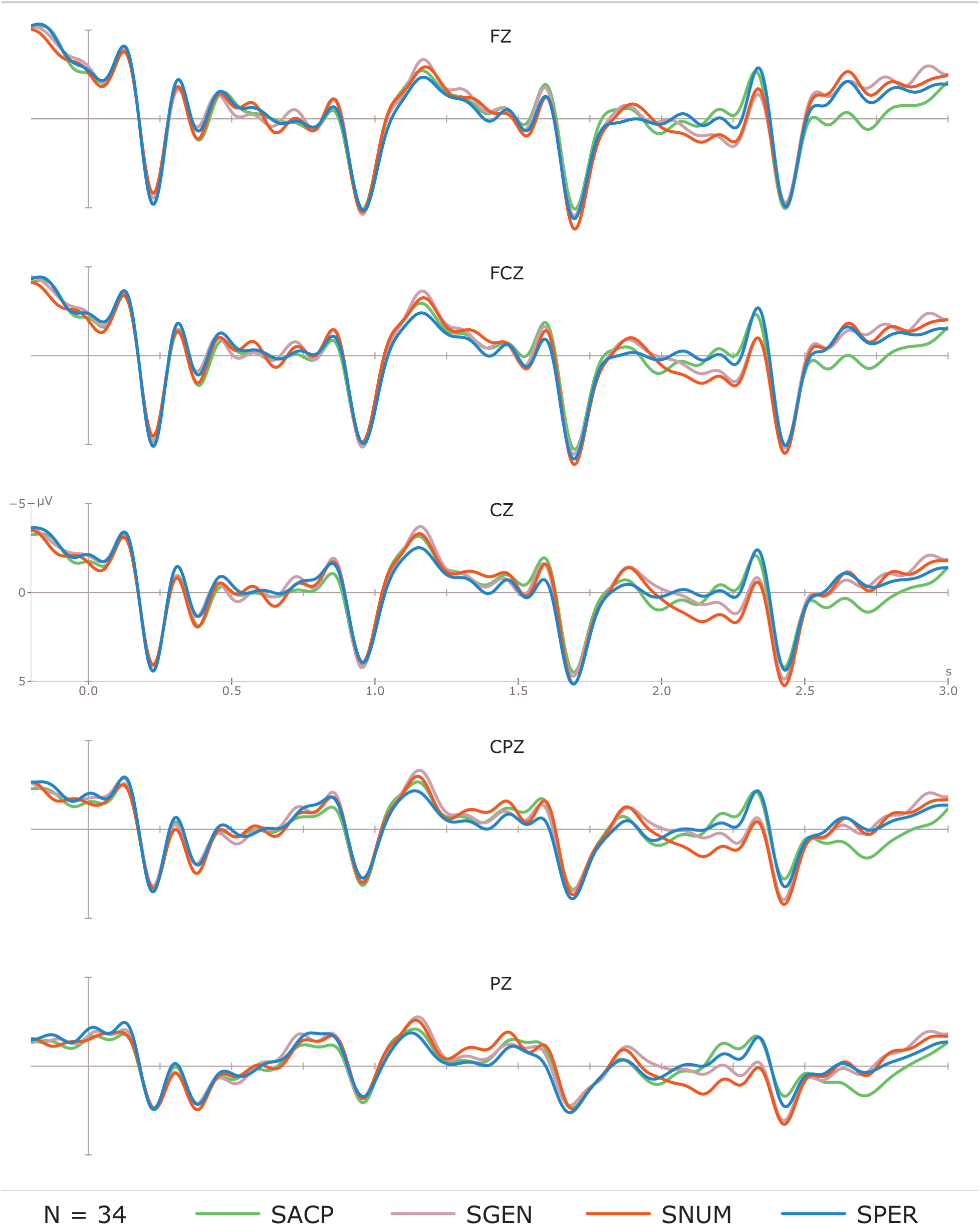
ERPs of the entire sentence epoch: Singular subject conditions. 0.3−20 Hz BPF.

**Figure S2:**
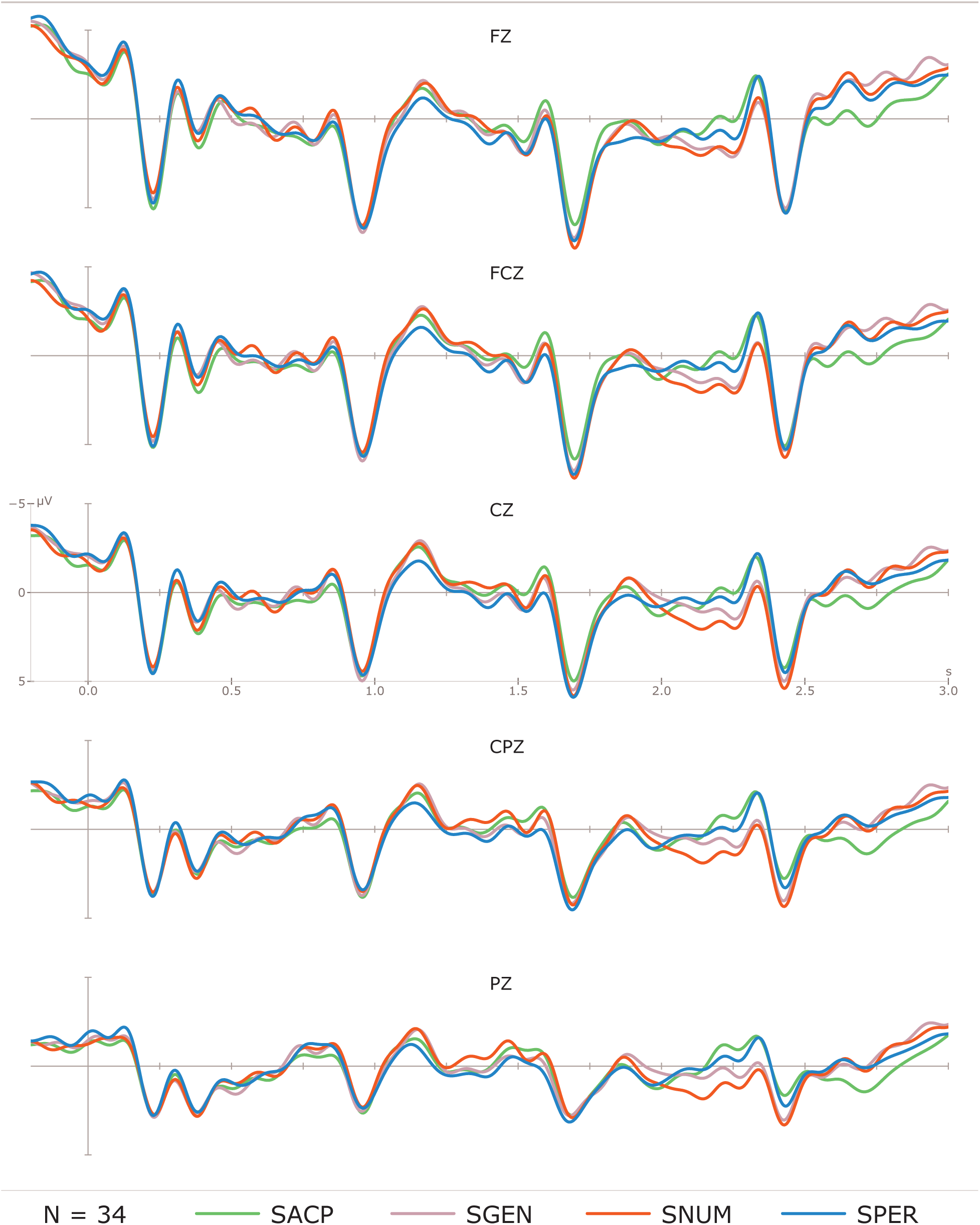
ERPs of the entire sentence epoch: Singular subject conditions. 0.3−30 Hz BPF.

**Figure S3:**
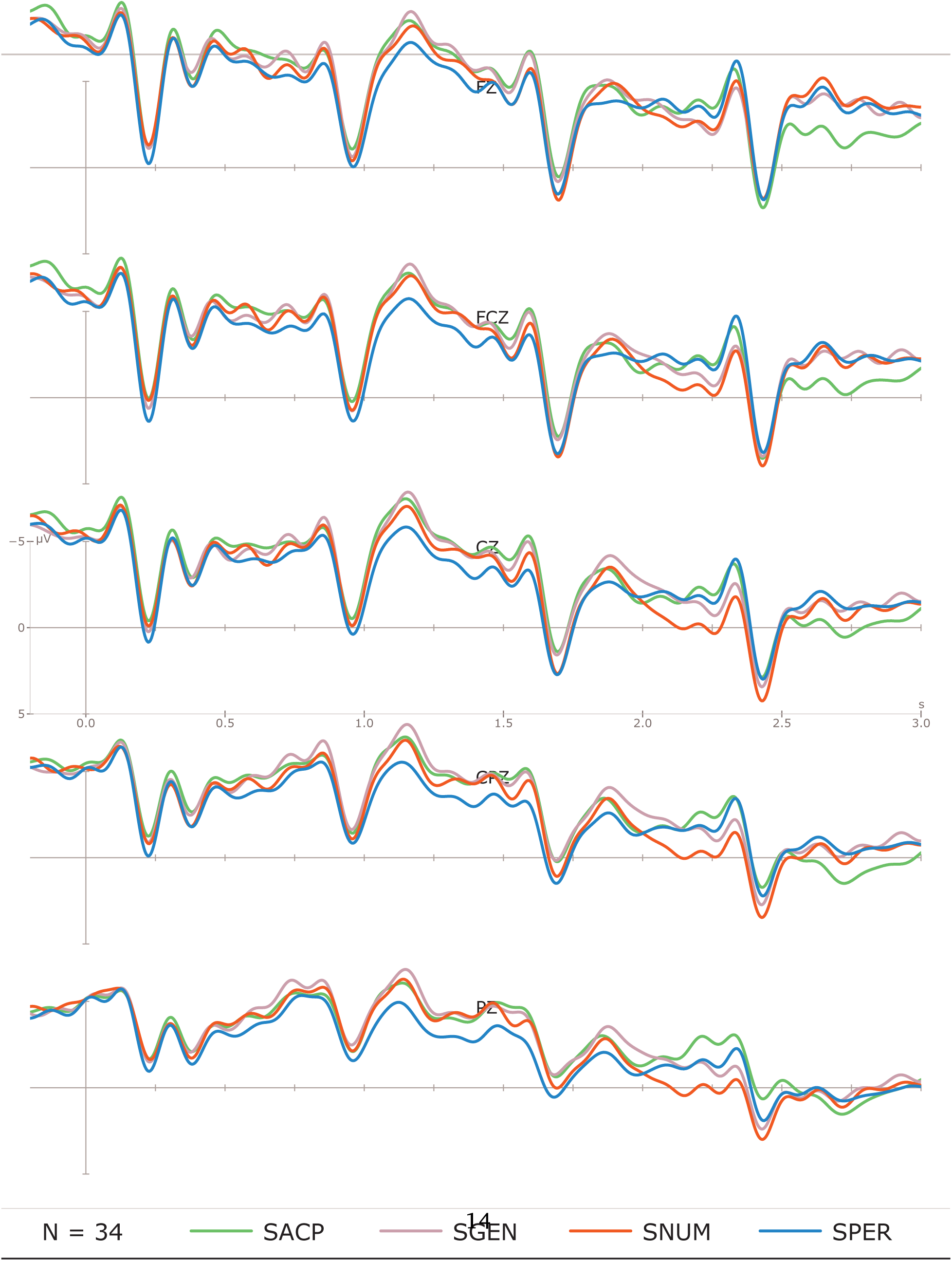
ERPs of the entire sentence epoch: Singular subject conditions. 0.1−30 Hz BPF.

**Figure S4:**
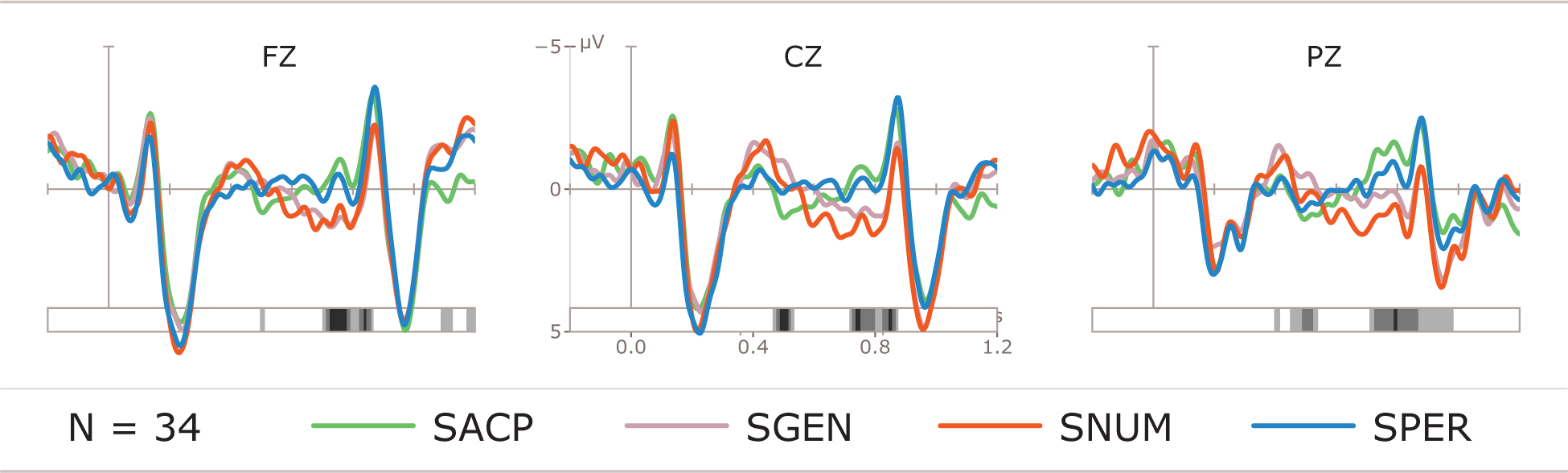
ERPs at the verb with running t-test to detect potentially significant time-windows: Singular subject conditions. 0.3−20 Hz BPF.

**Figure S5:**
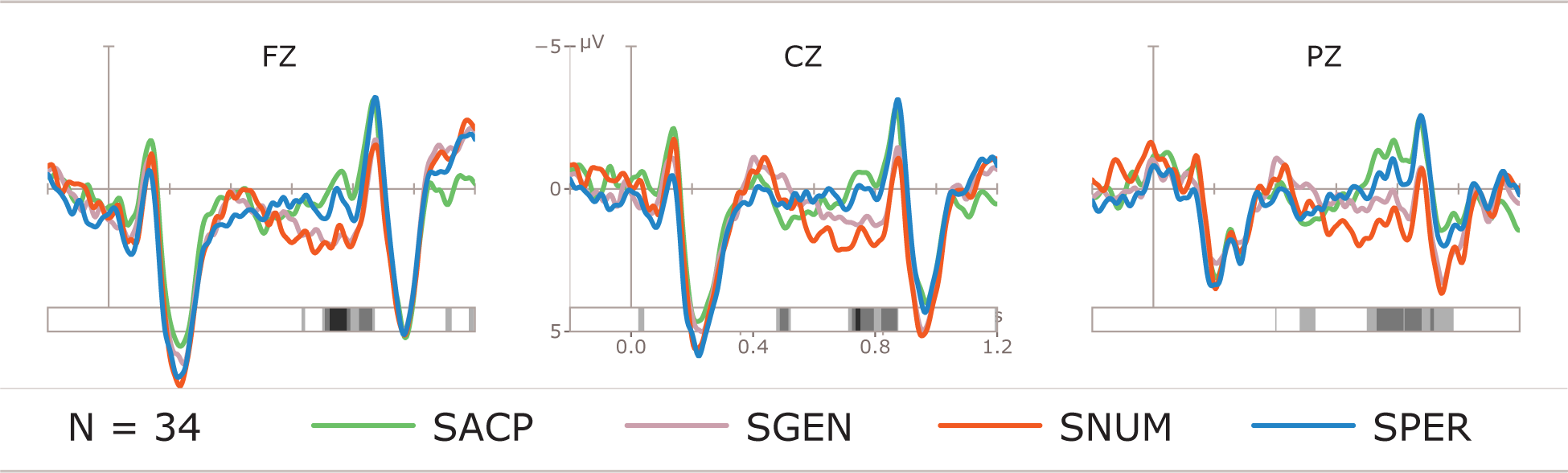
ERPs at the verb with running t-test to detect potentially significant time-windows: Singular subject conditions. 0.3−30 Hz BPF.

**Table S2:**
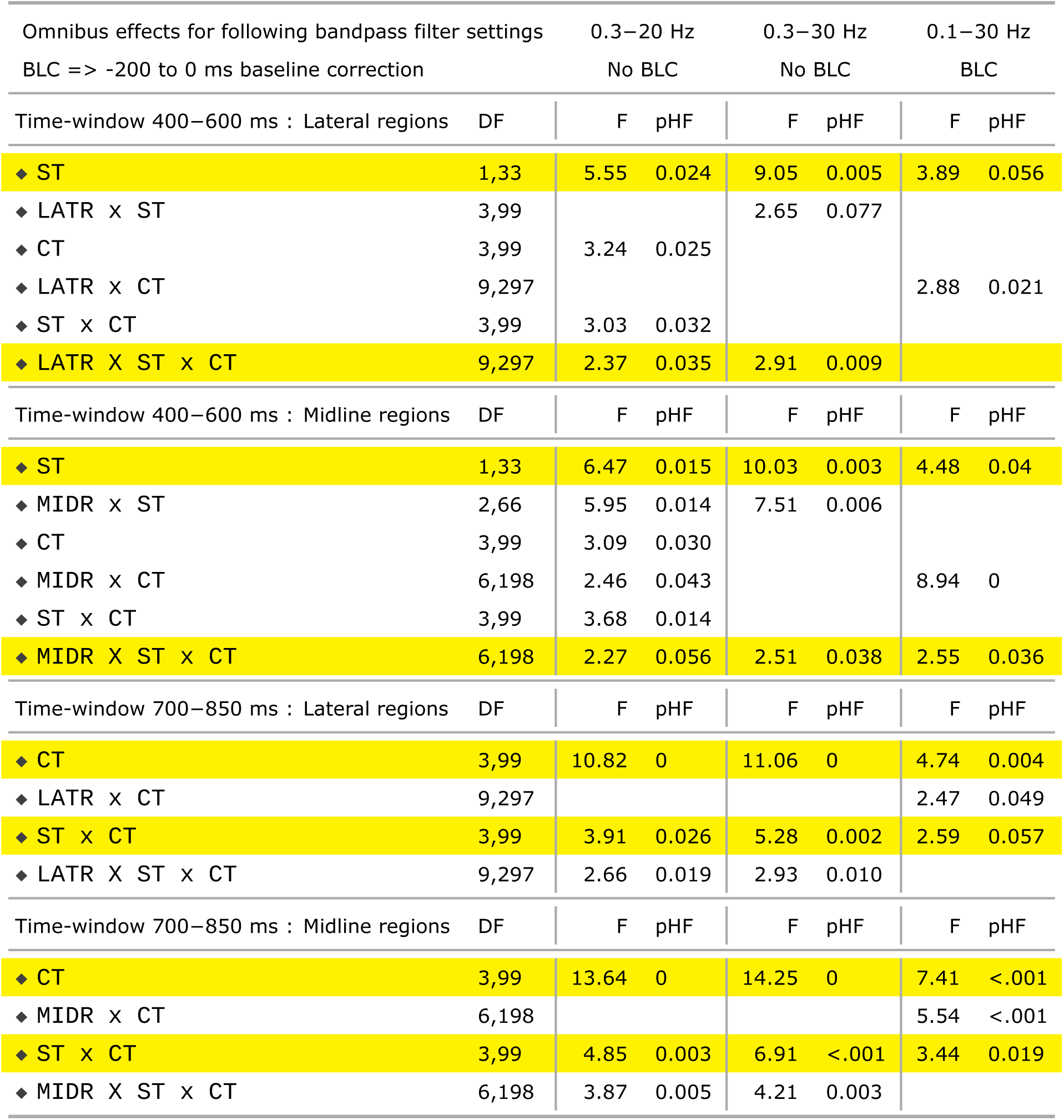
ANOVA: ERPs at the Verb. Comparable effects for three different filter settings. The main article presents results based on data analysis using a 0.3−20 Hz bandpass filter, which circumvents the need for a pre-stimulus baseline correction.

| Omnibus effects for following bandpass filter settings<br>BLC => -200 to 0 ms baseline correction |  | 0.3–20 Hz<br>No BLC |  | 0.3–30 Hz<br>No BLC |  | 0.1–30 Hz<br>BLC |  |
| --- | --- | --- | --- | --- | --- | --- | --- |
| Time-window 400–600 ms : Lateral regions | DF | F | pHF | F | pHF | F | pHF |
| ◆ ST | 1,33 | 5.55 | 0.024 | 9.05 | 0.005 | 3.89 | 0.056 |
| ◆ LATR x ST | 3,99 |  |  | 2.65 | 0.077 |  |  |
| ◆ CT | 3,99 | 3.24 | 0.025 |  |  |  |  |
| ◆ LATR x CT | 9,297 |  |  |  |  | 2.88 | 0.021 |
| ◆ ST x CT | 3,99 | 3.03 | 0.032 |  |  |  |  |
| ◆ LATR X ST x CT | 9,297 | 2.37 | 0.035 | 2.91 | 0.009 |  |  |
| Time-window 400–600 ms : Midline regions | DF | F | pHF | F | pHF | F | pHF |
| ◆ ST | 1,33 | 6.47 | 0.015 | 10.03 | 0.003 | 4.48 | 0.04 |
| ◆ MIDR x ST | 2,66 | 5.95 | 0.014 | 7.51 | 0.006 |  |  |
| ◆ CT | 3,99 | 3.09 | 0.030 |  |  |  |  |
| ◆ MIDR x CT | 6,198 | 2.46 | 0.043 |  |  | 8.94 | 0 |
| ◆ ST x CT | 3,99 | 3.68 | 0.014 |  |  |  |  |
| ◆ MIDR X ST x CT | 6,198 | 2.27 | 0.056 | 2.51 | 0.038 | 2.55 | 0.036 |
| Time-window 700–850 ms : Lateral regions | DF | F | pHF | F | pHF | F | pHF |
| ◆ CT | 3,99 | 10.82 | 0 | 11.06 | 0 | 4.74 | 0.004 |
| ◆ LATR x CT | 9,297 |  |  |  |  | 2.47 | 0.049 |
| ◆ ST x CT | 3,99 | 3.91 | 0.026 | 5.28 | 0.002 | 2.59 | 0.057 |
| ◆ LATR X ST x CT | 9,297 | 2.66 | 0.019 | 2.93 | 0.010 |  |  |
| Time-window 700–850 ms : Midline regions | DF | F | pHF | F | pHF | F | pHF |
| ◆ CT | 3,99 | 13.64 | 0 | 14.25 | 0 | 7.41 | <.001 |
| ◆ MIDR x CT | 6,198 |  |  |  |  | 5.54 | <.001 |
| ◆ ST x CT | 3,99 | 4.85 | 0.003 | 6.91 | <.001 | 3.44 | 0.019 |
| ◆ MIDR X ST x CT | 6,198 | 3.87 | 0.005 | 4.21 | 0.003 |  |  |

#### S1.3 Critical Items

##### S1.3.1 Items Distribution Scheme

The distribution of critical items into five sets was carried out as follows: out of the 120 critical items in each of the 8 critical conditions, half contained masculine and the other half contained feminine nouns as subjects. Thus 60 critical nouns in each condition were masculine and 60 were feminine. To distribute these items in a fully counterbalanced manner, the masculine and feminine items are distributed separately. In each condition, 18 sentences containing masculine nouns and 18 sentences containing femining nouns were selected for each of the five sets of stimuli. Therefore, in each of the 8 conditions, 36 sentences were selected in total per stimulus set. This ensured that, within each given set, all conditions are equiprobable, all items occurred equal number of times (albeit in different conditions), all participants saw all items (albeit in different conditions), and there were equal number of sentences in each of the 8 critical conditions, half of them masculine and half feminine.

Thus each pariticipant saw a total of 36 sentences in each of the 8 critical conditions, 18 of which contained a masculine noun, and 18 contained a feminine nouns. The 18 sentences for a given subject (noun) gender from each condition containing 60 sentences in that gender were selected in two stages (first 12, and then 6), again to ensure equiprobability of items in all respects. This was also in order to ensure that, across the five sets, each stimulus sentence is equifrequent across the sets. This is illustrated in Figure S6 schematically. One of the five lists was used for each participant. The presentation of the lists was counterbalanced across participants.

The pointed and unpointed frequency information for the critical nouns and verbs used in the stimuli were extracted from Aralex lexical database (Boudelaa & Marslen-Wilson, 2010) whenever available (because this information was not available for a small number of cases). This information is provided in the following pages. Then, the critical sentences used in the study are listed with their item numbers and condition codes. In the interest of space, only translations are provided and not the full phonological and morphological gloss, and only a list of the 240 acceptable sentences (60 x 2 (singular, plural) x 2 (masculine, feminine)) and not their violation counterparts.

**Figure S6:**
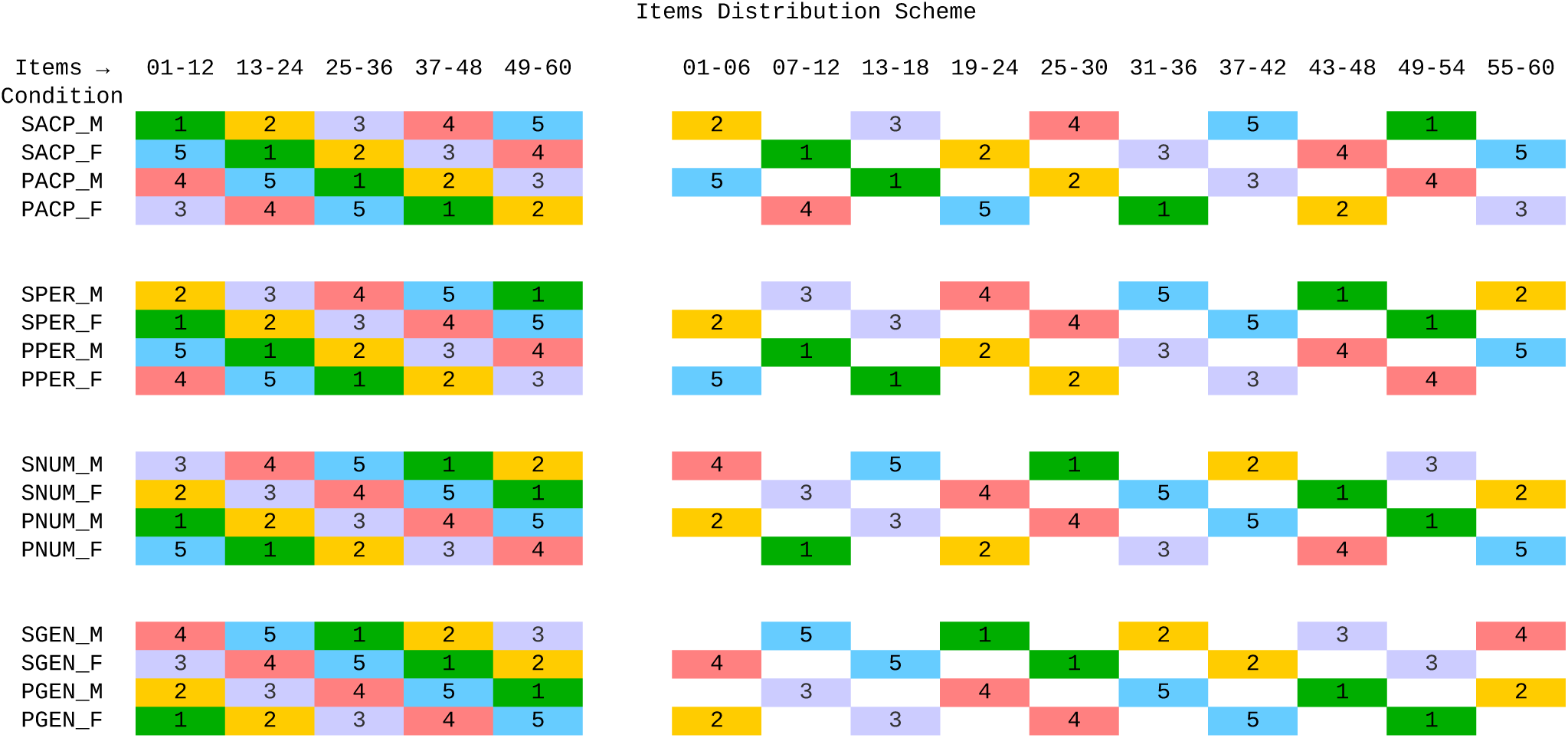
Items Distribution Scheme

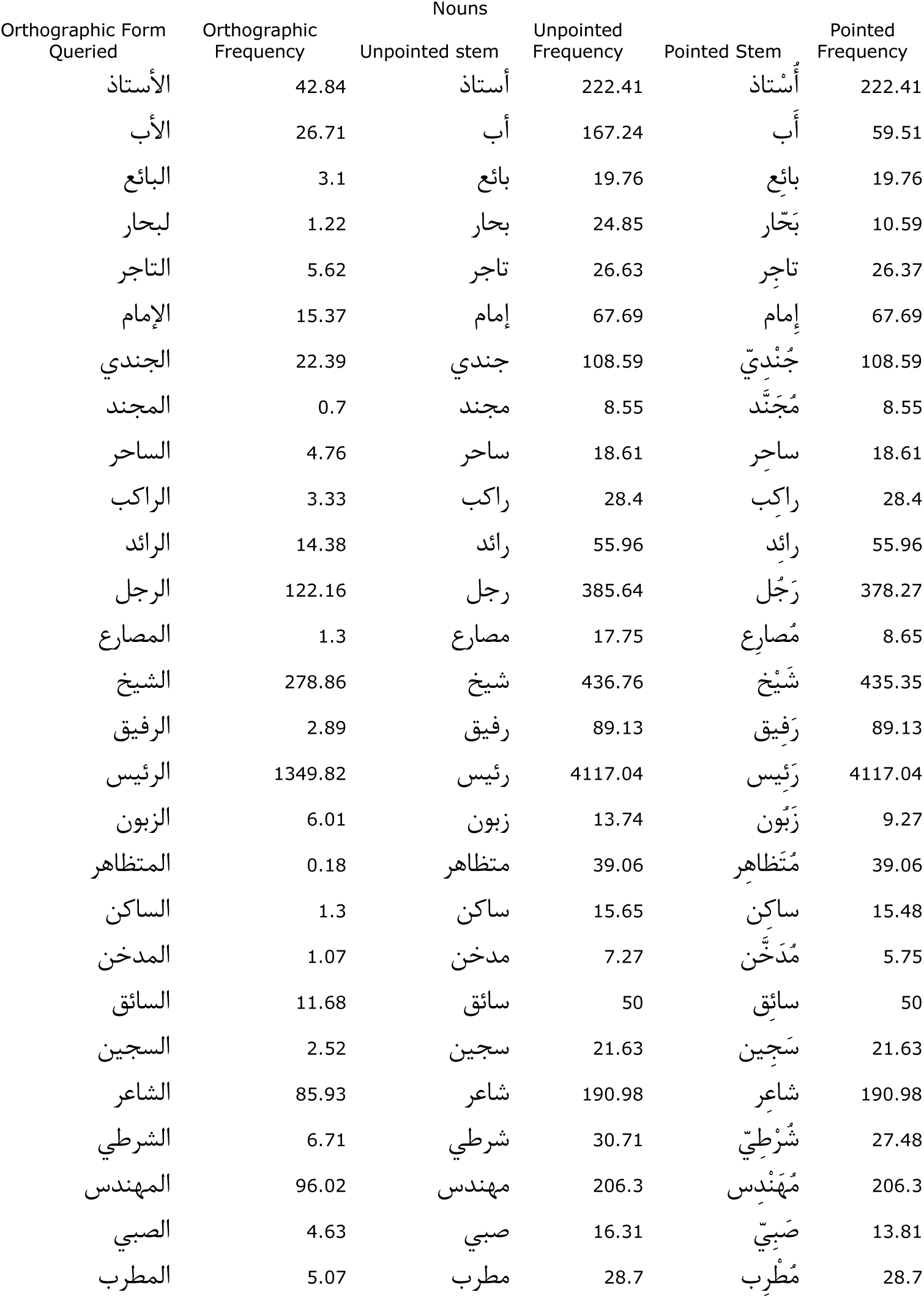
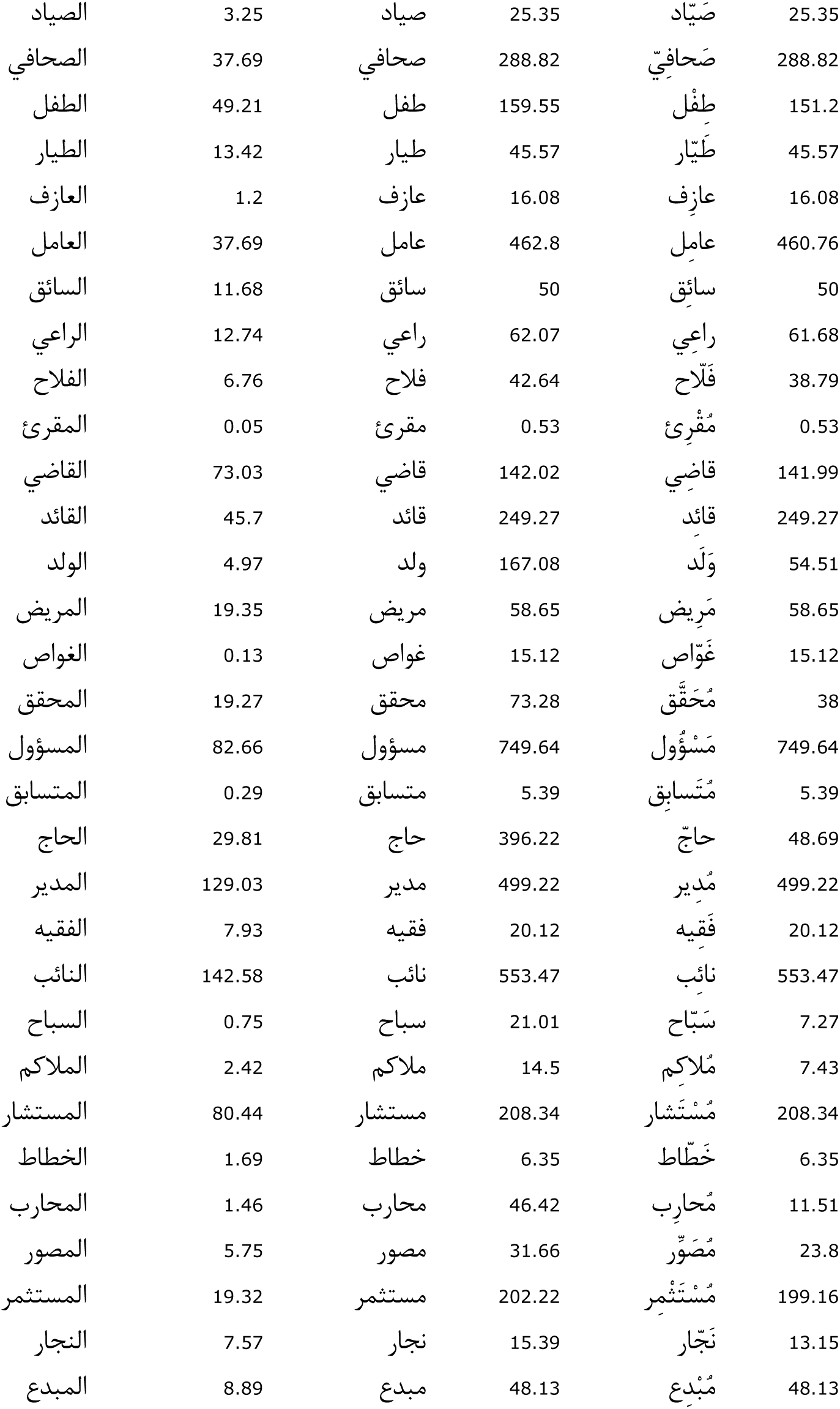
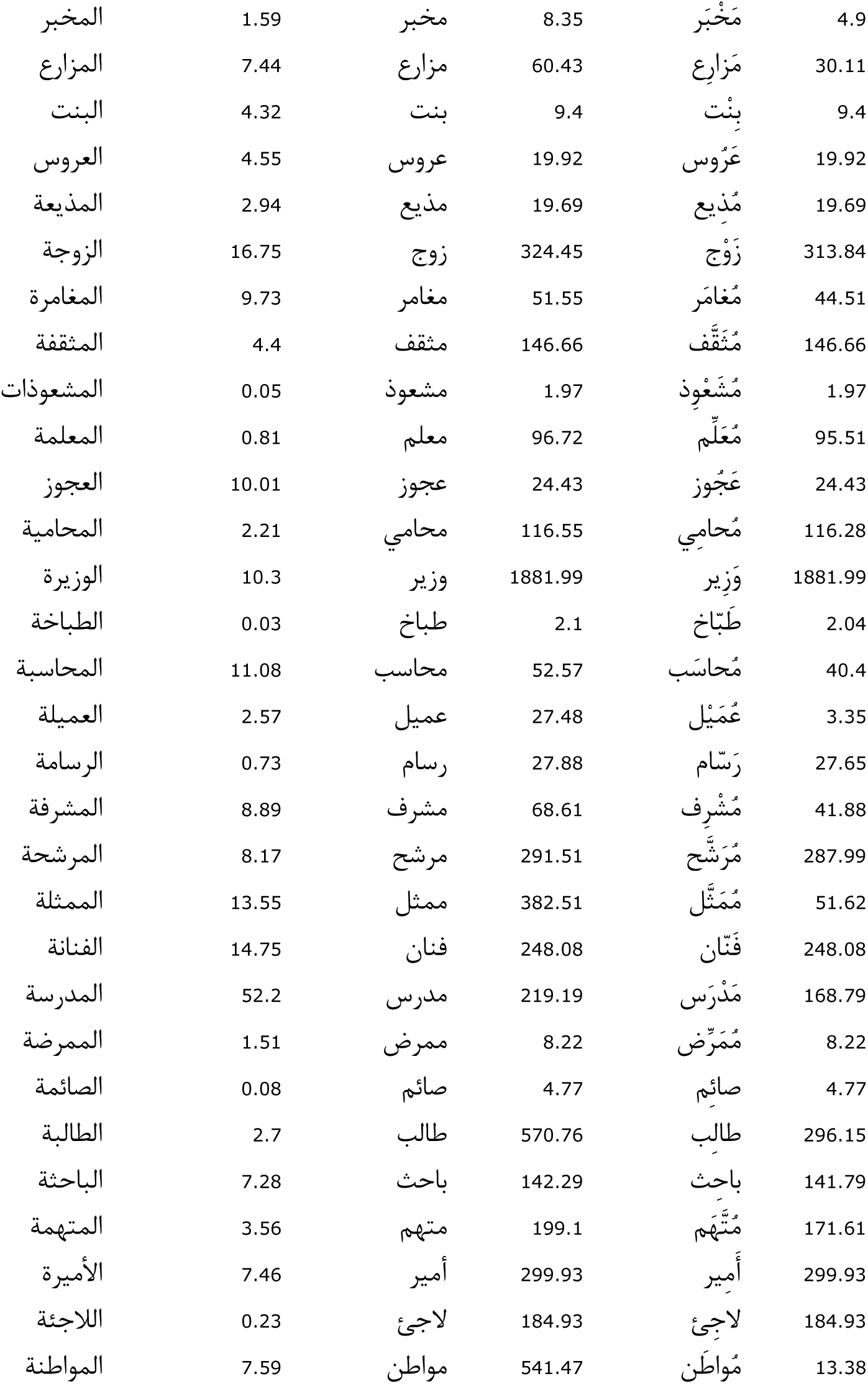
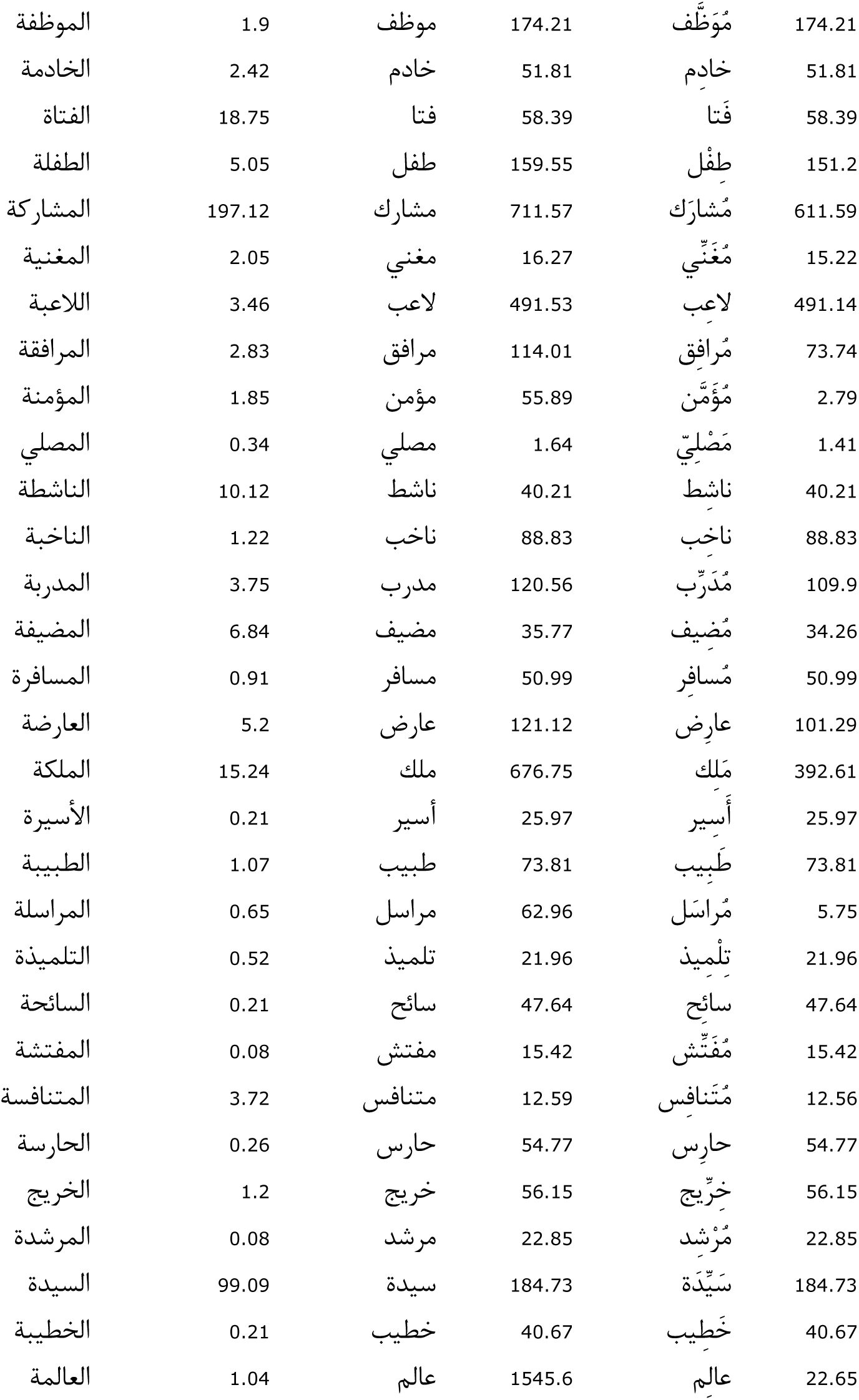
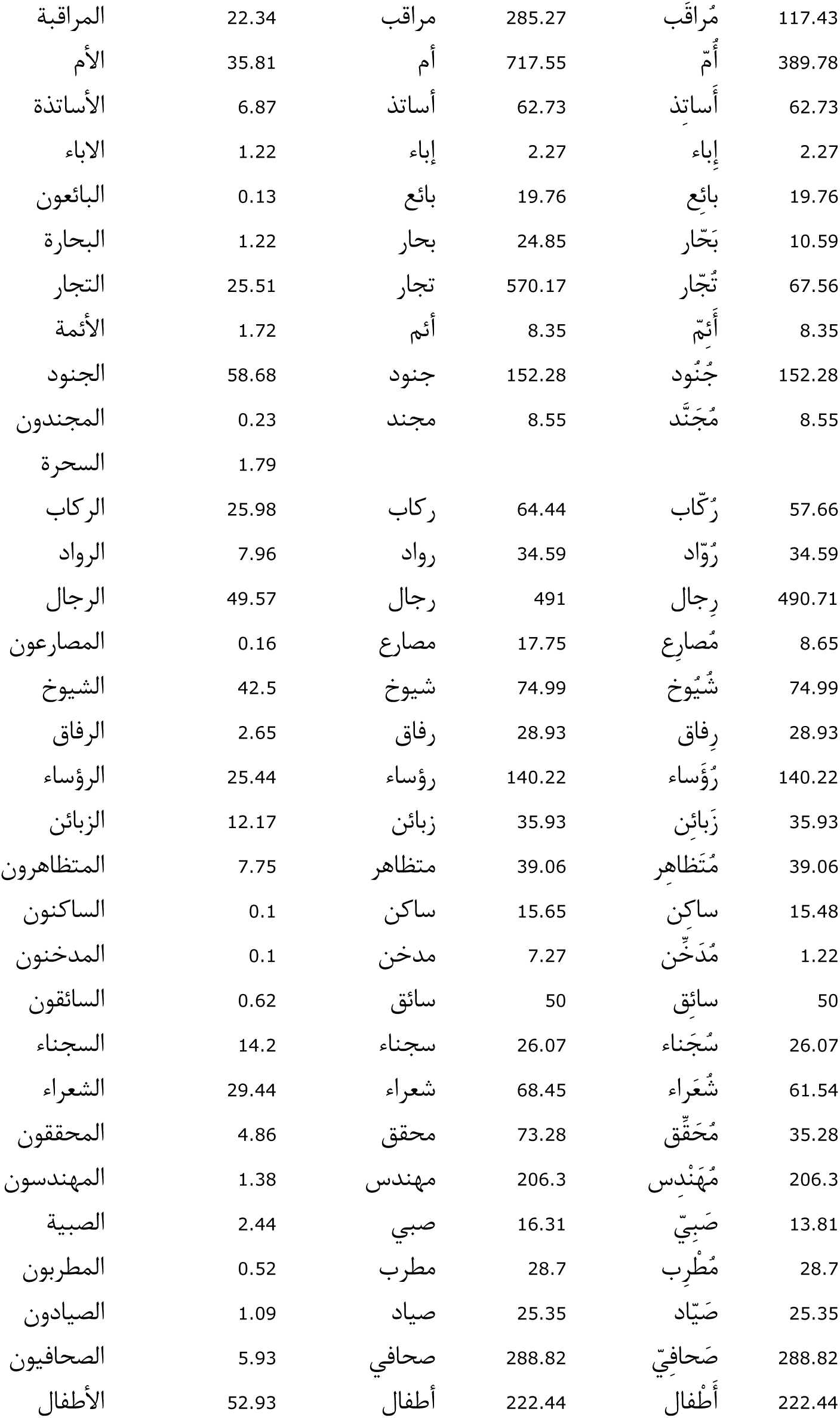
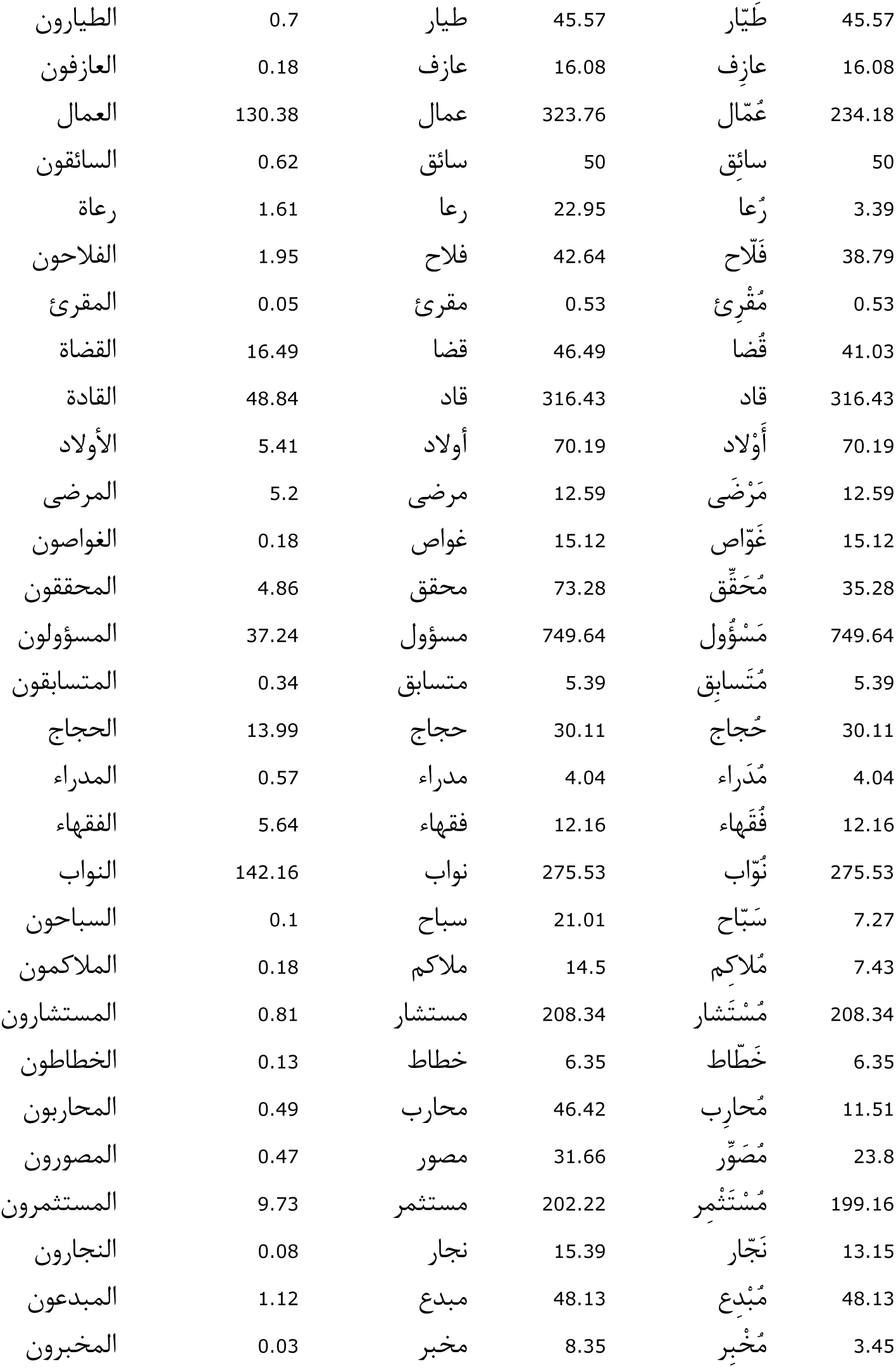
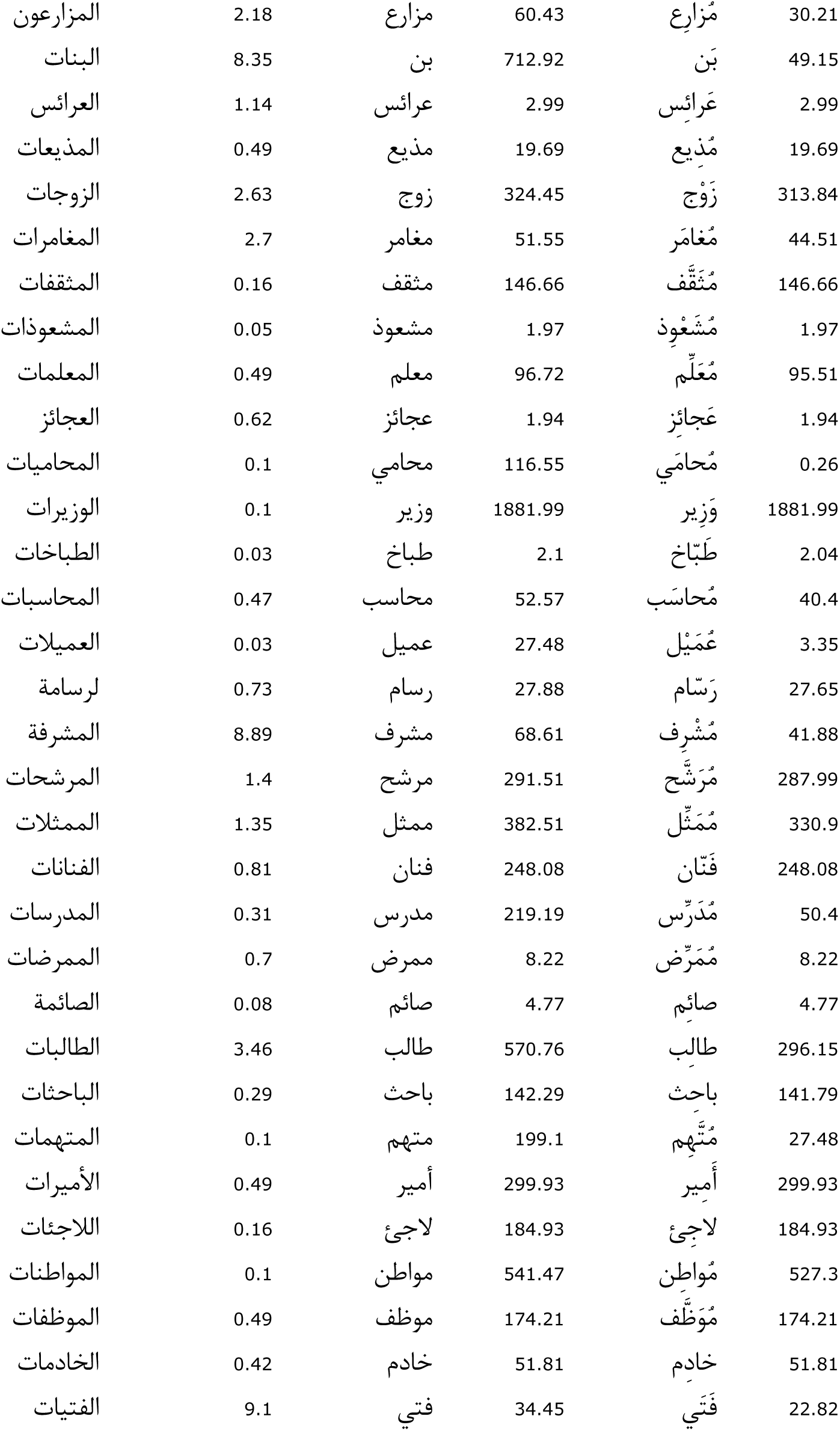
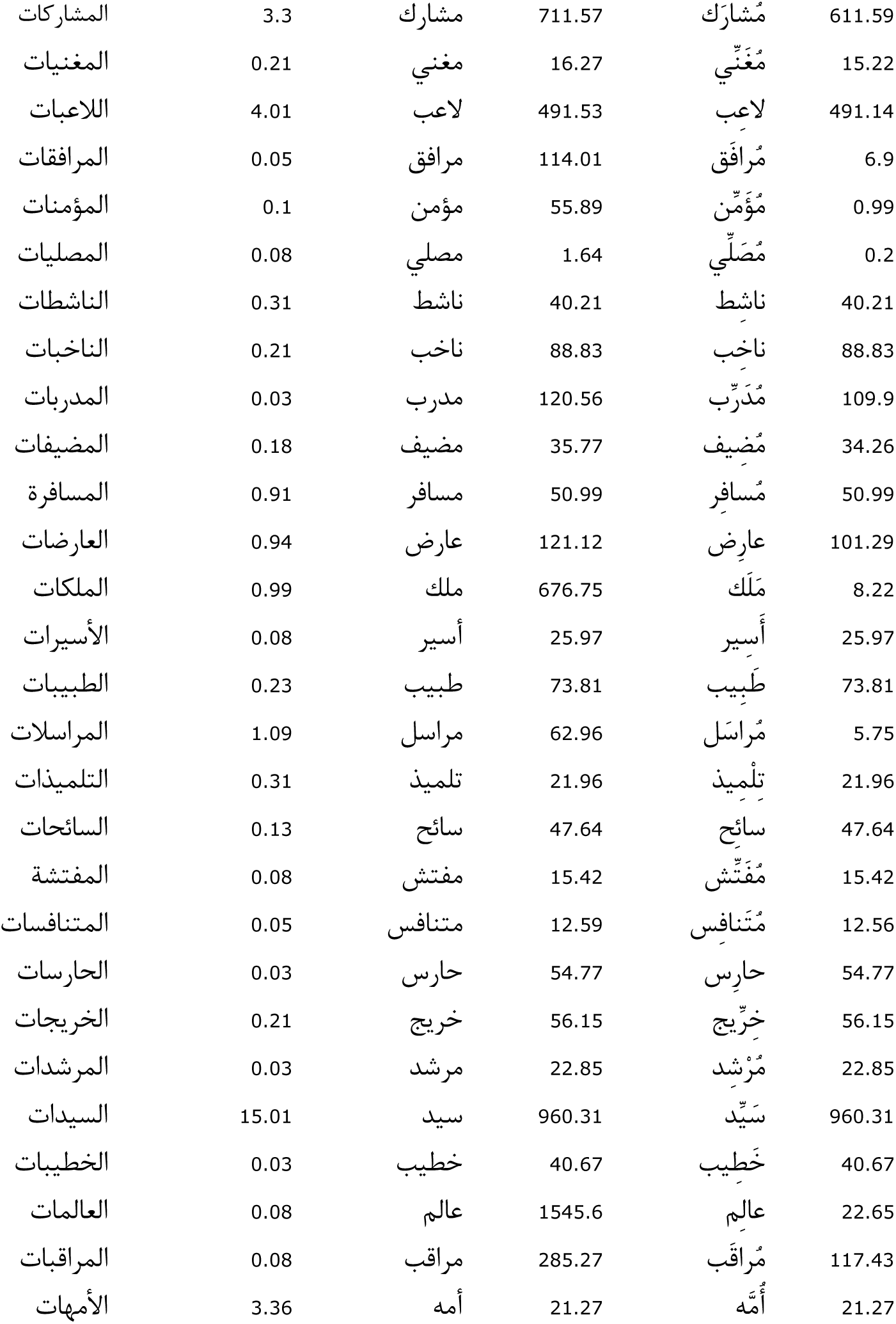

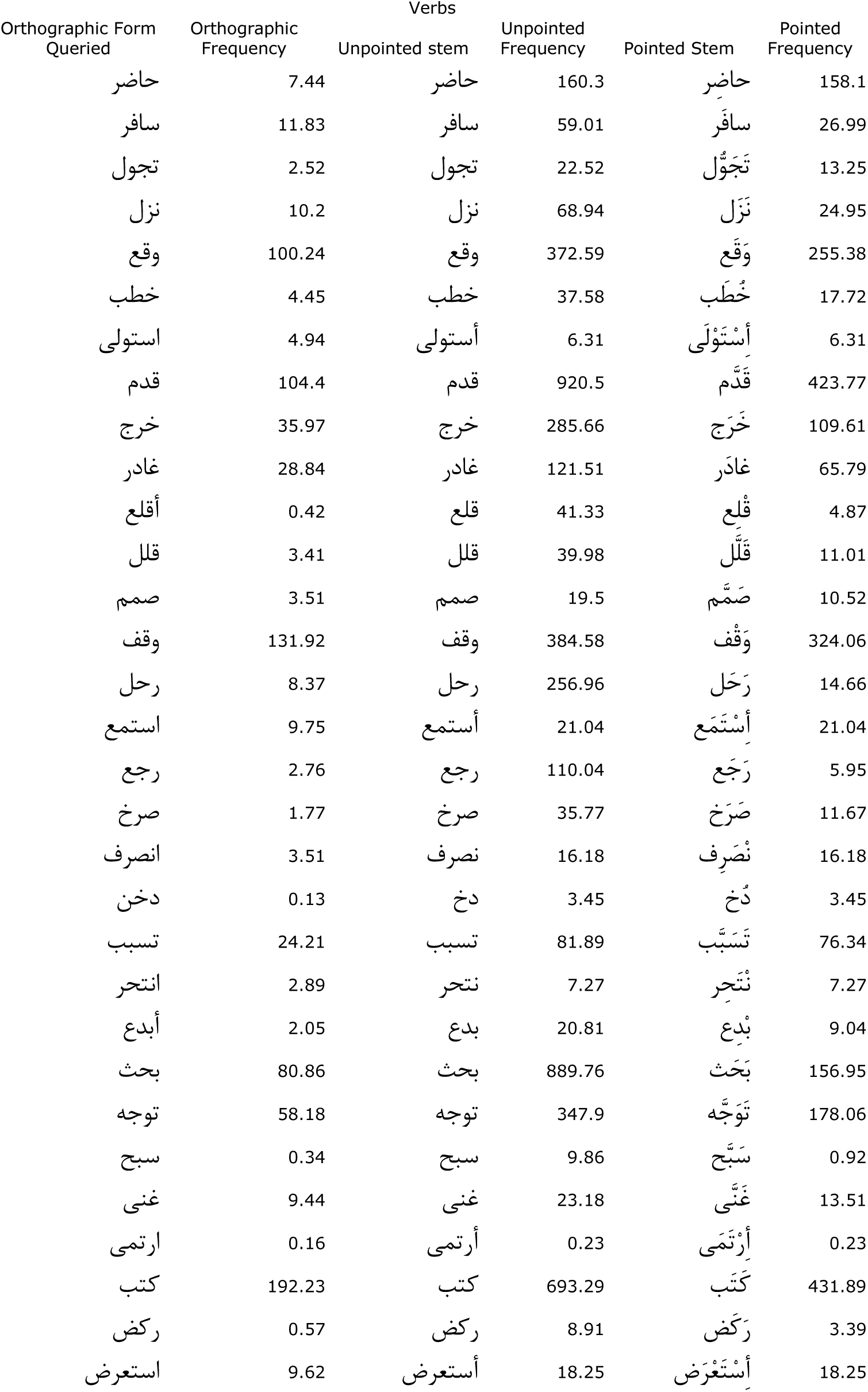
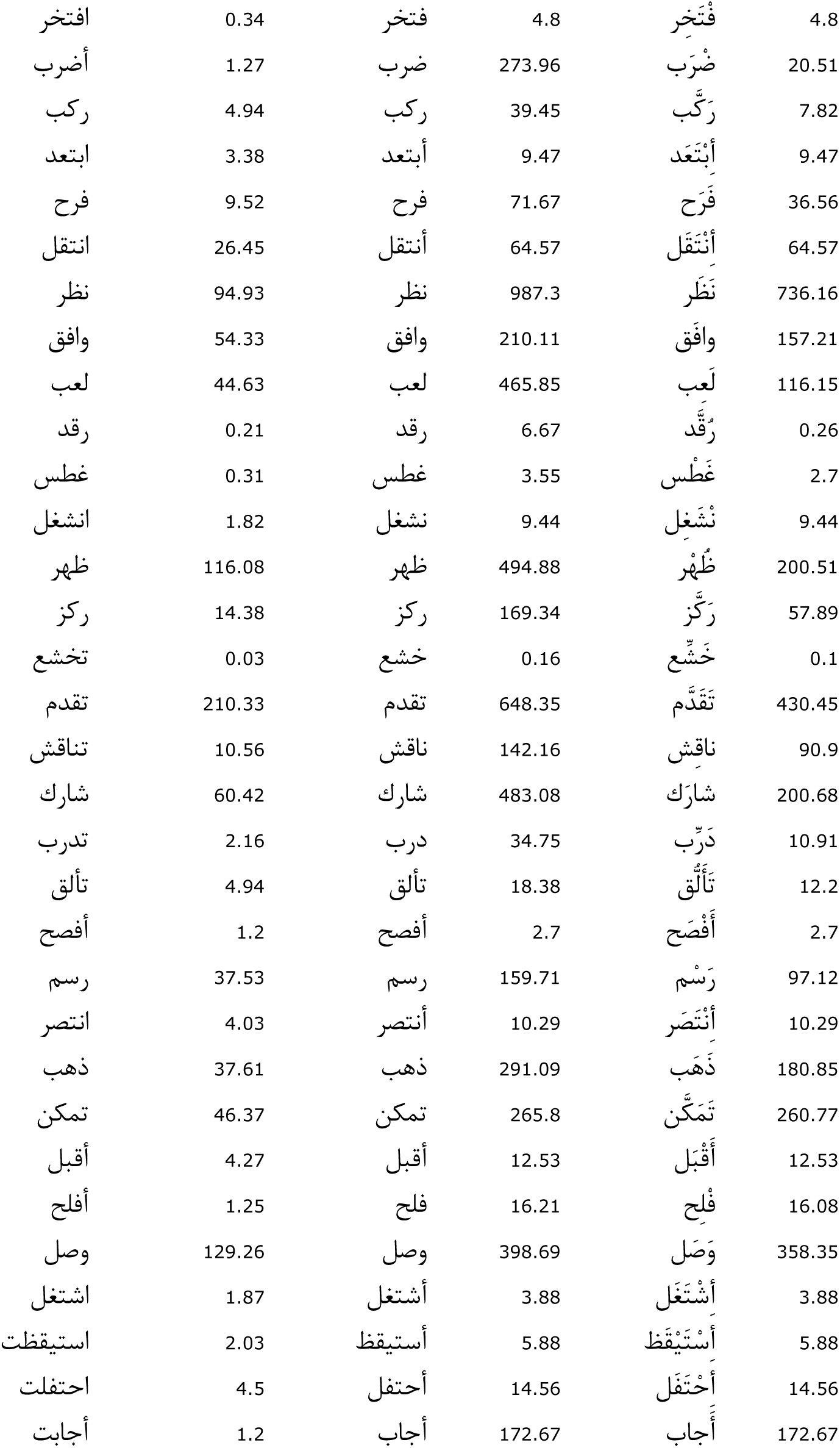
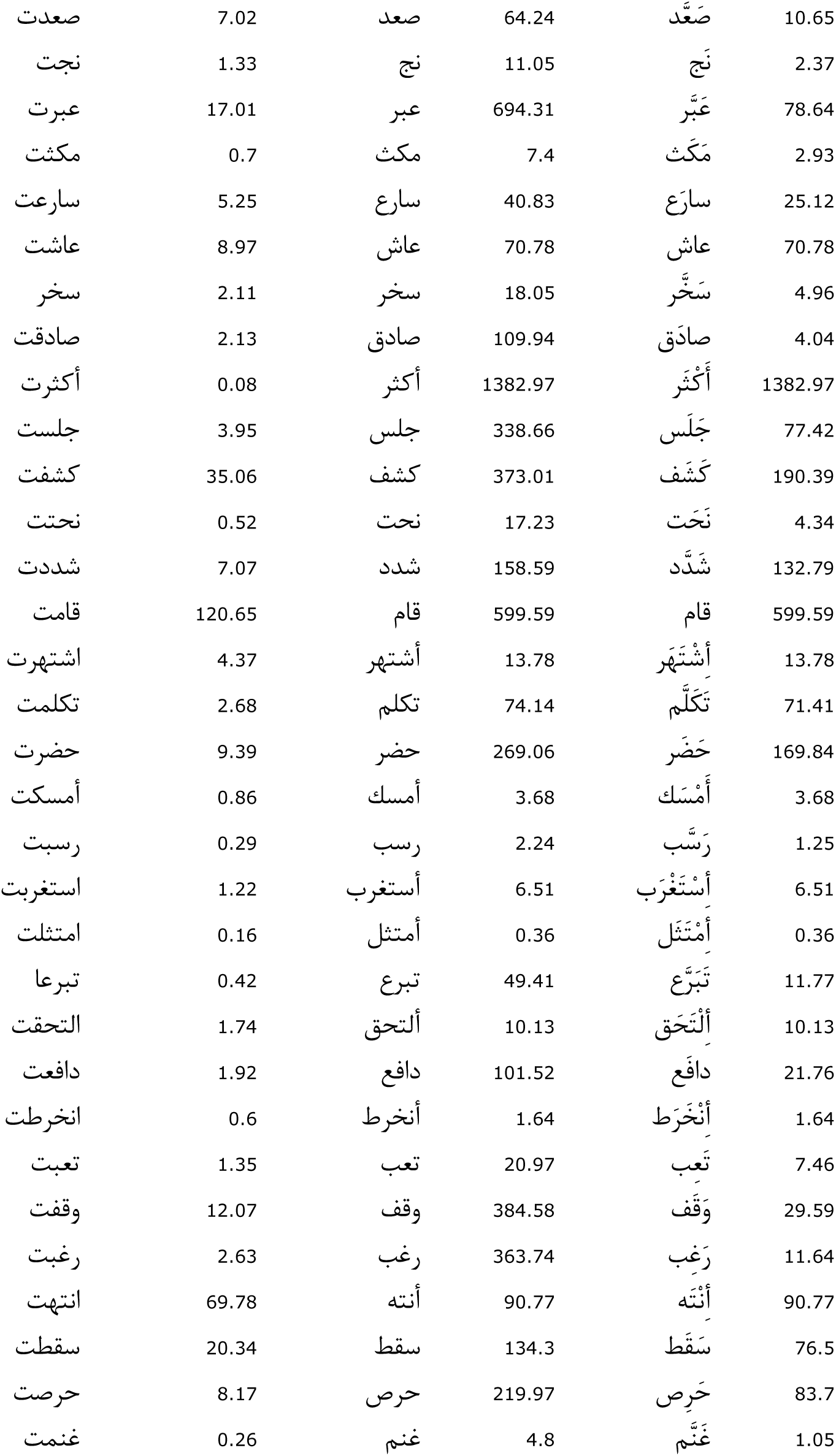
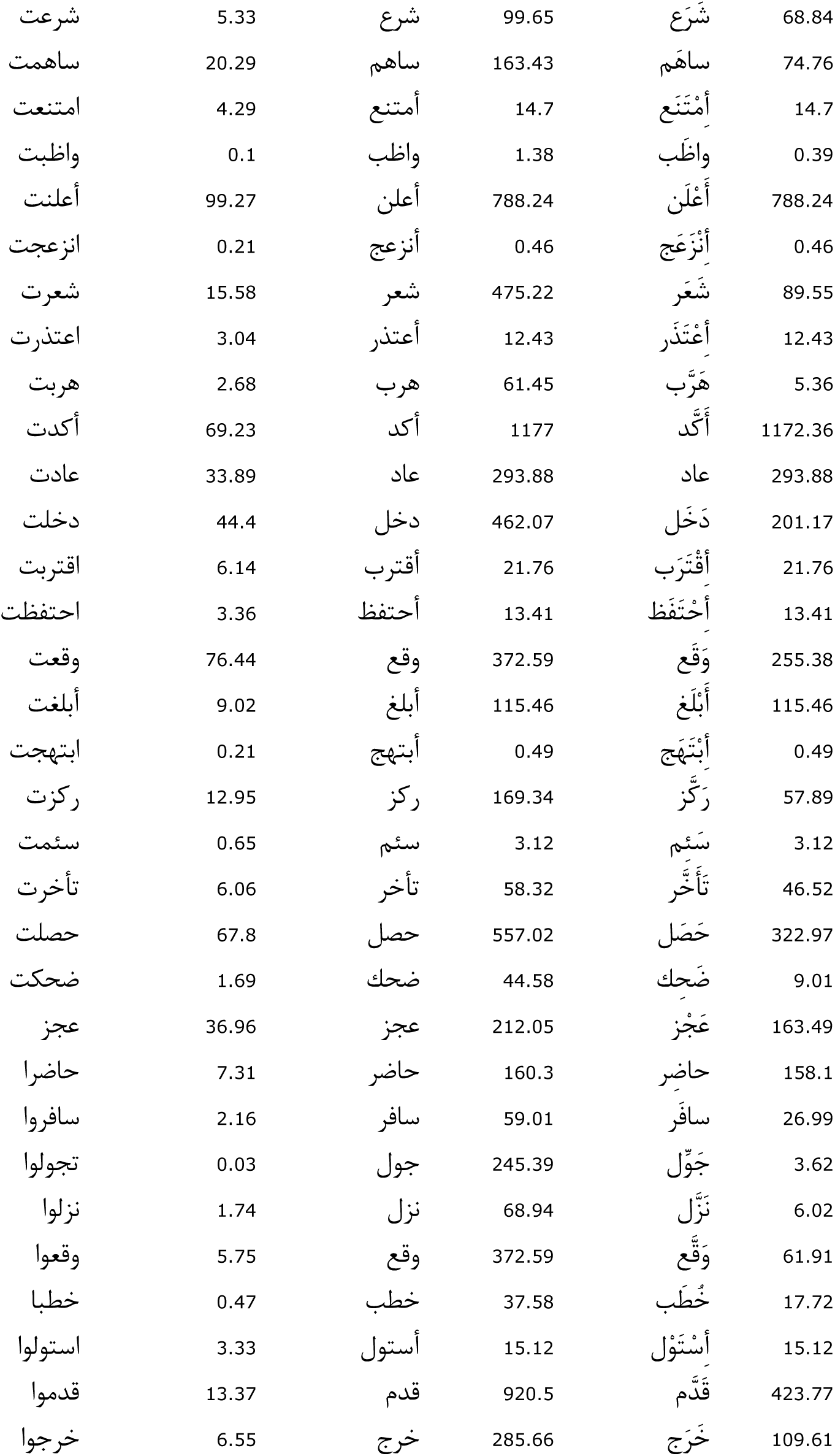
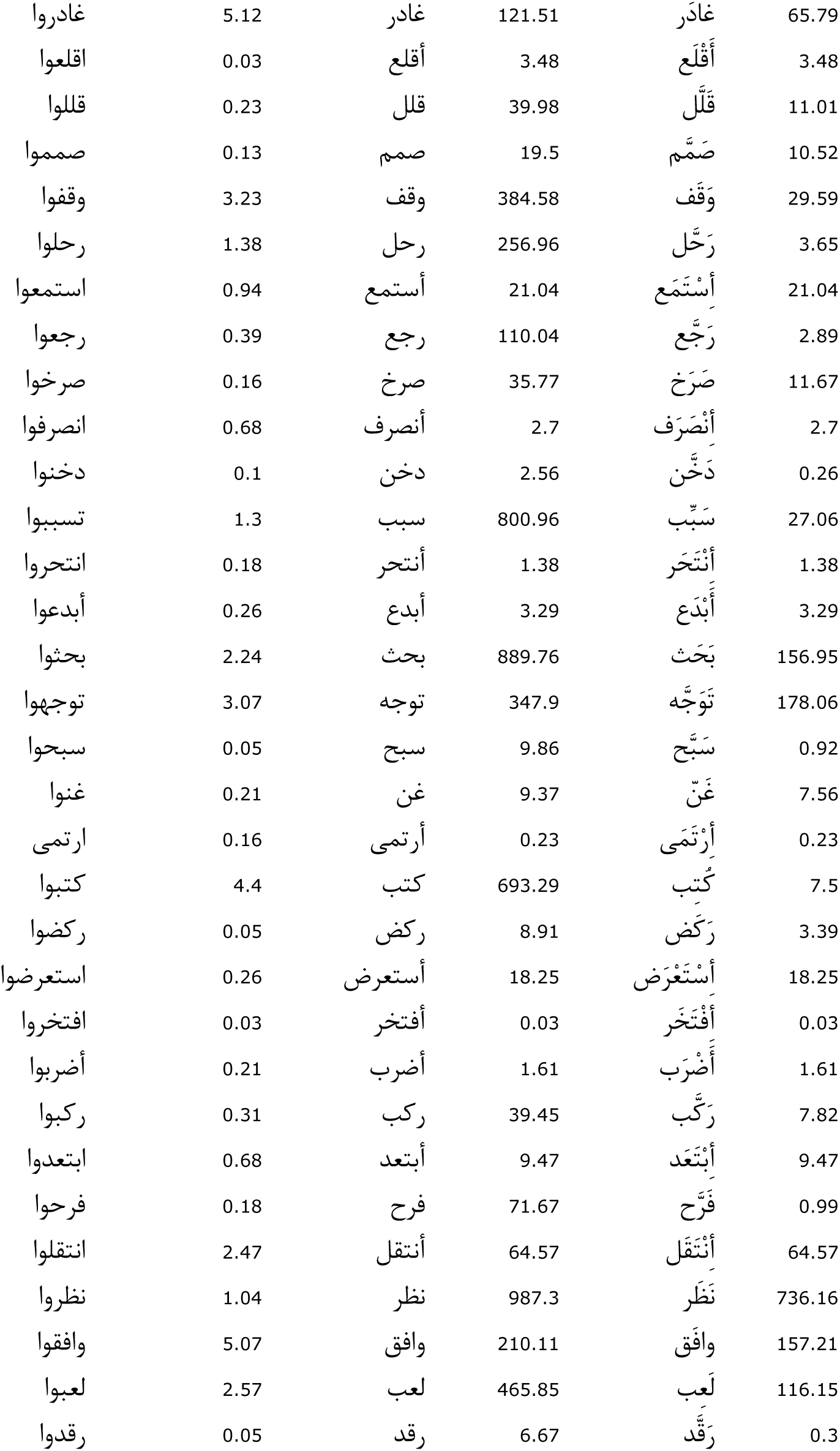
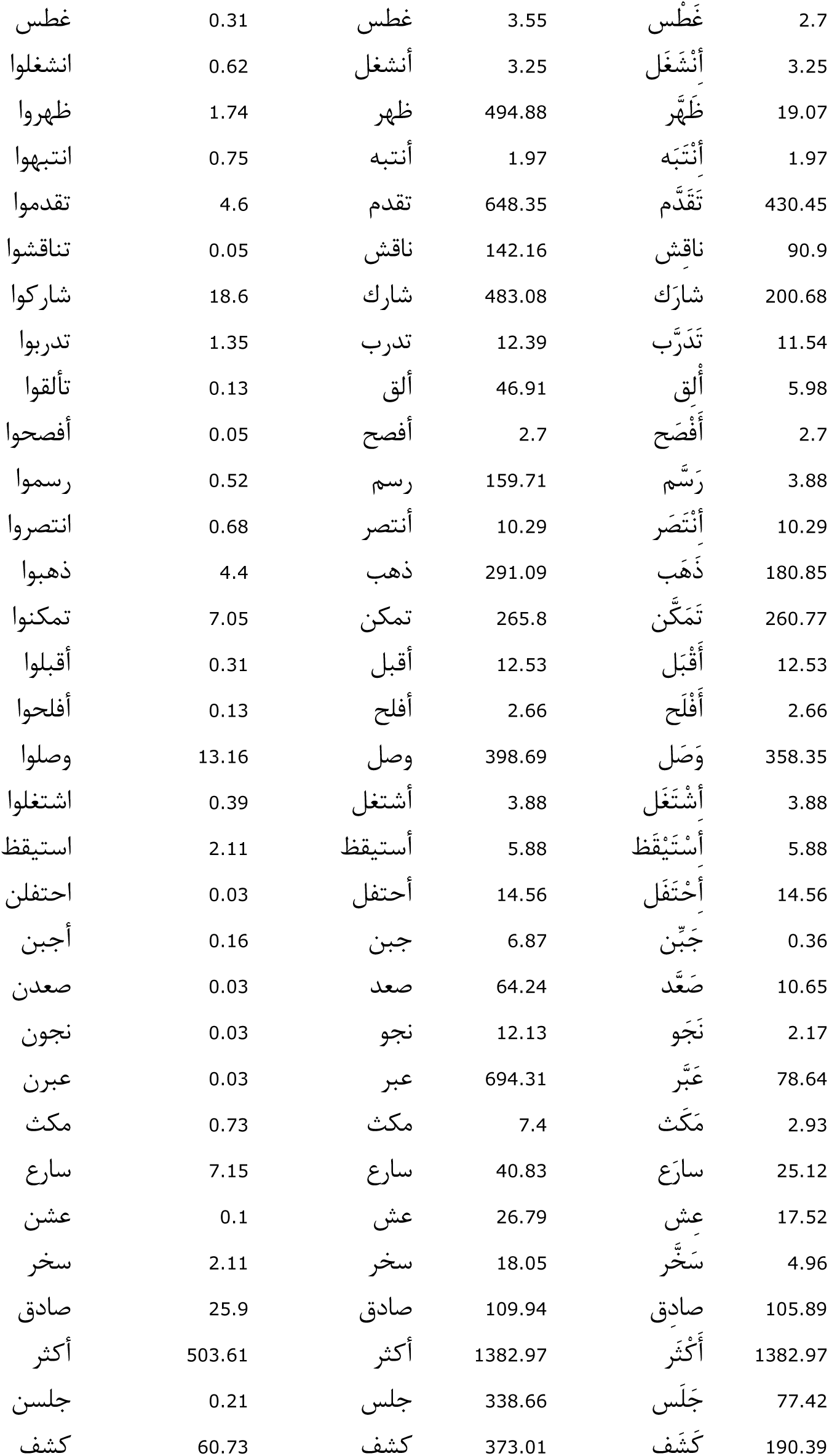
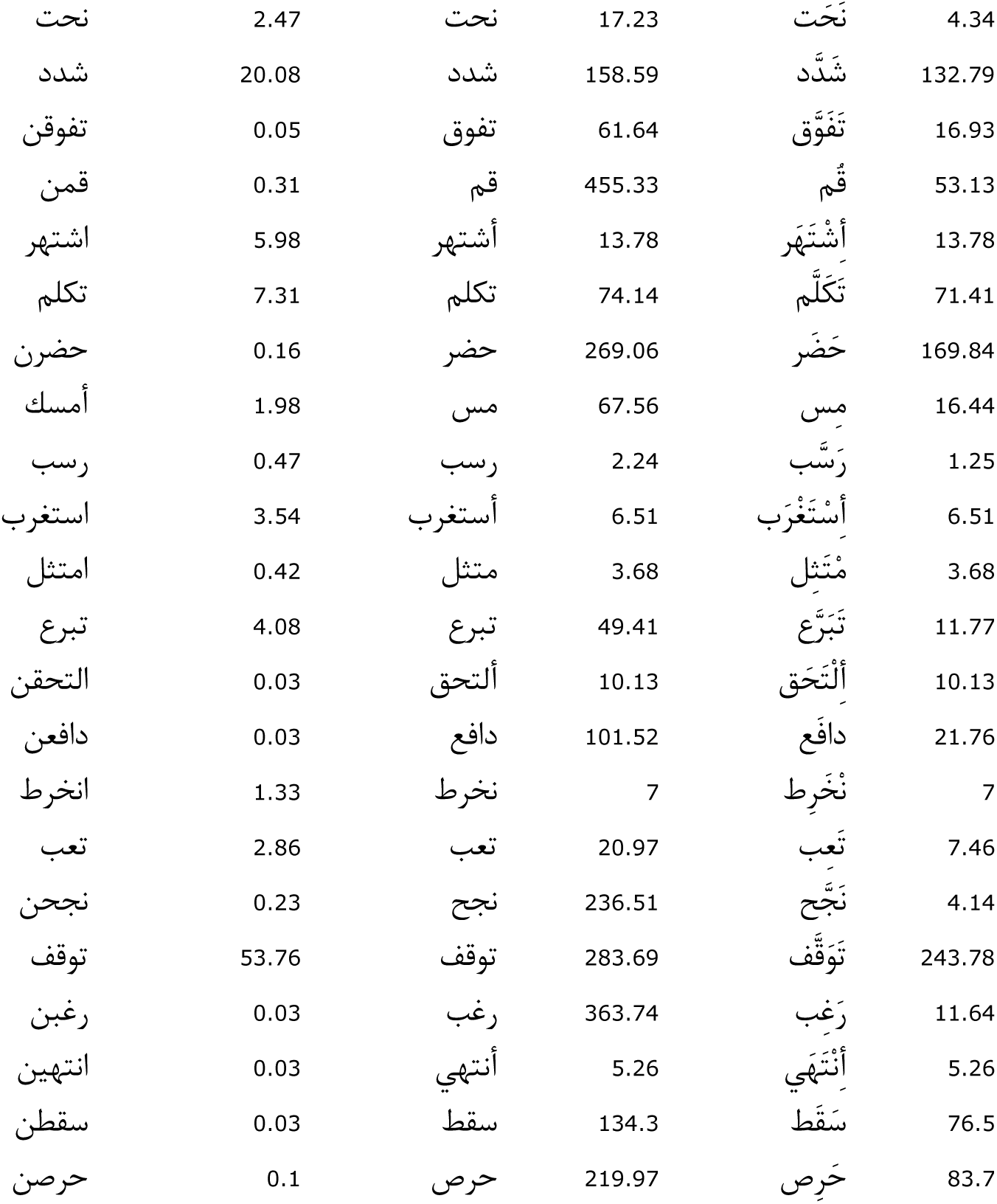
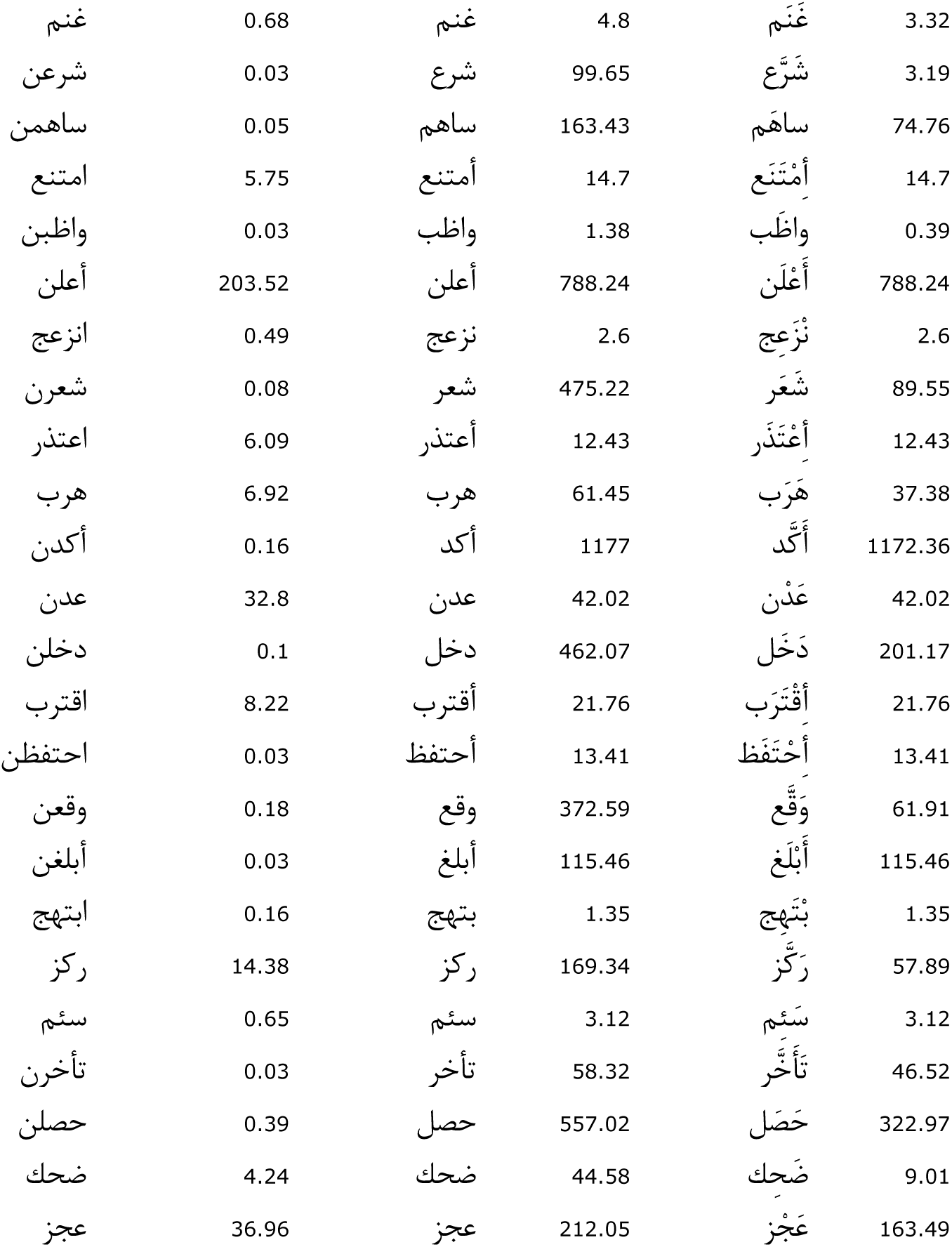

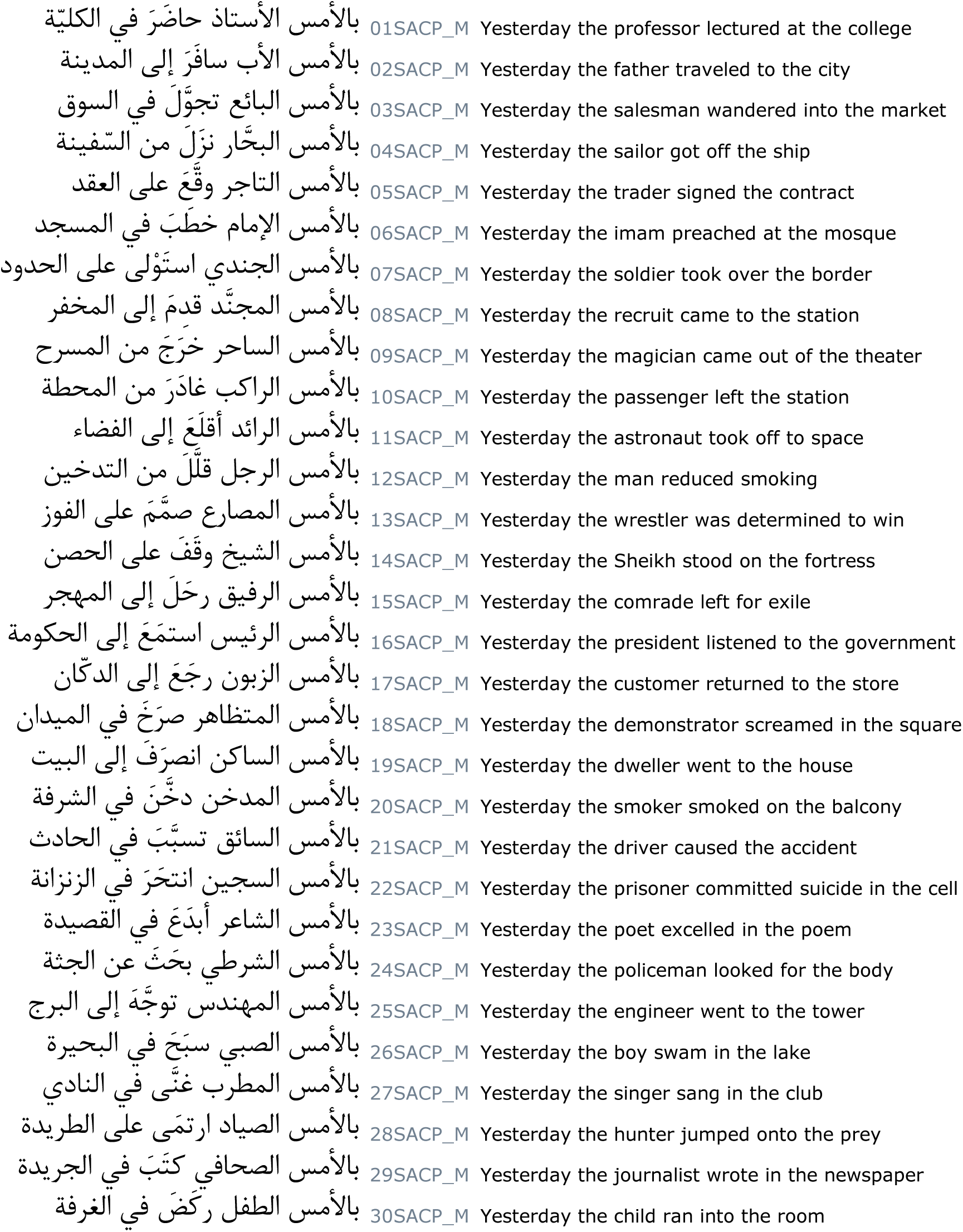
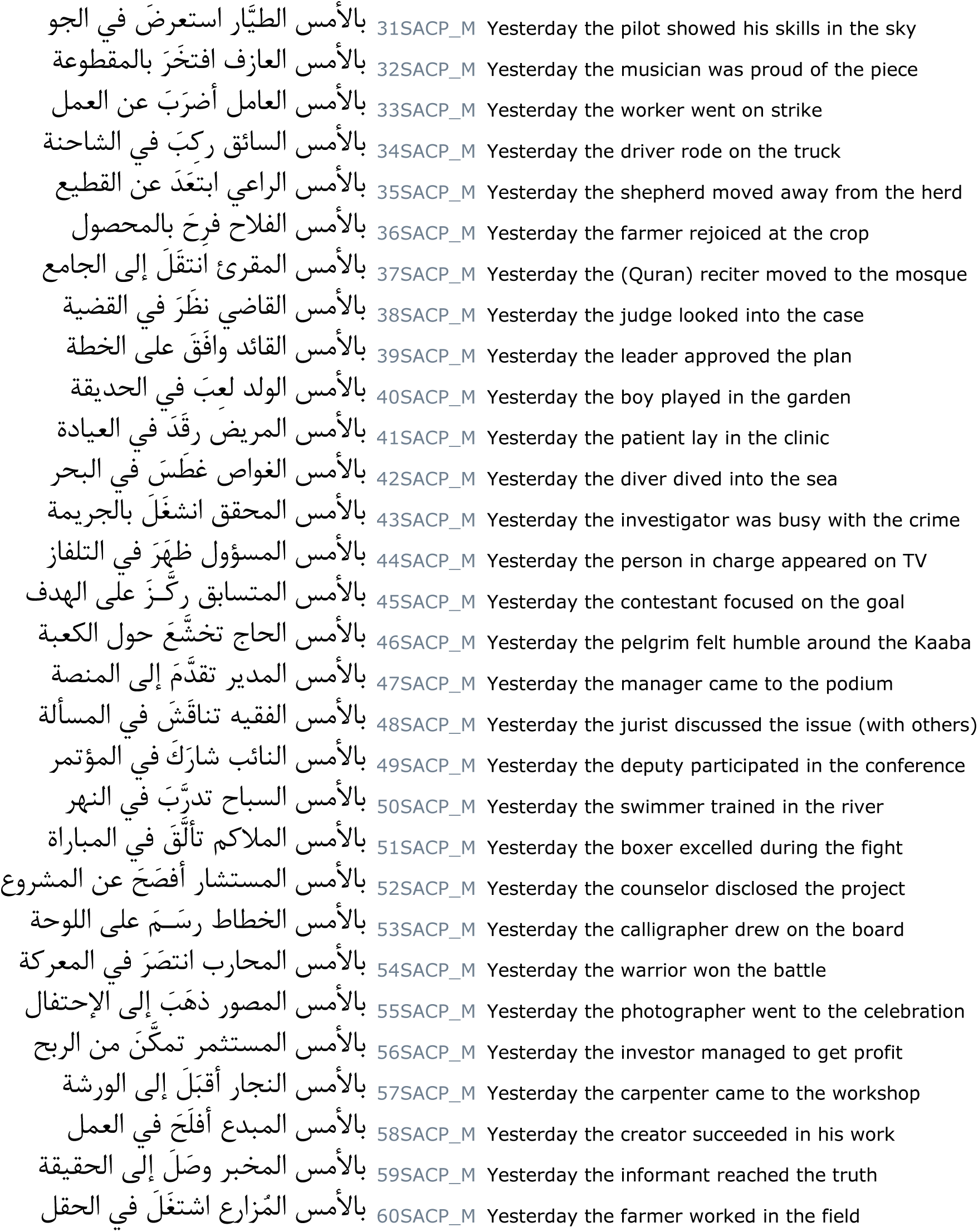
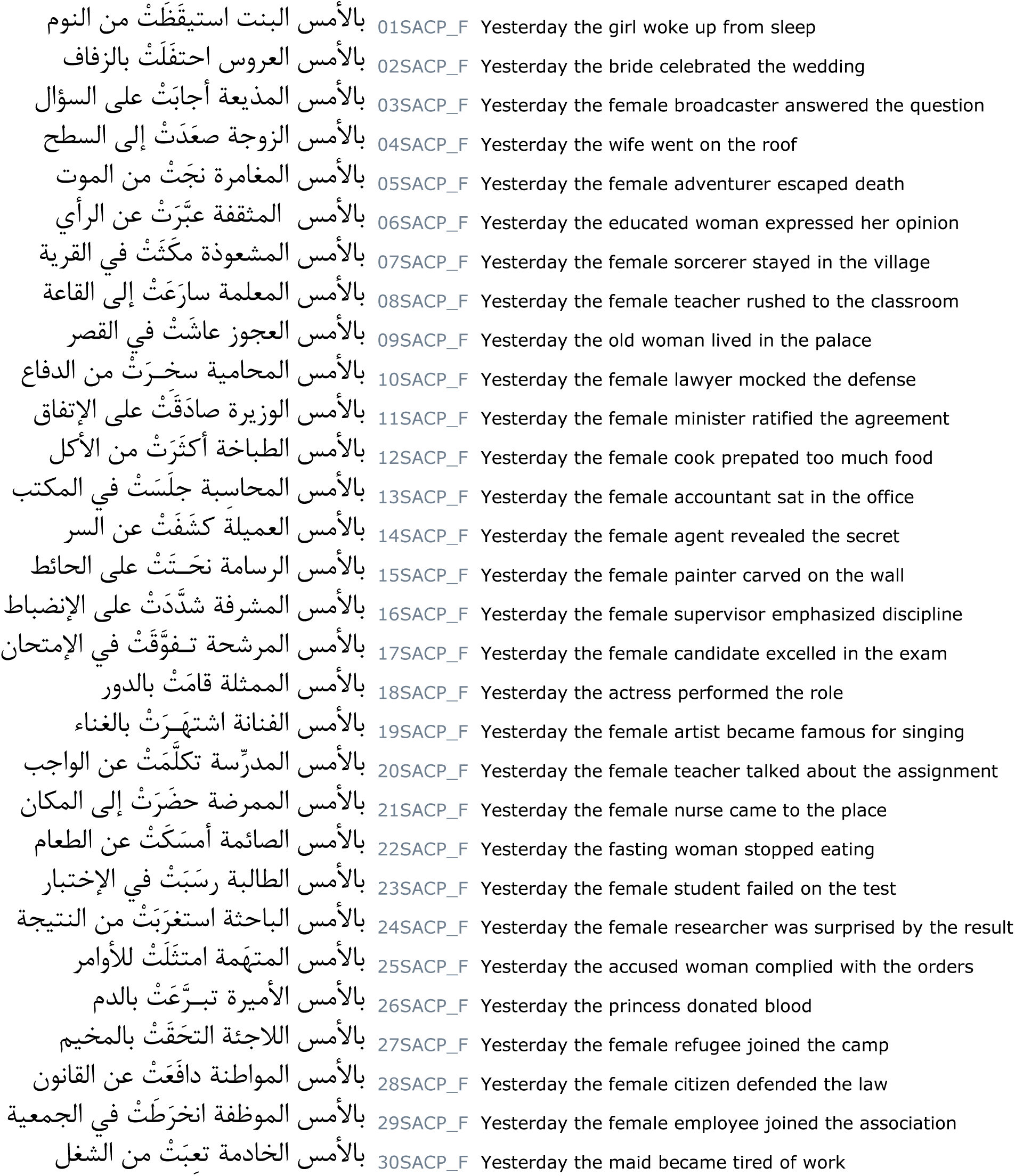
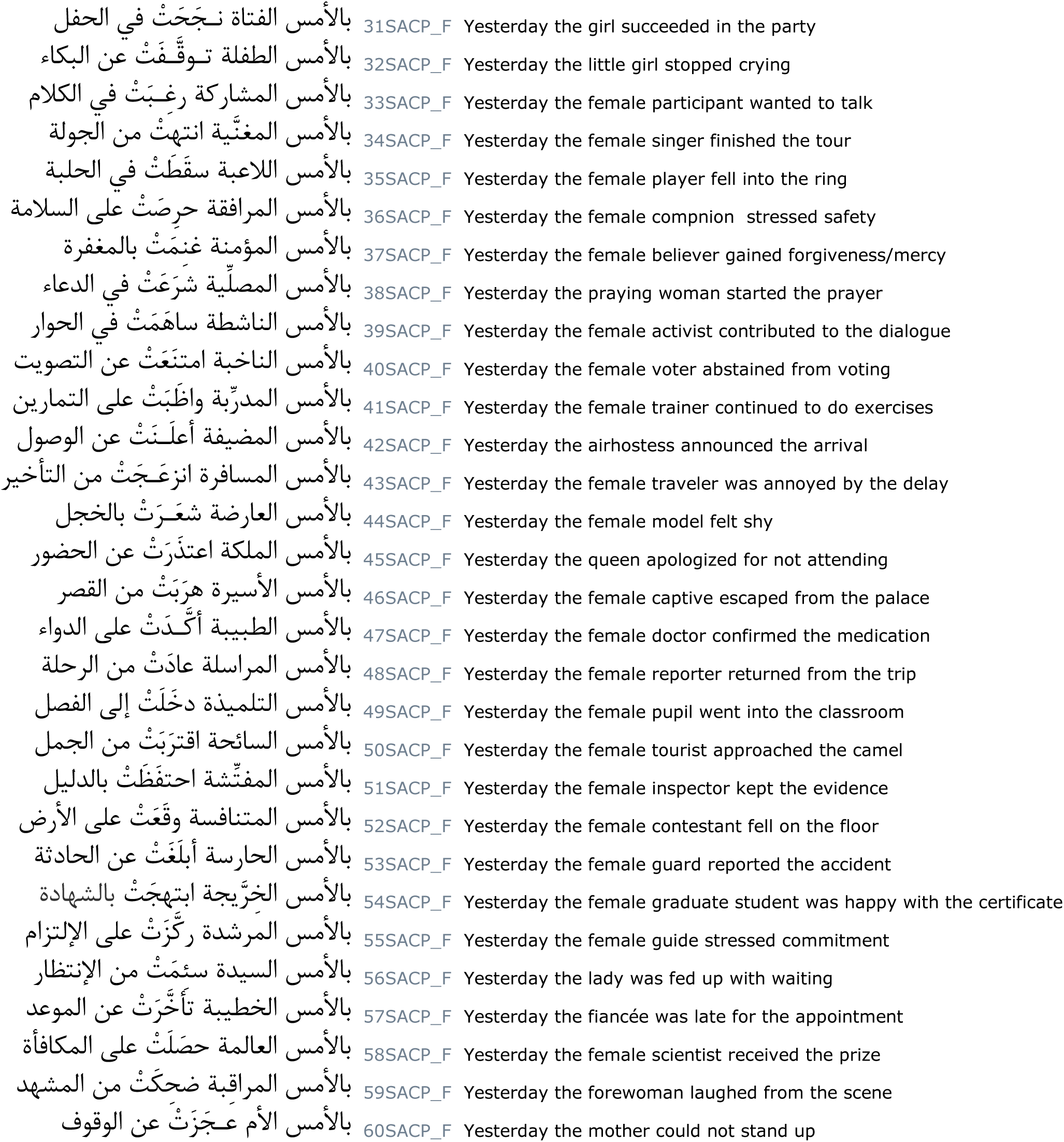
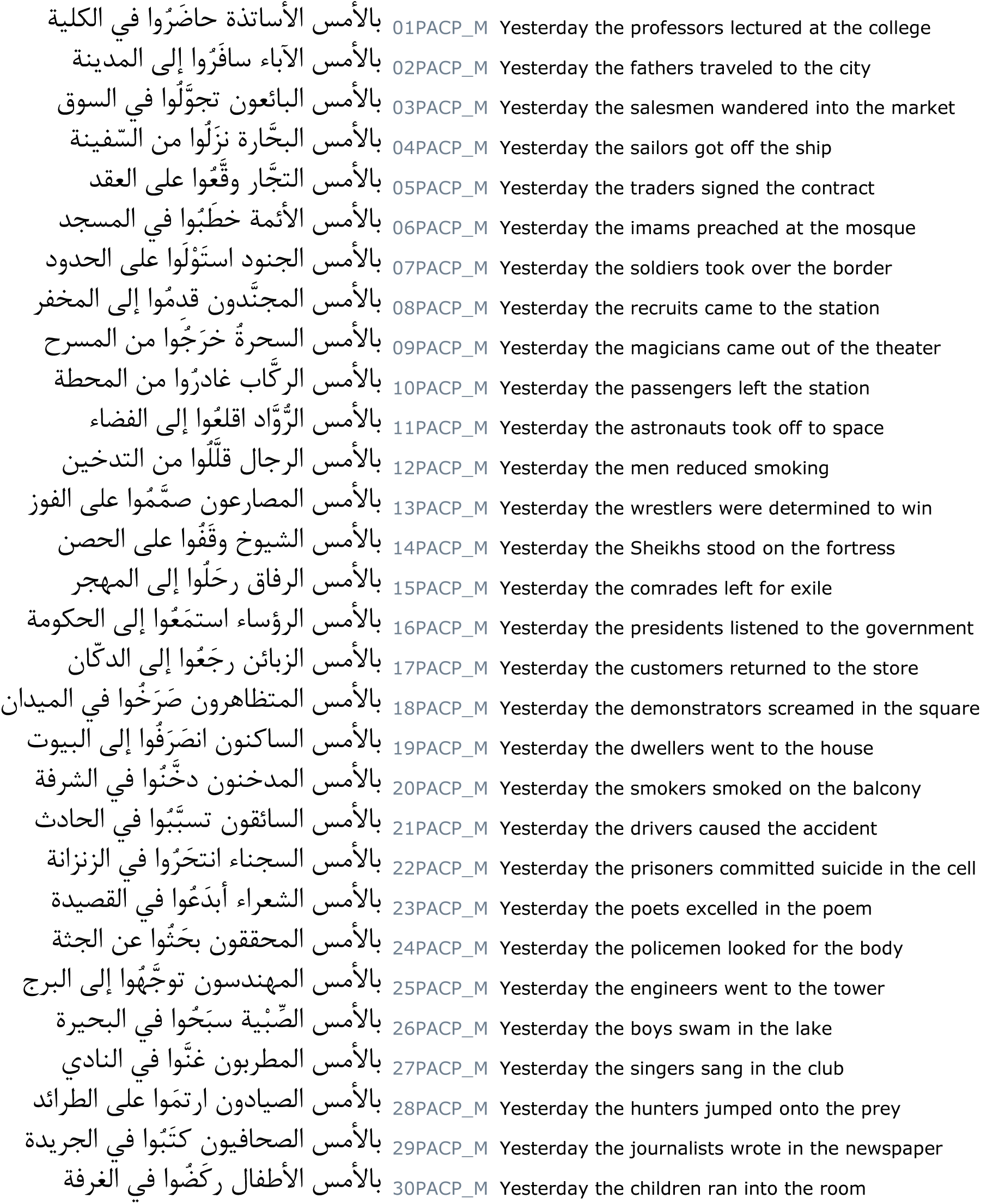
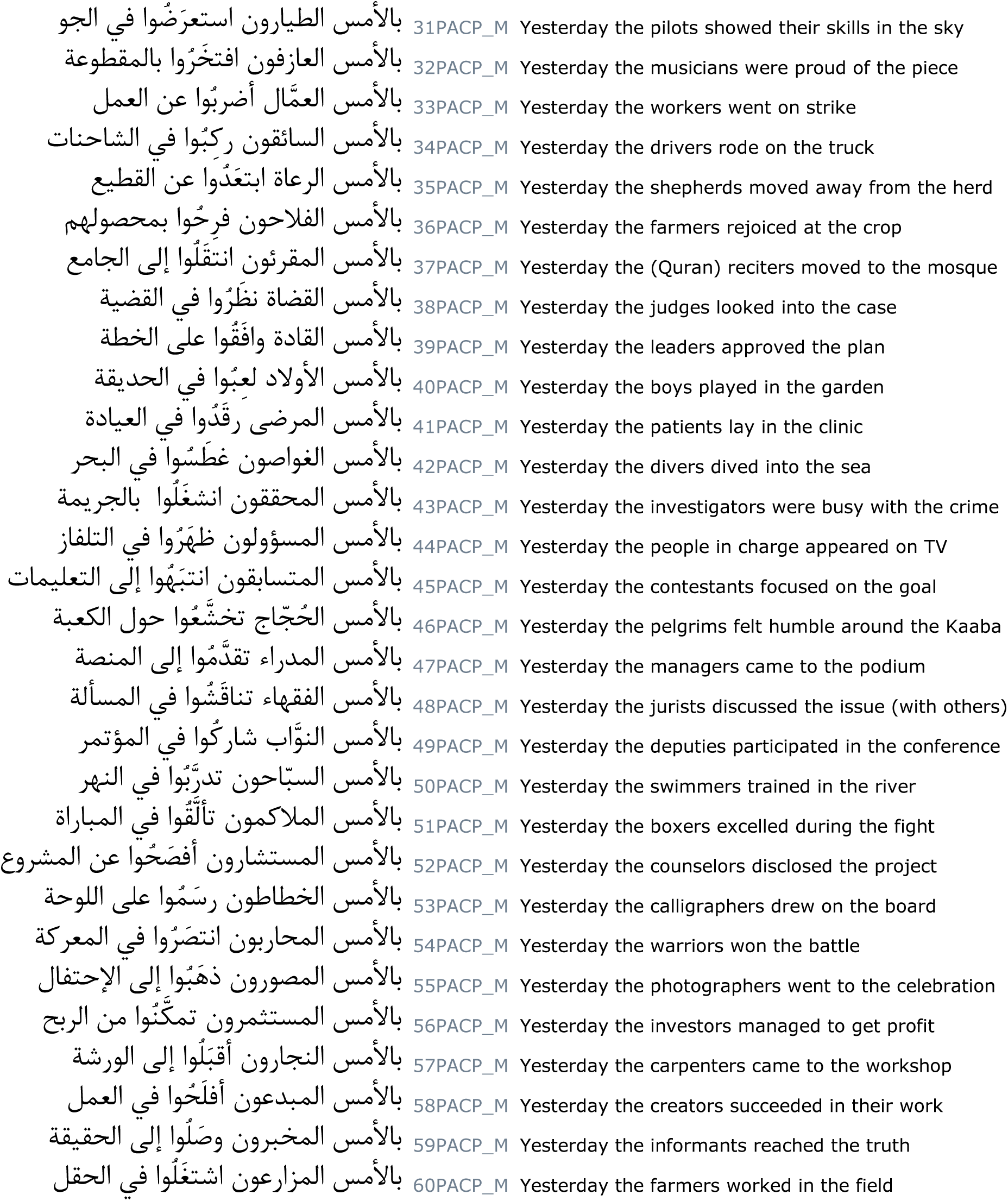
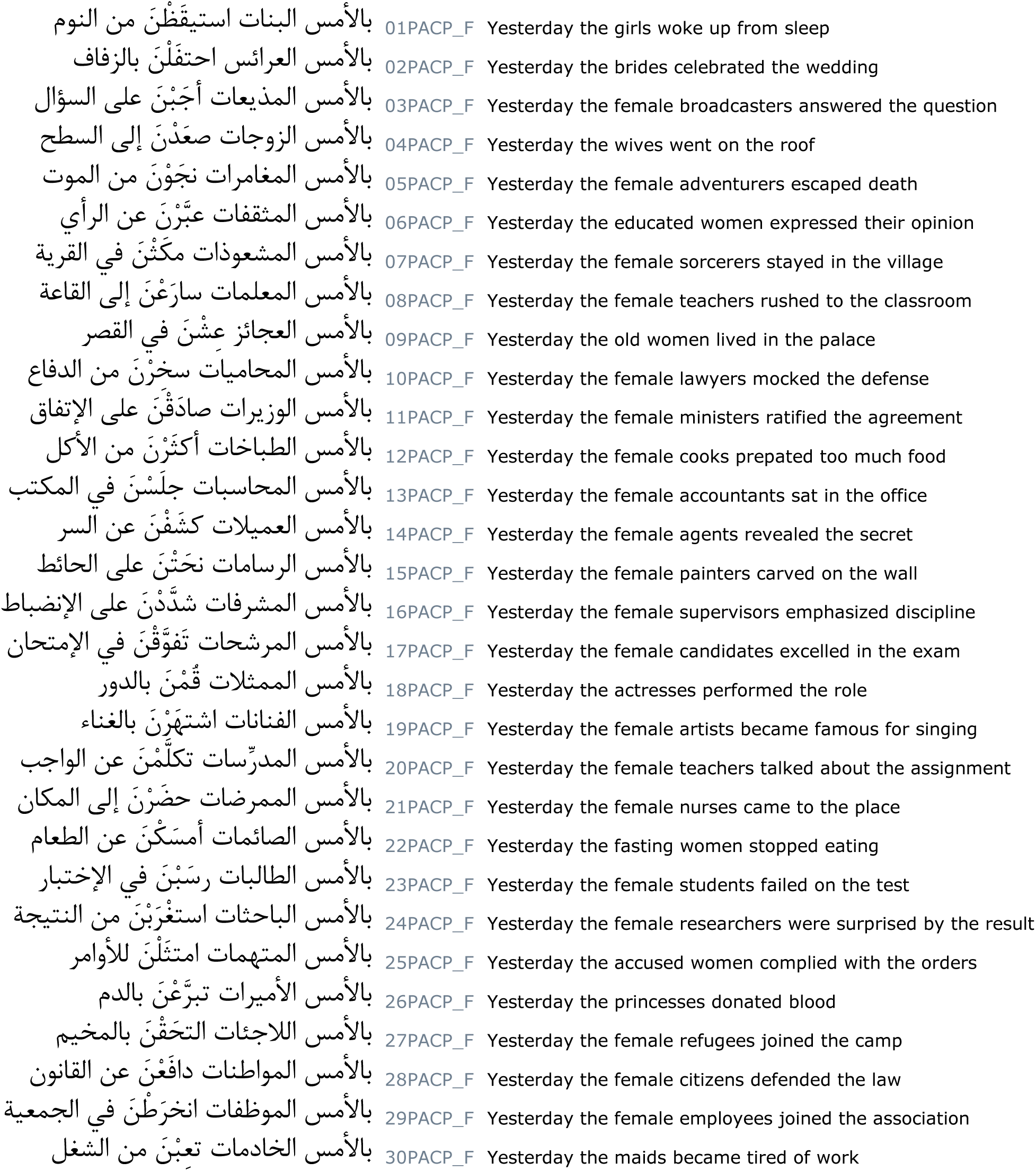
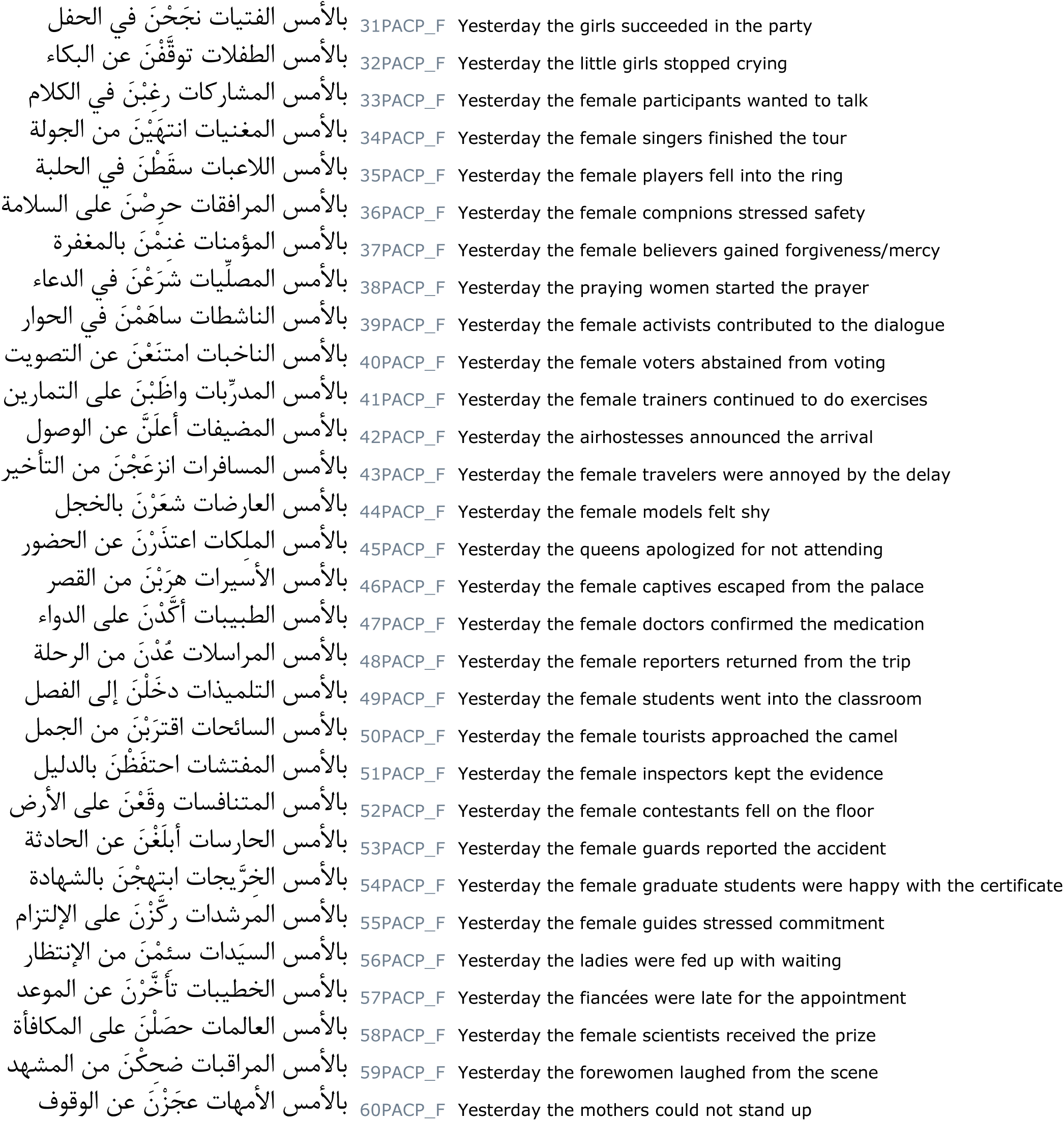

## Footnotes

1. Parts of the UAE University campus are gender-segretated, and the EEG laboratory of the Department of Linguistics was located on the male-only campus at the time the study was conducted, with strict entry restrictions for female undergraduate students. Furthermore, student/research assistants were not available for assistance with data collection, and the first author personally collected the data on his own. It was therefore very difficult to obtain access to the lab for female students to participate in our study. While it is unusual in ERP studies on language processing to recruit exclusively male / female participants, we believe that this need not be a cause for concern in interpreting our results. This is because, except in auditory ERP studies involving implicit voice-based emotional inferences (van den Brink et al., 2010) and the processing of emotional prosody (Schirmer, Kotz, & Friederici, 2002), there is no evidence we could find in the ERP literature on language processing in healthy adults that would suggest systematic gender-based ERP differences. Even the differences reported for implicit processing of emotional prosody disappear when explicit processing of prosodic information is required (Schirmer, Kotz, & Friederici, 2005; Schirmer et al., 2006). Indeed, in a critical review of several studies that investigated sex differences in language processing using a multitude of methods, Wallentin (2009) could find no evidence for a consistent difference between healthy adult female and male individuals, neither in their verbal abilities nor in their brain structure and function related to language processing, and concluded that‘sex should not be considered a large confounding factor in neuroimaging studies of language processing’ (Wallentin, 2009, p. 181).

2. Nevertheless, in order to verify that this is the case, we reanalysed the data post-hoc using two slightly broader bandpass filters offline, namely 0.1−30 Hz and 0.3−30 Hz. In the analysis using the 0.1−30 Hz bandpass filter, as an expected consequence of the slightly lower high-pass cutoff, slow signal drifts were apparent within the sentence epoch, which however could be rectified using a pre-critical baseline correction. In the analysis using the 0.3−30 Hz bandpass filter, due to the identical high-pass cutoff that we have employed in our analysis, no baseline correction was required. The overall pattern of ERPs from these analyses remained comparable to the pattern obtained in our analysis. An overview of this confirmatory analysis is provided in the supplementary supporting information.

3. As mentioned earlier, a confirmatory post-hoc analysis in an earlier time-window did not support an interpretation of some of the negativities in our study as instances of a LAN. Whilst the LAN versus N400 distinction is an important one (but see Bornkessel-Schlesewsky & Schlesewsky (2019) for a model that views all language-related negativities as a family of functionally related rather than distinct negativities), and our data does not rule out the possibility that some agreement violations may indeed lead to a LAN effect whereas others might engender an N400 effect, such a scenario of finding qualitatively different effects for agreement violations based on the feature violated and the language concerned would only strengthen the idea we are proposing here. See Discussion.

4. Since the focus in our study is on the differences in effects *within* a subject-type, it is not straightforward to compare our results with these studies. Nevertheless, restricting to subject-verb agreement and single feature violations, the P600 for the two subject-types in our results may be compared to at least two studies. The P600 difference in our results for the number feature between the two subject-types would be in line with a similar finding in Dutch that Kaan et al. (2000) reported. On the other hand, we found P600 effects for gender violations in both subject-types, whereas Deutsch & Bentin (2001) reported a P600 effect only when the subject was plural in Hebrew.

5. In their Basque study, Zawiswewski et al. (2016) found a significant difference in the 300-500 time-window, which they had initially interpreted as an N400 modulation for number violations as opposed to the other conditions; however, the statistics in their study as well as the topography of this effect with a posterior maximum in the P3 electrode region suggests that it is rather a P300 difference that they observe, as has been correctly pointed out by an anonymous reviewer of their manuscript. Their behavioural data also support this interpretation rather than their initial conjecture. Martinez de la Hidalga et al. (2019) report an N400 modulation in Basque, but only involving person and number features.

6. Nevertheless, we do not assume that these effects are strictly following each other in stages.

